# The SARS-CoV-2-like virus found in captive pangolins from Guangdong should be better sequenced

**DOI:** 10.1101/2020.05.07.077016

**Authors:** Alexandre Hassanin

## Abstract

Viruses closely related to SARS-CoV-2, which is the virus responsible of the Covid-19 pandemic, were sequenced in several Sunda pangolins (*Manis javanica*) seized in the Guangdong and Guangxi provinces of China between 2017 and 2019^1-3^. These viruses belong to two lineages: one from Guangdong (*GD/P*) and the other from Guangxi (*GX/P*). The *GD/P* viruses are particularly intriguing as the amino-acid sequence of the receptor binding domain of the spike protein is very similar to that of the human SARS-CoV-2 virus (97.4%)^2^. This characteristic suggests that *GD/P* viruses are capable of binding human ACE2 receptor and may therefore be able to mediate infection of human cells. Whereas all six *GX/P* genomes were deposited as annotated sequences in GenBank, none of the two *GD/P* genomes assembled in previous studies^2,3^ are currently available. To overcome this absence, I assembled these genomes from the Sequence Read Archive (SRA) data available for SARS-CoV-2-like viruses detected in five captive pangolins from Guangdong. I found the genome assemblies of *GD/P* virus of poor quality, having high levels of missing data. Additionally, unexpected reads in the Illumina sequencing data were identified. The *GD/P2S* dataset^2^ contains reads that are identical to SARS-CoV-2, suggesting either the coexistence of two SARS-CoV-2-like viruses in the same pangolin or contamination by the human virus. In the four other *GD/P* datasets^1^ many mitochondrial reads from pangolin were identified, as well as from three other species, namely, human, mouse and tiger. Importantly, I only identified three polymorphic nucleotide sites between the five *GD/P* sequences. Such low levels of polymorphism may reasonably be accounted for by sequencing errors alone, thus raising the possibility that the five pangolins seized in Guangdong in March 2019 were infected by the same virus strain, most probably during their captivity.

---

For each of the five *GD/P* samples sequenced on Illumina platforms (**Table 1**), I mapped the reads to the reference genome of the human SARS-CoV-2 virus (GenBank accession number: NC_045512)^4^ using Geneious Prime® 2020.0.3 and the “High sensitivity” option (maximum mismatch: 40%). Then, mapped reads were used for *de novo* assembly. All contigs were aligned to the SARS-CoV-2 genome and assembled into a consensus sequence used as reference to discover more reads in each *GD/P* dataset. All five *GD/P* genome assemblies (*GD/P2S, GD/P7L, GD/P8L, GD/P9L*, and *GD/P11L*) were of poor-quality having been previously sequenced at low depth (mean coverage between 0.2 and 6.5X) and therefore containing high levels (between 19% and 99%) of missing data (**Table 1**).

**Table 1.** Analyses of SRA data available for SARS-CoV-2-like viruses detected in captive pangolins from Guangdong.

| Illumina sequencing run |  |  | GD/P viral genome |  |  | Reads mapped to mitochondrial genomes |  |  |  |
| --- | --- | --- | --- | --- | --- | --- | --- | --- | --- |
| Code – Tissue | NCBI SRA | Reads | Reads | MC | MD | pangolin<br>NC_026781 | human<br>NC_012920 | tiger<br>NC_010642 | mouse<br>NC_005089 |
| GD/P2S<sup>2</sup> - Scale | SRR11093265 | 2,633 | 2,604 | 6.5X | 29% | 0 | 0 | 0 | 0 |
| GD/P7L<sup>1*</sup> - Lung 07 | SRR10168378 | 38,091,846 | 285 | 1.4X | 57% | 98,226 | 28 | 0 | 1,634 |
| GD/P8L<sup>1*</sup> - Lung 08 | SRR10168377 | 32,829,850 | 1,078 | 5.3X | 19% | 7,727 | 1,333 | 0 | 183 |
| GD/P9L<sup>1</sup> - Lung 09 | SRR10168376 | 36,135,230 | 36 | 0.2X | 88% | 13,770 | 47 | 3,447 | 0 |
| GD/P11L<sup>1</sup> - Lung 11 | SRR10168375 | 44,440,374 | 10 | 0.2X | 99% | 807,747 | 24 | 1,394 | 3 |

\*: SRA data used by Zhang *et al.*<sup>3</sup>. Abbreviations: MC: mean coverage; MD: missing data.

All 2633 reads sequenced for *GD/P2S*^2^ were mapped on the SARS-CoV-2 genome, indicating that all non-viral reads were removed. Curiously, within this pangolin dataset, I found 11 reads identical to the human SARS-CoV-2 genome (numbered 62, 412, 514, 786, 787, 1417, 1440, 1498, 2222, 2231, and 2403) and four reads very similar to SARS-CoV-2 (only a single mutation in reads 102, 502, 1390, and 1882) whereas several homologous reads (between 3 and 35) were found more divergent (between 2 and 9 mutations) but identical to other *GD/P* sequences. Two hypotheses can be proposed to explain this result: (1) the *GD/P2S* sample contained two different SARS-CoV-2-like viruses; or (2) the sample had been contaminated by the human SARS-CoV-2 virus. To choose between these two hypotheses the full raw dataset for *GD/P2S* is required. All Illumina reads generated for *GD/P2S* should consequently be deposited by the authors of the study^2^ in NCBI without any filtration process.

Less filtered SRA data were provided for other pangolin samples, i.e., *GD/P7L, GD/P8L, GD/P9L*, and *GD/P11L*^1^. It was therefore possible to extract mitochondrial sequences from the host (the Sunda pangolin) in order to determine the geographic origin of the seized animals. As shown in **Table 1**, many mitochondrial reads of pangolin (≥ 7727) were found for these four *GD/P* samples. Pairwise distances between assembled mitochondrial genomes were between 0.12% - 0.41%, confirming the four samples were collected on different pangolins, and suggesting they came from different localities in Southeast Asia. It should be noted mitochondrial reads could not be analysed for *GD/P2S* because the authors^2^ have removed all non-viral reads. It is therefore impossible to prove that the pangolin (*GD/P2S*) analysed by Lam *et al*.^2^ differs from those studied by Liu *et al*.^1^.

Surprisingly, numerous mitochondrial reads from other species (nucleotide identity = 100%) were also detected in the sequencing data. The *GD/P7L* dataset contains 1634 reads of mouse (*Mus musculus*), representing 2% of pangolin reads. The *GD/P8L* dataset contains 183 mouse reads (2%), and 1333 human reads (17%) (M7b haplogroup, which is found in humans from China and Southeast Asia). The *GD/P9L* dataset includes 3447 reads of tiger, subspecies *Panthera tigris altaica* (25%) and the *GD/P11L* dataset includes 1394 tiger reads (<1%). This unexpected range of mammalian species that I have identified clearly warrants an explanation. The most likely hypotheses are that laboratory experiments were contaminated by RNA molecules from multiple organisms, or that different RNA extractions were pooled into the same library.

When the five partial *GD/P* genomes were compared to each other, only three nucleotide sites were found to be polymorphic: (1) position 1807: A in 21 *GD/P2S* reads *versus* C in four *GD/P8L* reads; (2) position 5228: A in two *GD/P9L* reads, one *GD/P7L* read, and one *GD/P8L* read *versus* C in three *GD/P2S* reads; (3) position 24979: G in 15 *GD/P2S* reads and one *GD/P8L* read *versus* A in three *GD/P7L* reads. Considering the low-coverage of *GD/P* genomes, these differences can be interpreted as sequencing errors. I suggest therefore that all five pangolins were infected by the same *GD/P* virus strain, approximately at the same time, and most probably during their captivity^5^. I decided therefore to assemble a consensus *GD/P* genome by pooling together all reads sequenced from the five *GD/P* samples. The quality of the assembled genome is still very low with a mean coverage of 13X being composed of 26 fragments and containing 6.4 % missing data by comparison with the human SARS-CoV-2 genome. In particular, the sequence of the gene coding for the spike protein is composed of three fragments and includes 6.4 % missing data.

Reliable whole genome sequences of the virus detected in pangolins from Guangdong are crucial to better understand the origin of Covid-19. These sequences could be used in many studies, in particular to estimate mutation and recombination rates during the evolutionary history of viruses related to SARS-CoV-2. For this reason, I strongly encourage Chinese researchers to re-sequence the *GD/P* genome more deeply in order to reach a mean coverage of 30X, as often recommended for genomic studies^6^. In this spirit, I would encourage editorial boards of relevant journals to maintain their data publication standards by requiring authors to fully make available their unfiltered data for the benefit of scientific collaboration in tackling the current pandemic.

## Supporting information

The 15 GD/P2S reads showing high similarity with the SARS-CoV-2 human virus

GD/P consensus genome of SARS-CoV-2-like virus assembled from all SRA data generated from pangolins seized in the Guangdong province of China

Mitochondrial genomes assembled from SRA data published by Liu et al. (2019) (see Table 1 for details)

## Acknowledgements

I would like to thank Anne Ropiquet and Huw Jones for helpful comments on the first version of the manuscript.

## Notes

### Competing Interest Statement

The authors have declared no competing interest.

