## Supplementary material for "The SARS-CoV-2-like virus found in captive pangolins from Guangdong should be better sequenced": The 15 GD/P2S reads showing high similarity with the SARS-CoV-2 human virus: Supplementary_information_1.pdf

#### Supplementary data

#### <u>The 15 *GD/P2S* reads showing high similarity with the SARS-CoV-2 human virus</u>

The reads were aligned to SARS-CoV-2 (NC_045512), which was used as reference to show nucleotide variations. The GD/P consensus genome reconstructed from all SRA data generated from pangolins seized in the Guangdong province of China[1,2] (Table 1) was also aligned to SARS-CoV-2 for comparison.

##### <u>Read 62</u>

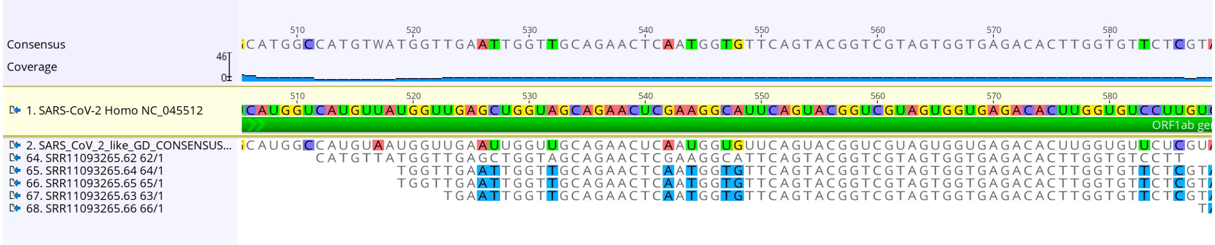


##### <u>Read 102</u>

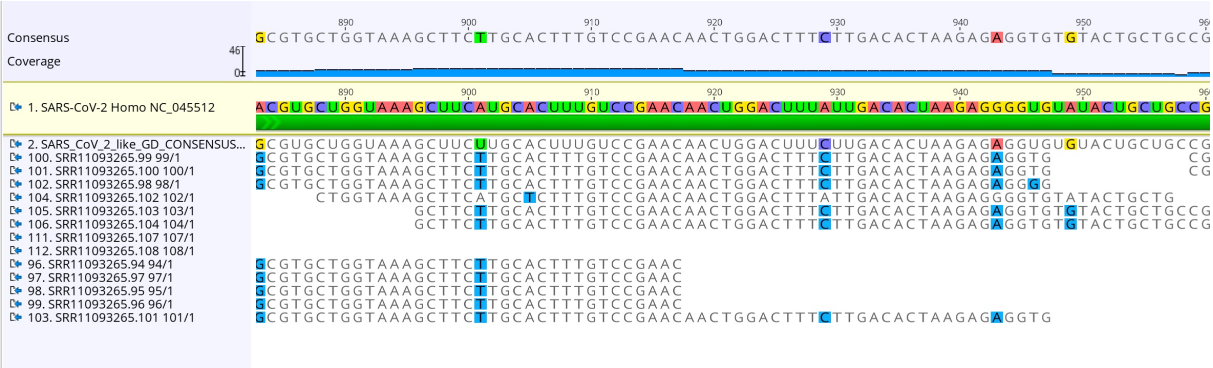


##### <u>Read 412</u>

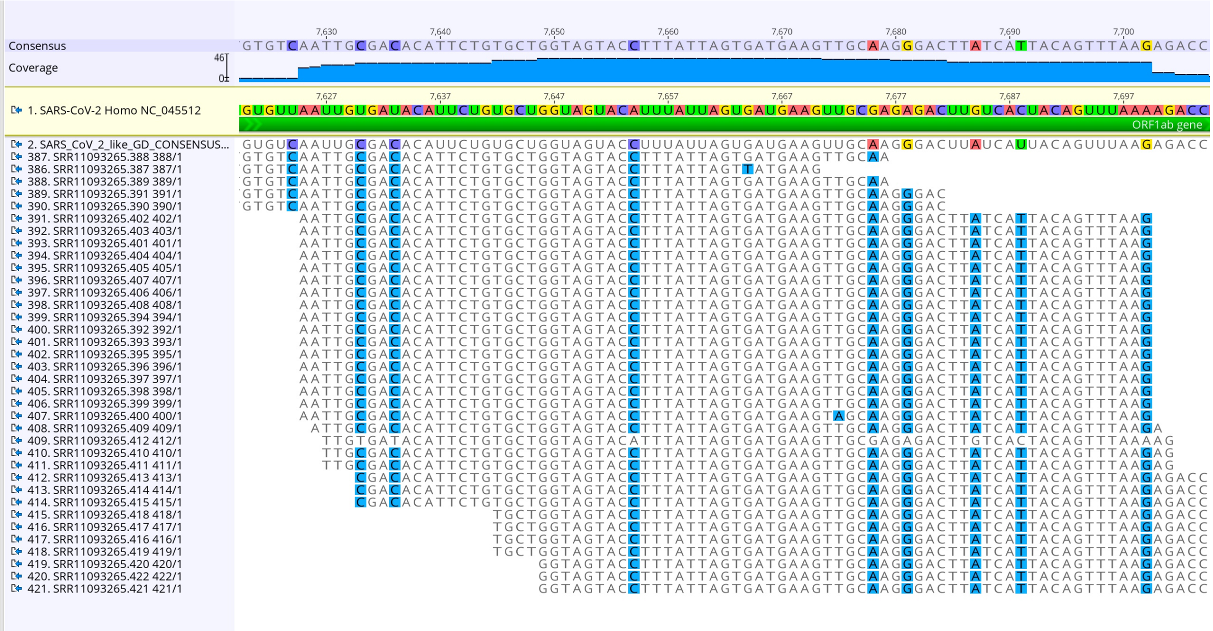

### Read 502

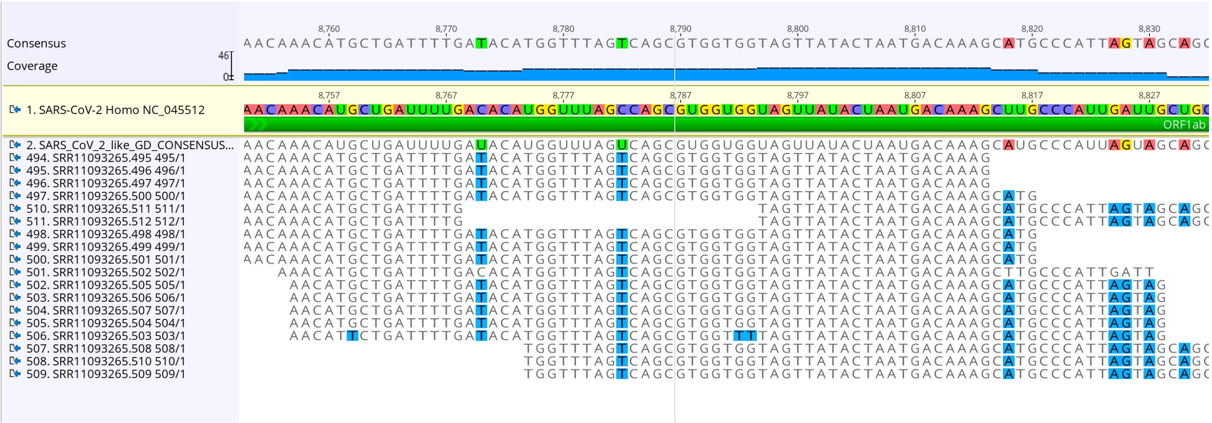


### Read 514

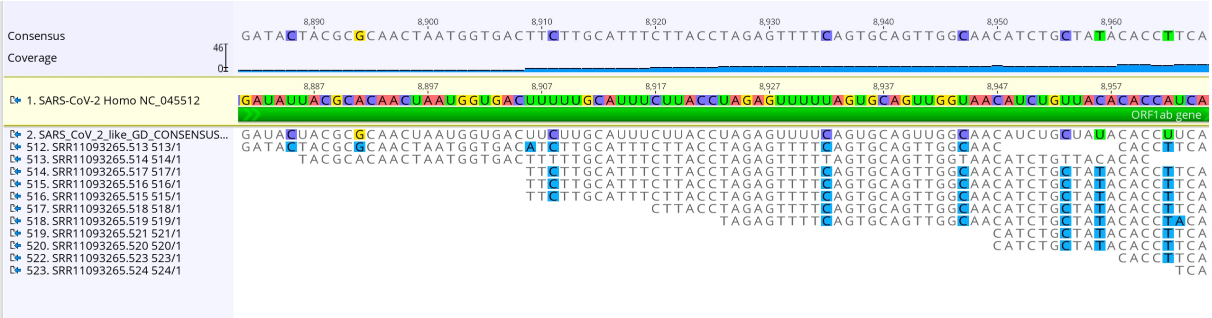


### Reads 786 and 787

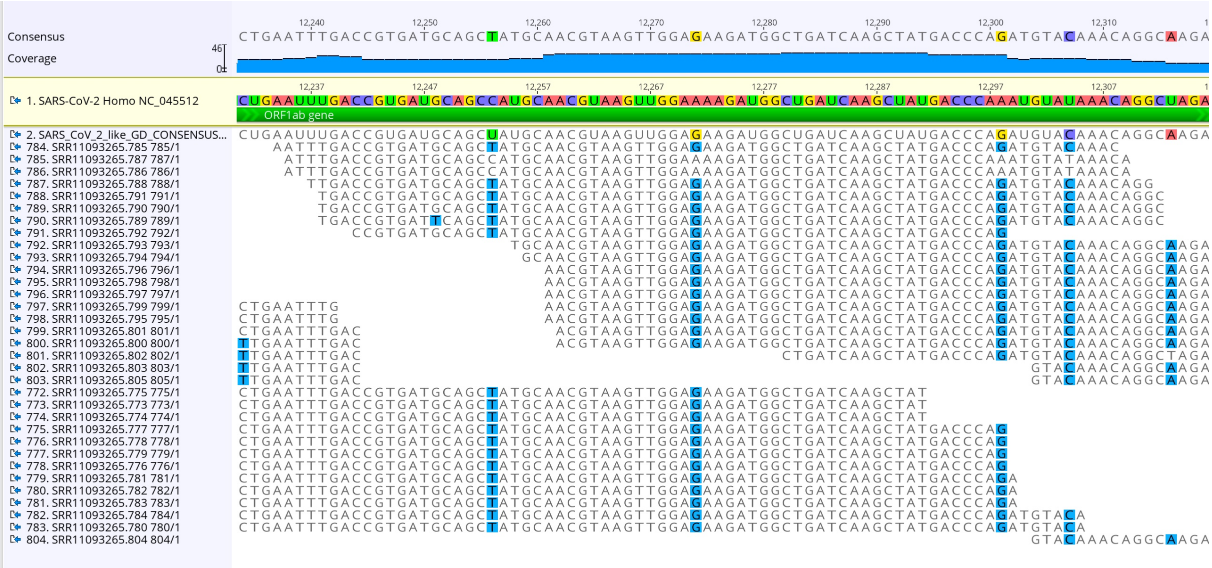


### Read 1390

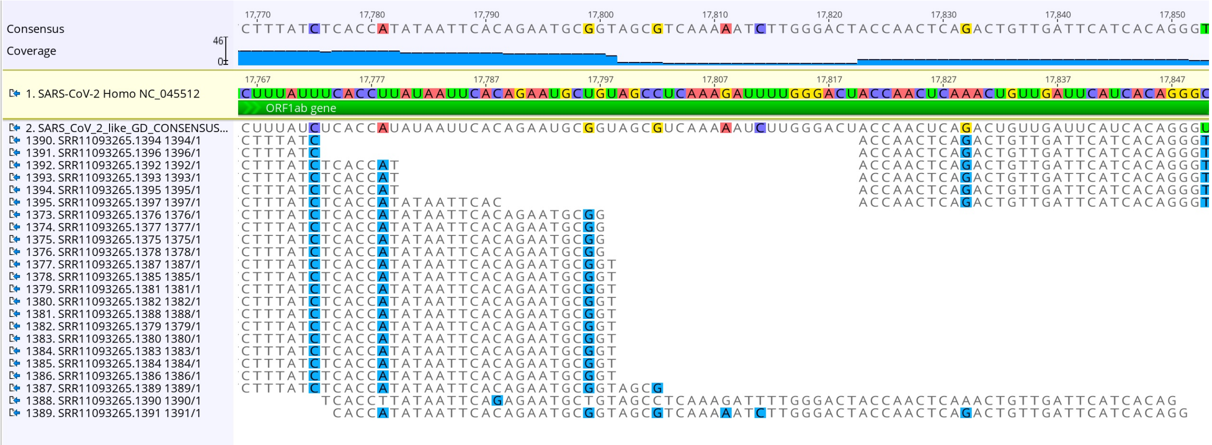

#### Read 1417

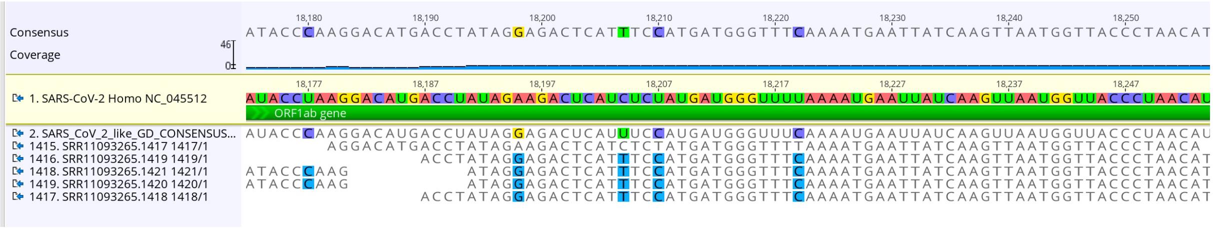


#### Read 1440

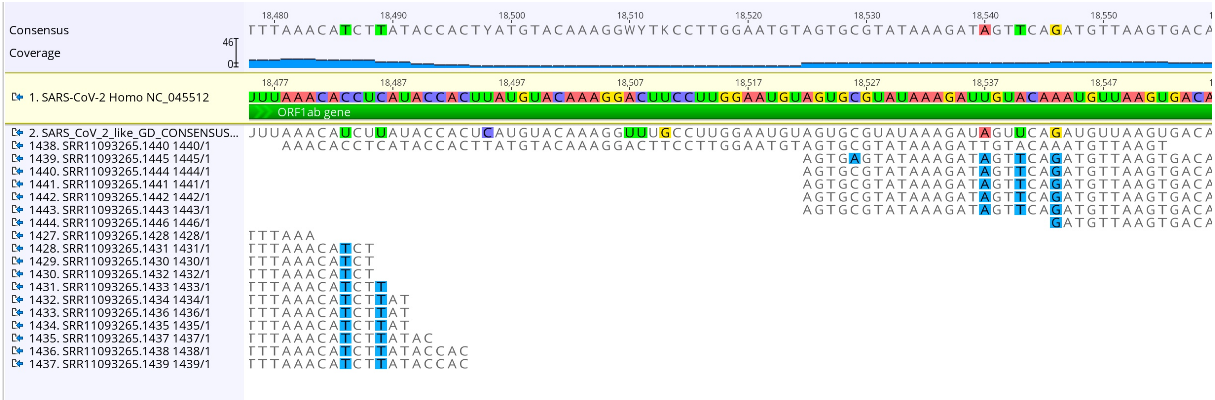


#### Read 1498

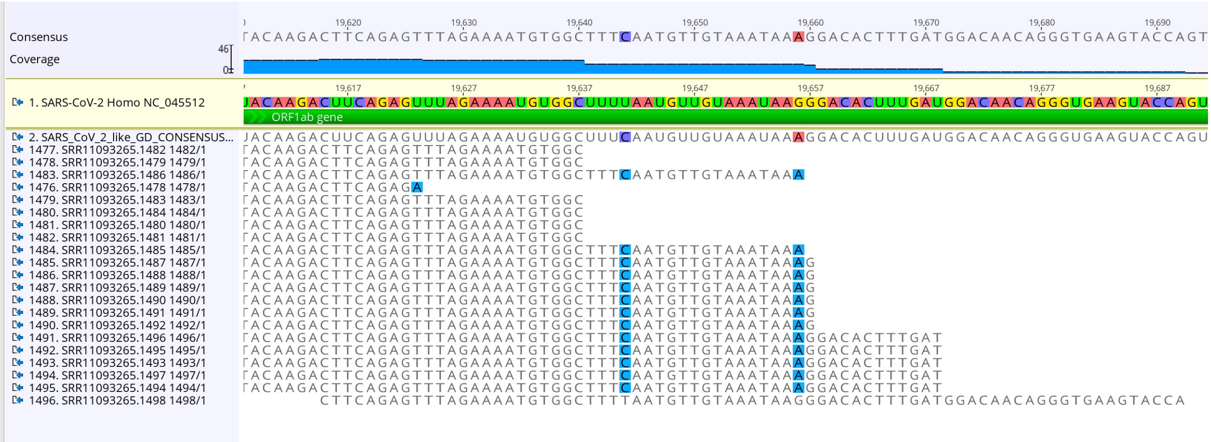


#### Read 1882

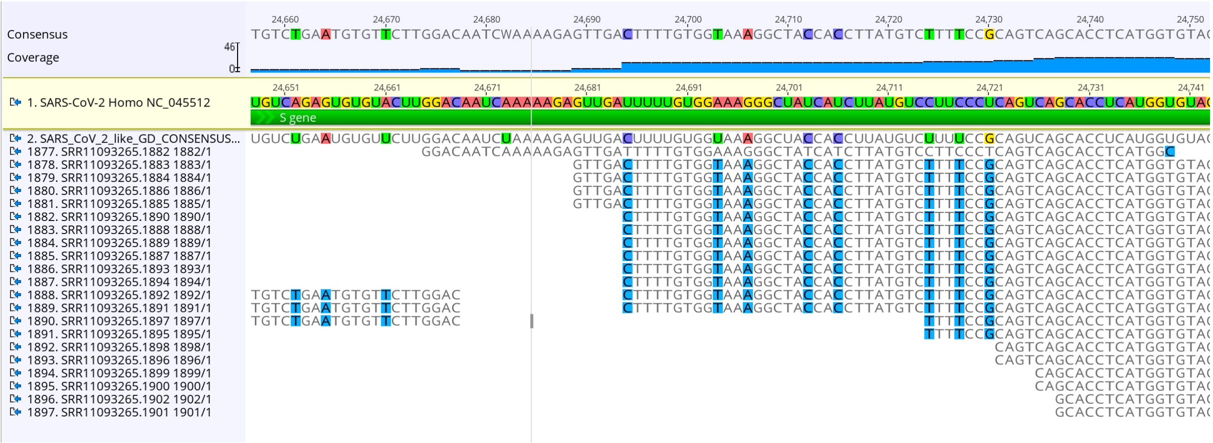

#### Read 2222

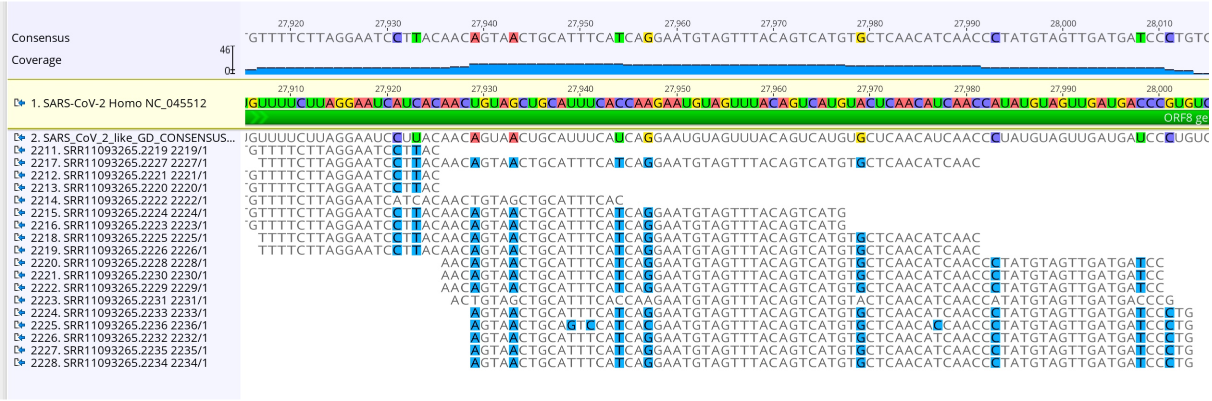


#### Read 2231

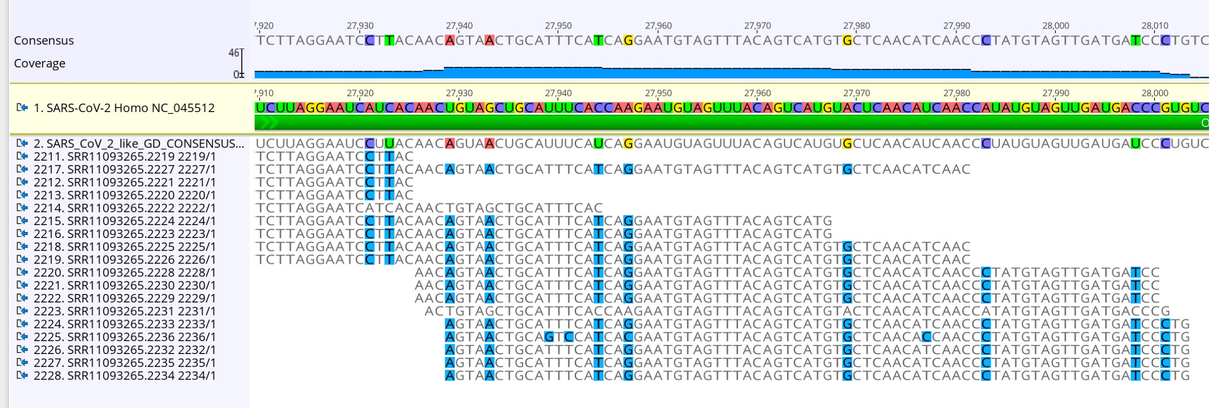


#### Read 2403

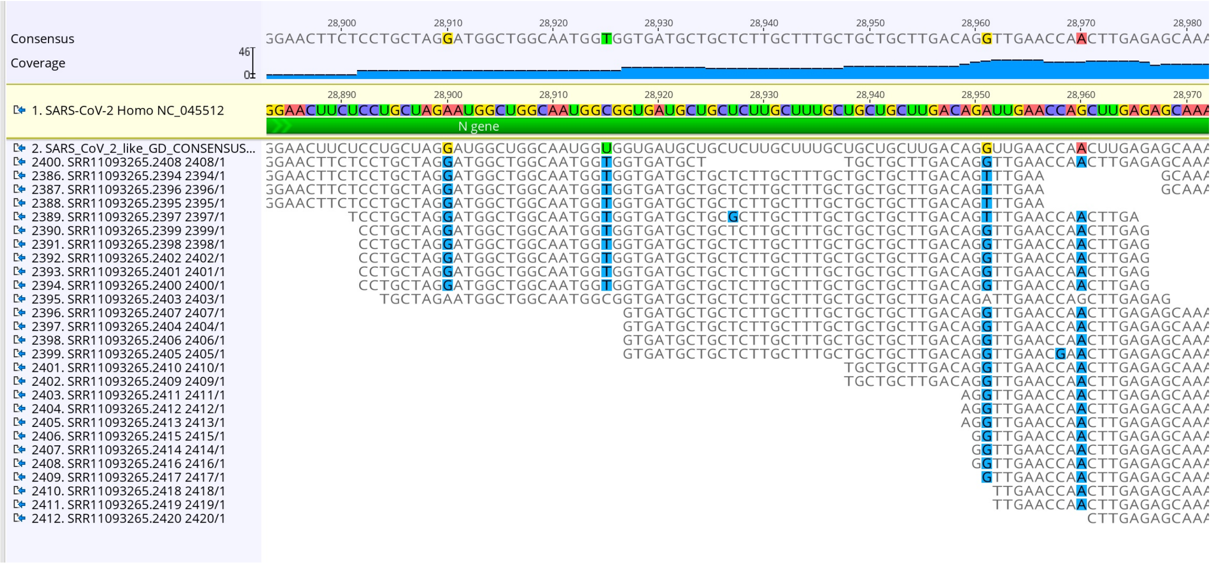
