## Supplementary material for "The SARS-CoV-2-like virus found in captive pangolins from Guangdong should be better sequenced": GD/P consensus genome of SARS-CoV-2-like virus assembled from all SRA data generated from pangolins seized in the Guangdong province of China: Supplementary_information_2.pdf

**GD/P consensus genome of SARS-CoV-2-like virus assembled from all SRA data generated from pangolins seized in the Guangdong province of China[1,2]** (see Table 1 for details)

>SARS_CoV_2_like_GD_CONSENSUS_26fragments [AH 20200501]
?????????????????????????????????????????????????????UUAAAAUC
UGUGUGGCUGUCACUCGGCUGCAUGCUUAGUGCACUCACGCAGUAUAAUUAAUAA
CUAAUUACUGUCGUUGACAGGACACGAGUAACUCGUCUAUCUUCUGCAGGCUGCU
UACGGUUUCGUCCGUGUUGCAGCCGAUCAUCAGCAUACCUAGGUUUCGUCCGGGU
GUGACCGAAAGGUAAGAUGGAGAGCCUUGUCCCUGGUUUCAACGAGAAAACACAC
GUCCAACUCAGUUUGCCUGUUUUACAGGUUCGCGACGUGCUCGUACGUGGCUUUG
GAGACUCCGUGGAGGAGGCUAUCUCAGAGGCACGUCAACAUCUCAAGGAUGGCAC
UUGUGGCUUAGUAGAGGUUGAAAAAGGCGUCUUGCCUCAACUUGAACAGCCCUAU
GUGUUCAUCAAACGUUCUGAUGCCCGAACUGCACCGCAUGGCCAUGUAAUGGUUG
AAUUGGUUGCAGAACUCAAUGGUGUUCAGUACGGUCGUAGUGGUGAGACACUUG
GUGUUCUCGUACCCCAUGUGGGUGAAACACCUGUUGCUUACCGCAAAGUUCUUCU
UCGCAAGAACGGUAAUAAAGGAGCUGGUGGUCACAGCUAUGGCGCCGAUCUAAAG
UCCUAUGACUUAGGUGACGAGCUGGGCACUGAUCCUUAUGAAGAUUAUCAAGAAA
ACUGGAACACUAAACAUGGCAGUGGUGUAACUCGUGAGCUCAUGCGUGAGCUUAA
UGGGGGCGCAUACACUCGCUAUGUCGAUAACAACUACUGUGGCCCUGAUGGCUAC
CCUCUUGAGUGCAUUAAAGACUUGCUGGCGCGUGCUGGUAAAGCUUCUUGCACUU
UGUCCGAACAACUGGACUUUCUUGACACUAAGAGAGGUGUGUACUGCUGCCGUGA
GCAUGACCAUGAAAUUGCAUGGUACACGGAACGCUCUGAUAAGAGUUAUGAAUU
GCAGACACCUUUUGAAAUUAAACUGGCAAAGAAAUUUGACACUUUUACUGGGGA
GUGCCCAAAUUUUGUAUUCCCUCUUAAUUCAACUAUCAAGACUAUUCAACCUAGA
GUUGAAAGGAAAAAGCUUGAUGGCUUUAUGGGUAGGAUUCGAUCUGUCUACCCU
GUUGCUUCACCUAAUGAAUGCAACCAAAUGUGCCUGUCAACUCUCAUGAAGUGUA
ACCAUUGUGGUGAAACUUCAUGGCAGACAGGCGAUUUUGUUAGAGCCACUUGUGA
GUUCUGUGGUACUGAAAAUUUGACUAAAGAUGGUGCUACAACUUGUGGUUACCU
UCCUCAAAAUGCUGUUGUCAAAAUUUACUGUCCAGCAUGUCAUAACCCAGAAAUA
GGACCUGAGCAUAGUCUUGCUGAAUACCACAAUGAAUCUGGUCUAAAGACCACUC
UUCGUAAGGGUGGUCGUACCAUUUCAUUUGGUGGUUGUGUCUUCUCCUAUGUUG
GCUGUCACAACAAAUGUGCCUAUUGG?????????????????ACAUAGGUUGCAAUCAC
ACAGGAGUUGUUGGAGAAGGUGCUGAAAGUCUGAAUGACAACCUUCUUGAAAUA
CUUCAGAAAGAUAAAGUCAAUAUCAACAUUGUUGGUGACUUUAAACUUAAUGAA
GAGAUUGCCAUUAUAUUGGCAUCUUUCUCUGCUUCUACAAGUGCUUUUGUAGAAA
CUGUCAAAGGUUUGGAUUAUAAAACAUUCAAACAGAUUGUUGAAUCCUGUGGUA
ACUUUAAAGUUACCAAAGGMAAAGCUAAGAAGGGUGCUUGGAAUAUUGGUGAGC
AAAAAUCAAUACUGAGUCCUUUGUAUGCAUUUGCAUCUGAAGCUGCUCGUGUUAU
ACGCGCCAUUUUCUCUCGUACUCUUGAAACUGCUCAACACUCAGUUCGUGUCCUA
CAACAGGCCGCUAUAACUAUUCUUGAUGGAAUUUCACAGUAUUCACUGAGACUCA
UUGAUGCUAUGAUGUUCACAUCUGAUUUAGUUACUGACAACCUUGUUGUGAUGG
CAUAUAUCACAGGUGGUGUUGUCCAAAUGACAUCACAGUGGCUUACAAAUAUCUU
UGGCACUGUUUAUGAAAAACUUAAACCUAUUCUUGAAUGGCUUGAAGA????????????
?????????????????GAGACGGUUGGGAAAUUGUUAAAUUUAUCUCAACCUGUGCUUG
UGAAAUUGUCGGUGGACAAAUUGUCACCUGUGCAAAGGAAAUUAAAGAAAGUGU
CCAAACAUUCUUUAAGCUUGUAAACAAAUUCCUAGCUUUGUGUGCUGACUCUAUC
AUUAUUGGUGGGGCUAAACUCAAGGCUGUGAAUUUGGGUGAAACAUUU????????????
????????????????????????????????????????????????????????????
????????????????????????????????????????????????????????????
UGAAGCUAUUGAUGCUCCACUAGUUGGUACACCAGUUUGUAUUAACGGGCUUAUG

UUGCUCGAAAUUAAAGAUACAGAGAAGUAUUGUGCUCUUGCACCUAACAUGAUG
GUUACAAAUAACACCUUCACACUUAAAGGUGGUGCACCAACAAAAGUUACUUUUG
GUGAUGACACCGUAAUUGAAGUACAGGGUUACAAAAGUGUAAGUAUCACUUUUG
AACUCGAUGAGAGAGUAGAUAAAGUACUUAAUGAAAAGUGCUCUAACUACACAG
UUGAA????????AGAAGUAAAUGAGUUUGCUUGUGUUGUCGCUGAUGCUGUUAUAA
AGACUUUACAGCCAGUAUCUGAAUUACUUACACCACUAGGCAUAGAUUUAGACGA
GUGGGGUAUGGCAACAUACUACUUGUUUGAUGAAUCUGGUGAGUUUAAUUUAGC
CUCUCAUAUGUACUGUUCUUUCUAUCCUCCAGAUGAGGAUUAUGAAGAAGAUGAG
UGUGAAGAGGAACAGUAUGAACCAUCAACUCAGUAUGAGUAUGGUACAGAGGAU
GAUUACCAAGGUAAAUCUUUGGAAUUUGGUUCAACCUCUUCUGCUUCUCAAAUAG
AAGAAGAACCAGAGGAAGAUUGGUUAGAAGAUGGCAAUGAGGAAAUUGCUAUGC
AAGAAGA?????????????????????????????????????????????????????????????????????????????
???????????????????CAGUGAACAAUUUUACUGGUUAUUUAAAACUCACUAACAAUGU
UUUCAUUAAAAAUGCAGACAUUGUGGAAGAAGCUAAACAGGUGAAACCAACAGU
AGUUGUUAAUGCAGCUAAUGUUUACCUUAAACAUGGAGGAGGUGUUGCUGGAGC
UUUAAACAAAGCUACAAACAACGCUAUGCAAGUAGAAUCAGACAAUUACAUAGCC
AC?????????????????????????????????????????????????????????????????????????????????????
????????????????????????????????????????????????????????????????????????????????????????
????????????????????????????????????????????????????????????????????????????????????????
????????????????????????????????????????????????????????????????????????????????????????
?????????????????????GAAAAUAAACCUUCAGUUGAACAAAAGCAACAGGCUGAAGA
AAAGAAAGUCAAGGCUUGUGUUGAAGAAGUUACAACCACUUUGGAAGAAACUAA
GUUUCUCACAGAGAAUUUGUUACUUUAUAUAGACAUUAAUGGCAACCUUCAUCCA
GAUUCUGCCAUGCUUGUUAAAGACACUGACACCACCUUUCUAAAGAAAGAUGCUC
CUUACAUAGUAGGUGAUGUCAUUAAGGAAGGUGUUUUAACUGCUGUAGUUAUAC
CUACUAAAAAGCUGGUGG??????????????????????????????????????????AGACAAUUA
UAUAACCACCUACCCUGGACAGGGUAAUAAUGGUUAUACAUUAGAAGAGGCAAAG
ACAGUGCUUAAAAAGUGCAAGAGCGCUUUUUACAUACUACCAUCCAUUGUCUCUA
AUGAGAAAGAAGAAAUUCUUGGAACUGUUUCUUGGAAUUUGCGAGAAAUGCUUG
CACAUGCUGAGGAAACACGUAAAUUAAUGCCUGUUUGUAUGGAAACUAAAGCUA
UAGUUUCAACCAUACAACGUAAAUACAAGGGUAUUAAAAUACAGGAAGGUGUGG
UUGAUUAUGGUGCUAGGUUUUACUUCUAUACUAGUAAAACAACAGUAGCAUCAC
UCAUCAACACACUUAACAAUCUAAAUGAGACUCUCGUUACAAUGCCUUUGGGUUA
UGUAACACAUGGCCUUAACCUAGAAGAAGCUGCGCGGUACAUGAGAUCUCUUAAG
GUACCAGCGACAGUUUCUGUUUCUUCACCAGAUGCUGUAACAGCAUAUAAUGGUU
AUCUUACUUCUUCUUCAAAAACACCUGAAGAACAUUUCAUAGAAACCAUCUCUCU
AGCUGGUUCUUACAAAGAUUGGUCCUAUUCUGGACAAUCAACACAACUUGGUAUA
GAAUUUCUUAAAAGAGGUGAUAAAACUGUUUAUUACACCAUUAACCCUAUUGCU
UUUCAUACAGAGGGUCAAAUUAUCACCUUUGAUGAUUUAAAAACACUUCUCUCAU
UGAGAGAGGUUAGAACAAUCAAAGUGUUUACAACUGUAGACAAUGUUAAUCUCC
AUACACAAGUUGUGGACAUGUCUAUGACUUAUGGACAACAGUUUGGUCCCACUUA
UUUGGAUGGAGCUGAUGUCACUAAAAUAAAACCUCAUAAUUCACAUGAAGGUAA
GACAUUUUAUGUCUUGCCUAAUGAUGACACAUUACGUGCAGAGGCUUUUGAGUAC
UACCAUACAACAGAUGCUAGUUUUCUUGGUAGGUACAUGUCAGCUUUAAACCAUA
CCMAGAAGUGGAAAUACCCACAGGUUAAUGGUUUAACUUCUAUUAAGUGGGCAG
AUAACAAUUGUUACCUUGCUACUGCUUUAUUAGCACUCCAACAAAUAGAGUUGAA
GUUUAAUCCACCUGCUUUGCAAGAUGCUUAUUAUAGAGCUAGAGCCGGUGAUGCU
GCUAAUUUCUGUGCACUUAUCUUAGCCUAUUGUAACAAAACAGUAGGAGAGUUA
GGUGAUGUUAGAGAAACAAUGAAUUAUUUGUUUCAACAUGCCAAUUUGGAUUCU
UGUAAGAGAGUUCUAAAUGUGGUGUGUAAAACUUGUGGACAGCAACAAACCACU
CUUAAGGGUGUGGAAGCUGUUAUGUACAUGGGUACACUUUCUUAUGAACAACUU
AAGAAAGGUGUCCAAGUACCUUGUGUGUGCGGUAAACAAGCCACACAAUAUUUAG

UCCAACAAGAGUCACCUUUUGUUAUGAUGUCUGCACCACCUGCCGACUAUGAAUU
GAAGCAUGGUACCUUUGUUUGUGCUAGUGAAUACACUGGUAAUUACCAAUGUGG
CCACUACAAACACAUAACUUCUAAGGAAACUCUGUAUUGCAUAGAUGGUGCUUUA
CUGACAAAGUCUUCUGAAUACAAAGGUCCUAUUACAGAUGUUUUCUAUAAAGAA
AACAGUUACACAACAACCAUUAAACCAGUUACAUAUAAACUGGAUGGUGUUGUUU
GUACAGAAAUUGAUCCUAAAUUGGACAAUUACUAUAAAAAAGACAAUGCUUAUU
UCACAGAGCAACCAAUUGAUCUUGUACCAAACCAACCUUACCCUAAUGCAAGCUU
UGAUAAUUUCAAGUUUGUAUGUGAUAAUAUUAAAUUUGCUGAUGAUCUAAACCA
GCUGGCUGGUUAUAAGAAACCUGCUUCGAGAGAACUUAAAGUUACAUUUUUUCCU
GACUUAAAUGGUGAUAUAGUAGCUAUUGAUUACAAACACUACACGCCUUCUUUUA
AGAAAGGUGCUAAGUUACUGCACAAACCUGUUGUUUGGCAUGUUAACAAUACAAC
UAACAAAGCUACGUAUAAACCAAAUACUUGGUGUAUACGUUGUCUCUGGAGUACA
AAACCAGUUGAAACAUCAAAUUCAUUUGAUGUACUGGAGUCAGAAGACACACAGG
GAAUGGAUAAUCUUGCCUGUGAAGAUCUCAAACACGUCUCUGAAGAAGUAGUGG
AAAAUCCUACCAUACAGAAAGACAUAUUAGAGUGUAAUGUAAAAACUACCGAAG
UUGUAGGUGACGUUAUACUCAAACCAGCAAAUGAUGGUGUAAAAAUU???????????
????????????????????????????????????????????????????????????????????????????????
????????????????????????????????????????????????????????????????????????????????
???????????UACUAACAUAGUUACGCGAUGUUUAAAUCGUGUAUGUACUAAUUAUA
UGCCUUAUUUCUUUACUCUAUUGCUACAAUUAUGUACUUUUACUAAGAGUACUAA
UUCUAGGAUUAAAGCAUCUAUGCCAACCACUAUAGCAAAGAAUACUGUUAAAAGU
GUUGGUAAAUUUGUGUAGAG?????????????????????????UAAUUUUCUAAAGUGA
UAAAUGUUGUAAUUUGGUUUUUACUAUUAAGUGUUUGUUUAGGUUCUUUAAUCU
AUUCAACUGCUGCAUUAGGUGUUUUAAUGUCUAAUCUGGGCAUGCCUUCUUAUUG
UACAGGUUACAGAGAAGGUUACUUAAACUCUACUAAUGUCACUACUGCAACAUAC
UGUACUGGUUCUAUACCUUGUAGUGUUUGUCUUAGUGGUUUAGAUUC???????????
????????????????????????????????????????????????????????????????????????????????
??????CUCUUUACUAGGUUCUUUUAUGUACUUGGACUAGCUGCAAUUAUGCAAUUG
UUUUUCAGCUAUUUUGCUGUACAUUUUAUUAGUAAUUCUUGGCUUAUGUGGUUG
AUAAUUAACCUUGUACAAAUGGCUCCAAUUUCAGCUAUGGUCAGAAUGUACAUAU
UCUUUGCUUCAUUCUAUUAUGUAUGGAAAAGUUAUGUGCAUGUUGUGGAUGGUU
GUACUUCAUCUACUUGUAUGAUGUGUUACAAGCGUAAUAGAGCAACAAGAGUUG
AAUGUACAACUAUUGUUAAUGGUGUUAGAAGAUCCUUUUAUGUCUAUGCUAAUG
GAGGUAAAGGAUUUUGUAAACUACACAACUGGAAUUGUGUCAAUUGCGACACAU
UCUGUGCUGGUAGUACCUUUAUUAGUGAUGAAGUUGCAAGGGACUUAUCAUUAC
AGUUUAAGAGACCAAUUAACCCAACUGACCAGUCAUCAUAUGUUGUUGACAGUGU
UAGCGUAAAGAAUGGUUCUAUCCAUCUUUAUUUC??????????????????????????????
?????????ACUUUGUCAAUUUGGACAAUUUAAGGGCUAAUAACACUAAAGGUUCAU
UGCCUAUUAAUGUUAUAGUAUUUGAUGGUAAAUCUAAAUGUGAAGAAUCAUCUG
CAAAAGCAGCUUCUGUUUAUUACAGUCAGCUUAUGUGUCAGCCUAUACUACUACU
AGAUCAGG???????????????UUGGUGAUAGUACAGAGGUUGCAGUUAAAAUGUUUGA
UGCUUAUGUUAAUACAUUUUCAUCAACCUUUAAUGUGCCAAUGGAGA???????????
???????????????AGCUGAACUUGCAAAGAAUGUGUCUUUAGACAAUGUCUUAGCUAC
UUUUAUCUCAGC???????????????UGUUGAUUCAGAUGUAGAAACUAAGGAUGUUGU
UGAGUGUCUUAAAUUGUCACACCAAUCUGACAUUGAAGUUACUGGUGAUAGUUG
UAAUAAUUACAUGCUCACCUAUAACAAAGUUGAAAACAUGACACCUCGUGAUCUU
GGUGCUUGUAUCGAUUGCAGUGCACGUCAUAUUAAUGCACAAGUAGCAAAAAGUC
ACAAUAUUGCCCUGAUCUGGAAUAUUAAAGAUUUCAUGUCACUGUCUGAACAACU
ACGAAAACAAAUACGUAGUGCUGCAAAGAAGAACAACCUGCCAUUUAAGUUGACA
UGCGCAACUACUAGGCAAGUUGUUAAUGUUGUUACAACAAAGAUAGCACUUAAA
GGUGGUAAAAUUGUAAAUAACUGGCUGAAGCAGCUGAUUAAAGUUACAUUAGUG
UUUCUUUUGGUUGCUGCUACCUUUUAUUUAAUAACACCUGUUCAUGUCAUGUCCA

AACAUACUGACUUUGCAAGUGAAAUUAUAGGAUAUAAGGCUAUUGAUGGCGGUG
UCACACGUGACAUAUCAUCGACAGAUACUUGUUUUGCCAACAAACAUGCUGAUUU
UGAUACAUGGUUUAGUCAGCGUGGUGGUAGUUAUACUAAUGACAAAGCAUGCCC
AUUAGUAGCAGCUGUCAUAACAAGAGAGGUGGGUUUUGUUGUACCUGGUUUACC
UGGUACGAUACUACGCGCAACUAAUGGUGACUUCUUGCAUUUCUUACCUAGAGUU
UUCAGUGCAGUUGGCAACAUCUGCUAUACACCUUCAAAACUUAUAGAGUACACAG
ACUUUGCAACGUCAGCUUGUGUUUUAGCUGCUGAAUGCACAAUUUUCAAGGAUGC
UUCUGGUAAACCAGUGCCAUAUUACUAUGACACUAAUGUAUUAGAAGGUUCUGU
UUCUUAUGAAAGU???????????????????????????????????UAUAAUUCAAUUCCCUAA
CACUUACCUUGAAGGUUCUGUUAGAGUGGUAACAACUUUUGACUCUGAGUACUGU
AGACACGGUACAUGUGAAAGAUCUGAAGCAGGCAUUUGUGUAUCCACUAGUGGU
AGAUGGGUACUCAAUAAUGAUCAUUACAGAUCUCUACCAGGAGUGUUUUGUGGU
GUAGAUGCUGUGAAUUUACUUACUAAUAUGUUCACACCACUAAUUCAACCGAUUG
GUGCUUUGGACAUAUCUGCAUCUAUUGUAGCGGGAGGCAUCGUUGCUAUUAUAG
UAACAUGUCUUGCUUACUACUUUAUGAGGUUUAGAAGAGCUUUUGGUGAAUACA
GUCAUGUAGUUGCCUUCAAUACUCUGUUAUUCCUUAUGUCAUUCACUGUACUCUG
CUUGACACCAGUUUAUUCGUUCUUACCUGGUGUUUACUCUGUUAUUUACUUGUAC
UUGACAUUCUAUCUUACUAAUGAUGUUUCUUUCUUAGCACAUAUCCAAUGGAUGG
UUAUGUUCACACCUUUAGUACCUUUCUGGAUAACAAUUGUUUAUGUCAUUUGUA
UUUCCACAAAGCAUUUUUAUUGGUUCUUUAGCAACUACCUAAAGAGACGUGUUGU
CUUUAAUGGUGUUUCCUUUAGUACAUUUGAAGAGGCUGCUUUAUGCACCUUUCUC
UUAAAUAAAGAAAUGUAUCUGAAAUUGCGCAGUGAUGUACUUCUGCCUCUUACGC
AAUAUAACAGAUAUUUAGCUCUUUACAAUAAGUACAAGUAUUUUAGUGGAGCCA
UGGACACUACAAGCUAUAGAGAAGCUGCUUGUUGUCAUCUCGCCAAGGCUCUUAA
UGAUUUUAGCAACUCAGGCUCUGAUGUUCUCUACCAACCACCACAAACUUCUAUC
ACAUCUGCUGUCUUACAGAGUGGUUUUAGAAAAAUGGCAUUCCCAUCUGGUAAAG
UUGAGGGCUGCAUGGUACAAGUUACUUGUGGUACAACUACUCUUAAUGGUCUUU
GGCUUGAUGAUGUAGUUUACUGUCCACGACAUGUGAUCUGCACUUCUGAAGACAU
GCUUAACCCUAAUUAUGAAGAUUUGCUCAUUCGUAAG??????????????????????????
GUAAUGUUCAACUUAGAGUAAUUGGACAUUCUAUGCAAAAUUGUGUUCUUAAGC
UUAAAGUUGACACAGCUAAUCCUAAGACACCUAAGUAUAAGUUUGUGCGCAUACA
ACCAGGACAGACUUUUUCAGUACUAGCUUGUUAUAAUGGUUCACCAUCAGGUGUU
UACCAAUGUGCUAUGAGAC???????????????????UCAUUCCUUAAUGGUUCUUGUGG
UAGUGUUGGUUUUAACAUAGAUUAUGACUGUGUCUCUUUUUGCUACAUGCACCAC
AUGGAAUUACCAACUGGAGUUCAUGCUGGCACAGAUUUAGAAGGUACCUUCUAUG
GGCCUUUUGUUGACAGACAAACAGCACAAGCAGCUGGUACAGACACAACUAUCAC
??????????????????UGUAUGCUGCUGUUAUAAAUGGAGAUAGGUGGUUUCUCAAUC
GAUUCACCACAACUCUUAAUUACUUUAACCUUGUGGCUAUGAAGUACAACUAUGA
ACCUUUGACACAAGAUCAUGUUGACAUACUAGGACCUCUUUCAGCUCAAACUGGA
AUUGCAGUUCUAGAUAUGUGUGCUUCCUUAAAAGAAUUACUACAAAAUGGUAUG
AAUGGACGUACCAUAUUGGGUAGUGCUUUAUUAGAAGAUGAAUUUACACCUUUU
GAUGUUGUUAGACAAUGCUCAGGUGUUACUUUUCAGAGUGCAGUAAAGAGGACA
AUCAAGGUAC????????????????????????????????????????????????????????????????
?????????????????????????????????????????????????????????????????UUGUUACCUU
CUCUUGCUACUGUAGCUUAUUUUAACAUGGUCUACAUGCCUGCUAGUUGGGUGAU
GCGUAU??????????????????GUUGACACUAGUUUGUCUGGUUUCAAACUAAAGGACU
GUGUUAUGUAUGCAUCAGCUGUAGUGUUAUUAAUCCUUAUGACAGCAAGAACUG
UGUAUGAUGAUGGUGCUAGAAGAGUUUGGACACUUAUGAAUGUUCUGACACUUG
UUUAUAAAGUCUAUUAUGGUAAUGCUUUAGAUCAAGCUAUUUCUAUGUGGGCUC
UUAUAAUCUCUGUAACUUCUAACUACUCAGGUGUAGUUACAACUGUCAUGUUUAU
GGCCAGAGGUAUUGUUUUUAUGUGUGUUGAGUAUUGCCCUAUCUUCUUCAUAAC
UGGUAAUACACUUCAGUGUAUAAUGCUAGUUUAUUGUUUCUUAGGCUAUUUCUG

UACUUGUUACUUUGGCCUCUUCUGUUUACUCAACCGCUAUUUUAGACUGACUCUU
GGUGUUUAUGAUUAUUUAGUUUCUACACAGGAGUUUAGAUAUAUGAAUUCCCAA
GGAUUACUUCCUCCUAAGAAUAGCAUAGAUGCCUUCAAACUUAACAUCAAGUUGU
UGGGUGUUGGAGGUAAACCAUGCAUUAAAGUAGCCACUGUACAGUCUAAAAUGU
CAGAUGUAAAGUGUACGUCAGUAGUUUUACUUUCAGUUUUACAACAACUUAGAG
UAGAAUCGUCUUCUAAAUUGUGGGCUCAAUGUGUUCAGCUCCAUAAUGAUAUUCU
CUUAGCUAAGGAUACUACUGAAGCCUUUGAAAAAAUGGUUUCAUUACUUUCUGU
UCUGCUUUCUAUGCAAGGUGCUGUAGACAUAAACAAGCUUUGUGAAGAAAUGCUC
GAUAACAGGGCAACCUUACAAGCCAUAGCUUCAGAGUUUAGUUCUCUCCCAUCAU
AUGCAGCUUUUGCUACUGCUCAGGAAGCUUAUGAGCAGGCUGUUGCUAAUGGUGA
CUCUGAAGUUGUUCUUAAAAAGUUAAAGAAAUCUUUGAAUGUGGCUAAAUCUGA
AUUUGACCGUGAUGCAGCUAUGCAACGUAAGUUGGAGAAGAUGGCUGAUCAAGC
UAUGACCCAGAUGUACAAACAGGCAAGAUCUGAAGACAAAAGGGCAAAAGUUACU
AGUGCUAUGCAAACAAUGCUUUUCACUAUGCUUAGAAAGUUGGAUAAUGAUGCA
CUUAACAACAUUAUCAACAAUGCAAGAGAUGGUUGUGUACCGUUGAACAUAAUAC
CACUCACUACUGCAGCCAAAUUAAUGGUUGUCAUACCAGACUAUAACACAUAUAA
GAACACGUGUGAUGGUACUACUUUUACUUAUGCAUCAGCACUAUGGGAAAUCCAG
CAAGUUGUUGAUGCAGAUAGUAAAAUUGUUCAGCUUAGUGAGAUUAGUAUGGAC
AAUUCACCUAAUCUAGCAUGGCCUCUCAUUGUAACAGCCUUGAGGGCCAAUUCUG
CUGUCAAAUUACAGAAUAAUGAGCUUAGUCCUGUUGCACUACGACAGAUGUCAUG
UGCCGCCGGUACAACACAAACAGCAUGCACUGAUGAUAAUGCUCUAGCCUACUAU
AACACUACAAAGGGAGGUAGGUUUGUAUUAGCAUUACUAUCUGAUUUACAAGAC
UUGAAGUGGGCUAGGUUCCCUAAGAGUGAUGGAACUGGCACUAUUUAUACGGAA
CUGGAACCACCUUGUAGGUUUGUUACAGACACACCAAAGGGUCCUAAAGUGAAAU
ACUUGUAUUUUAUUAAGGGUCUAAACAAUCUAAAUAGAGGUAUGGUAUUGGGUA
GUUUAGCUGCUACAGUACGCUUACAGGCUGGCAAUGCAACAGAAGUACCUGCCAA
UUCAACUGUGCUAUCUUUUUGUGCUUUUGCUGUAGAUGCAGCUAAGGCUUAUAA
AGAUUACCUAGCUAGUGGAGGACAACCAAUCACUAAUUGUGUUAAGAUGUUGUG
UACACACACUGGUACUGGUCAGGCAAUAACAGUUACACCAGAAGCCAAUAUGGAU
CAAGAAUCCUUUGGCGGUGCAUCGUGUUGUCUGUACUGUCGUUGCCACAUAGAUC
AUCCAAAUCCUAAAGGGUUUUGUGAUUUGAAAGGUAAAUAUGUACAAAUACCUA
CAACUUGUGCUAAUGACCCUGUGGGUUUUACACUUAAAAACACAGUCUGUACCGU
CUGCGGUAUGUGGAAAGGUUAUGGCUGUAGUUGUGAUCAACUCCGCGAACCCAUG
CUUCAGUCAGCUGACGCACAGUCGUUUUUAAACGGGUUUGCGGUGUAAGUGCAGC
CCGUCUUACACCGUGCGGCACAGGCACUAGUACUGAUGUCGUAUAUAGGGCUUUU
GACAUCUACAAUGACAAAGUAGCUGGUUUUGCUAAAUUCCUAAAAACUAAUUGU
UGUCGCUUCCAAGAGAAAGAUGAAGAUGGCAAUUUAAUUGACUCUUAUUUCAUA
GUUAAGAGACACACUUUCUCUAACUAUCAACAUGAGGAAACAAUUUACAACUUAC
UUAAGGAUUGUCCAGCUGUUGCUAAACAUGACUUUUUUAAGUUUAGAAUAGACG
GUGACAUGGUACCACAUAUAUCACGUCAACGUCUUACUAAAUACACAAUGGCUGA
CCUUGUCUAUGCUUUGCGGCAUUUUGAUGAGGGUAACUGUGACACAUUAAAAGA
AAUACUUGUUACUUACAACUGUUGUGAUGAUGAGUAUUUUAACAAAAAAGACUG
GUAUGAUUUUGUAGAAAACCCAGACAUAUUACGCGUAUAUGCUAACUUAGGUGA
GCGUGUACGCCAAGCUUUGUUAAAAACAGUACAAUUCUGUGAUGCCAUGCGAGAU
GCUGGCAUUGUUGGUGUACUGACAUUAGAUAAUCAAGAUCUUAACGGUAACUGG
UAUGAUUUCGGUGAUUUCAUACAGACCACACCAGGUAGUGGAGUUCCCGUUGUAG
AUUCUUAUUAUUCAUUGUUAAUGCCUAUAUUAACAUUGACAAGAGCAUUAACUG
CUGAGUCACAUGUUGACACUGAUCUAACAAAGCCUUACAUAAAAUGGGAUUUGUU
AAAGUAUGAUUUCACGGAAGAGAGGUUAAAACUCUUUGACCGUUAUUUCAAGUA
UUGGGAUCAAACUUAUCACCCAAAUUGUGUUAAUUGUUUGGAUGACAGAUGCAU
UCUGCAUUGUGCAAACUUUAAUGUUUUGUUCUCUACGGUCUUCCCACCAACAAGU
UUUGGUCCUUUAGUGAGAAAGAUUUUUGUUGAUGGUGUUCCAUUUGUUGUUUCA

ACUGGUUACCACUUCAGAGAGCUAGGAGUUGUACAUAAUCAGGAUGUAAACUUAC
AUAGCUCCAGACUUAGUUUUAAGGAAUUACUUGUGUAUGCUGCUGAUCCUGCUAU
GCAUGCUGCUUCUGGUAAUUUAUUACUAGAUAAGCGUACAACAUGCUUUUCAGUA
GCUGCACUUACCAACAACGUGGCAUUUCAAACUGUCAAACCAGG??AUUUUAAUAA
AGACUUUUAUGACUUUGCAGUCUCUAAAGGUUUUUUCAAGGAAGGAAGUUCUGU
UGAAUUAAAACACUUCUUCUUUGCCCAAGAUGGUAAUGCAGCAAUAAGUGAUUA
UGAUUACUAUCGCUACAAUUUACCAACUAUGUGUGACAUUAGACAAUUACUUUUC
GUAGUAGAAGUAGUUGAUAAGUAUUUUGAUUGCUAUGACGGUGGUUGUAUUAAU
GCUAAUCAAGUCAUAGUUAAUAAUUUAGACAAGUCUGCUGGUUUUCCAUUUAAU
AAAUGGGGCAAGGCUAGGUUAUAUUAUGAUUCUAUGAGUUAUGAGGACCAAGAU
GCAUUGUUCGCUUAUACUAAGCGUAAUGUCAUCCCAACUAUAACUCAAAUGAAUC
UUAAAUAUGCUAUUAGUGCUAAAAAUAGAGCUCGUACAGUUGCUGGCGUAUCUA
UUUGUAGCACUAUGACAAACAGACAGUUCCAUCAGAAACUUCUUAAGUCUAUAGC
AGCCACCAGAGGUGCCACAGUUGUUAUAGGCACUAGUAAGUUCUAUGGUGGUUGG
CAUAAUAUGUUGAAAACUGUUUACAGUGAUGUAGAAAAUCCCCAUCUUAUGGGU
UGGGAUUACCCUAAAUGUGACAGAGCAAUGCCUAACAUGCUUAGAAUCAUGGCCU
CACUCGUGCUUGCUCGUAAACAUACAACCUGUUGCAGUCUGUCACACCGUUUCUA
UAGAUUAGCUAAUGAGUGUGCACAGGUAUUAAGUGAAAUGGUCAUGUGUGGUGG
UUCACUAUAUGUUAAACCAGGUGGAACUUCAUCAGGAGAUGCAACAACUGCUUAU
GCUAAUAGUGUUUUUAACAUUUGUCAAGCUGUUACAGCUAAUGUCAAUGCACUU
UUAUCCACUGAUGGUAACAAAAUUGCUGAUAAAUAUAUCCGCAAUUUGCAGCACA
GACUUUAUGAGUGUCUCUAUAGAAAUAGAGAUGUUGAUACAGACUUUGUGAAUG
AGUUUUAUGCAUAUUUGCGUAAACACUUCUCAAUGAUGAUACUCUCUGAUGAUGC
UGUUGUGUGCUUUAAUAGCACUUAUGCGUCUCAAGGUUUAGUGGCUAGCAUAAA
GAACUUCAAGUCAGUUCUUUAUUACCAAAAUAAUGUUUUUAUGUCUGAGGCUAA
AUGCUGGACUGAGACUGACCUUACUAAAGGACCUCAUGAAUUUUGCUCUCAGCAU
ACAAUGCUAGUCAAACAAGGUGAUGAUUAUGUGUACCUGCCCUAUCCUGAUCCAU
CAAGAAUUUUAGGAGCUGGCUGUUUUGUUGAUGACAUCGUAAAAACAGAUGGUA
CAUUAAUGAUAGAACGAUUUGUGUCUUUAGCUAUAGAUGCUUAUCCACUUACUA
AACAUCCAAAUCAGGAGUAUGCUGAUGUCUUUCAUUUGUAUUUACAAUACAUAA
GAAAGUUACAUGAUGAAUUAACAGGACAUAUGUUAGACAUGUAUUCUGUUAUGC
UUACUAAUGAUAACACUUCUAGGUAUUGGGAACCUGAAUUUUAUGAAGCUAUGU
ACACACCUCAUACAGUCUUACAGGCUGUUGGAGCCUGUGUUCUUUGUAAUUCACA
GACUUCAUUAAGAUGUGGUGCGUGUAUACGCAGACCAUUCUUAUGUUGUAAAUG
CUGUUAUGACCAUGUCAUAUCAACAUCACAUAAAUUAGUCUUGUCUGUUAAUCCU
UAUGUUUGCAAUGCUCCAGGUUGUGAUGUCACAGAUGUGACUCAACUUUACUUAG
GAGGUAUGAGCUAUUACUGCAAGUCACACAAACCGCCUAUUAGCUUUCCUUUAUG
UGCUAAUGGACAGGUUUUUGGUUUAUAUAAAAACACAUGUGUUGGUAGCGACAA
CGUUACUGACUUUAAUGCAAUAGCCACAUGUGAUUGGACAAAUGCAGGUGAUUAC
AUUCUUGCUAACACCUGUACUGAGAGACUUAAACUGUUCGCUGCUGAAACAUUGA
AAGCAACAGAAGAGACCUUUAAACUAUCUUACGGCAUUGCCACUGUGCGUGAAGU
GUUGUCUGAUAGAGAGUUACACCUUUCAUGGGAGGUUGGAAAACCUAGACCACCA
CUCAAUAGAAAUUAUGUCUUUACUGGUUACCGUGUAACUAAAAAUAGUAAAGUA
CAAAUAGGAGAGUACACCUUUGAAAAAGGUGACUAUGGAGAUGCUGUUGUAUAU
CGAGGUACAACAACCUACAAAUUAAAUGUUGGUGACUAUUUUGUACUAACAUCAC
AUACAGUAAUGCCUUUGAGUGCGCCUACACUAGUACCACAAGAGCAUUAUGUUAG
AAUAACUGGCUUGUACCCGACACUCAACAUCUCAGAUGAGUUUUCUAGCAAUGUU
GCAAAUUAUCAAAAGGUUGGUAUGCAAAAGUAUUCUACACUCCAGGGACCUCCUG
GUACUGGUAAGAGUCAUUUUGCUAUUGGCUUAGCUCUCUACUACCCGUCUGCGCG
CAUAGUGUAUACAGCUUGCUCUCAUGCUGCUGUCGAUGCGCUUUGCGAGAAGGCA
UUAAAAUAUUUGCCUAUAGACAAAUGUAGUAGAAUUAUACCUGCACGCGCUCGUG
UAGAGUGUUUUGACAAAUUCAAAGUGAAUUCAACAUUAGAACAGUAUGUCUUUU

GCACUGUAAAUGCAUUGCCAGAAACAACUGCUGAUAUAGUUGUUUUUGAUGAAA
UUUCAAUGGCUACAAAUUAUGACUUGAGUGUUGUCAAUGCUAGACUACGUGCUA
AGCACUAUGUUUACAUUGGCGAUCCUGCUCAACUACCAGCACCACGCACAUUGCU
AACUAAAGGCACACUAGAACCAGAAUAUUUUAAUUCAGUGUGUAGACUUAUGAA
AACUAUAGGUCCAGACAUGUUCCUUGGAACCUGUCGUCGCUGUCCUGCUGAAAUA
GUCGACACUGUAAGUGCUCUAGUUUAUGACAAUAAGCUGAAAGCACAUAAAGAA
AAAUCAGCACAAUGCUUUAAAAUGUUUUAUAAGGGUGUUAUUACACAUGAUGUC
UCAUCUGCAAUAAACAGACCUCAAAUAGGCGUAGUAAGAGAAUUUCUUACACGCA
AUCCUGCUUGGAGAAAAGCUGUCUUUAUCUCACCAUAUAAUUCACAGAAUGCGGU
AGCGUCAAAAAUCUUGGGACUACCAACUCAGACUGUUGAUUCAUCACAGGGUUCU
GAAUAUGACUAUGUCAUAUUCACGCAAACCACUGAAACAGCUCACUCUUGUAAUG
UUAAUAGAUUUAAUGUUGCUAUUACUAGAGCGAAAGUAGGCAUACUUUGCAUAA
UGUCAGAUAGAGACCUUUAUGACAAGUUGCAAUUUACAAGUCUUGAAAUUCCACG
UAGAAAUGUGGCAACUUUACAAGCAGAAAAUGUAACAGGACUAUUUAAAGAUUG
UAGCAAAGUGAUCAAUGGAUUACAUCCUACACAAGCACCUACACACCUCAGUGUU
GAUACCAAAUUUAAAACUGAAGGUCUAUGUGUUGACAUACCAGGUAUACCCAAGG
ACAUGACCUAUAGGAGACUCAUUUCCAUGAUGGGUUUCAAAAUGAAUUAUCAAG
UUAAUGGUUACCCUAACAUGUUCAUCACCCGAGAAGAAGCCAUAAGACAUGUACG
CGCAUGGAUUGGUUUCGAUGUCGAAGGGUGUCAUGCUACAAGAGAAGCUGUAGG
UACUAAUUUGCCUUUACAGUUAGGCUUUUCUACAGGUGUUAAUUUAGUUGCUGU
ACCCACAGGCUAUGUUGACACACCUAAUAAUACAGAUUUCACCAGAGUUAGUGCU
AAGCCACCACCUGGAGACCAGUUUAAACAUCUUAUACCACUCAUGUACAAAGGUU
UGCCUUGGAAUGUAGUGCGUAUAAAGAUAGUUCAGAUGUUAAGUGACACACUUA
AAAAUCUUUCUGACAGAGUUGUGUUCGUACUUUGGGCACACGGCUUUGAAUUAAC
AUCCAUGAAGUAUUUUGUAAAAAUAGGUCCUGAACGCACUUGCUGUCUCUGUGAC
AGACGUGCUACCUGUUUUUCCACAGCUUCUGAUACUUAUGCAUGCUGGCAUCACU
CAAUUGGGUUCGACUACGUCUAUAAUCCUUUCAUGAUUGAUGUUCAGCAAUGGGG
UUUUACAGGUAACUUACAGAGUAACCAUGACUUGUAUUGUCAAGUACAUGGUAA
UGCACAUGUUGCUAGUUGUGAUGCUAUCAUGACUAGAUGUCUGGCAGUUCAUGA
AUGCUUUGUUAAGCGUGUUGACUGGACUGUAGAGUACCCUAUAAUAGGUGAUGA
ACUGAAGAUUAAUGCAGCUUGCAGAAAAGUACAGCACAUGGUUGUUAAGGCUGC
AUUACUUGCAGAUAAAUUCUCAGUUCUUCACGACAUUGGUAACCCUAAAGCUAUU
AAGUGUGUACCGCAGGCUGAAGUUGAGUGGAAAUUCUACGAUGCUCAGCCCUGUA
GUGAUAAAGCUUACAAAAUAGAAGAAUUGUACUACUCGUAUGCUACACACUCUGA
UAAGUUUACAGAUGGUGUUUGUUUAUUCUGGAAUUGCAAUGUAGAUAGAUACCC
UGCUAAUUCUAUUGUGUGUAGAUUUGAUACUAGAGUAUUAUCAAACCUAAACUU
ACCAGGUUGUGAUGGUGGUAGUUUAUAUGUCAACAAACAUGCCUUUCACACACCA
GCAUUUGAUAAGAGUGCCUUUGUCAAUUUAAAACAAUUGCCUUUCUUCUACUACU
CUGAUAGCCCCUGCGAAUCUCAUGGAAAACAGGUUGUGUCAGAUAUAGAUUAUGU
ACCACUAAAAUCUGCUACGUGUAUAACACGUUGUAAUUUAGGUGGUGCUGUUUG
UAGACAUCAUGCUAAUGAGUAUAGAUUAUAUCUUGACGCUUAUAAUAUGAUGAU
CUCAGCUGGCUUUAGCUUAUGGGUUUAUAAACAAUUUGAUACUUACAACCUCUGG
AAUACUUUUACAAGACUUCAGAGUUUAGAAAAUGUGGCUUUCAAUGUUGUAAAU
AAAGGACACUUUGAUGGACAACAGGGUGAAGUACCAGUUUCCAUCAUUAAUAACA
CUGUUUACACAAAAGUUGAUGGUGUUGAUGUAGAAUUAUUUGAAAACAAAACAA
CAUUACCAGUUAAUGUAGCAUUUGAGCUUUGGGCUAAACGCAACAUUAAACCGGU
ACCAGA??????????????????CUUGGGUGUUGACAUUGCUGCUAAUACAGUGAUUUGG
GACUAUAAAAGAGAAGCCCCUGCACAUGUUUCUACAAUAGGAGUUUGUACUAUGA
CUGACAUAGCAAAGAAACCUACUGAAAGUGUUUGCGCACCUCUCACCGUCUUCUU
UGAUGGUAGAGUUGAUGGCCAAGUAGACUUGUUCAGAAACGCCCGUAAUGGUGU
UCUUAUUACAGAAGGCAGUGUUAAAGGUUUACAACCAUCUGUUGGUCCUAAACAA
GCUAGUCUUAAUGGAGUCACAUUAAUUGGAGAAGCAGUAAAAACACAGUUCAAU

UAUUACAAGAAAGUAGAUGGUGUUGUACAGCAACUACCUGAAACUUAUUUUACCC
AAAGUAGAAAUUUACAAGAAUUCAAACCCAGGAGUCAAAUGGAAAUUGAUUUCU
UAGAAUUAGCUAUGGAUGAAUUCAUUGAACGAUAUAAACUAGAAGGCUAUGCCU
UCGAACAUAUCGUUUAUGGAGAUUUUAGUCACAGUCAAUUAGGGGGCUUACACU
UAUUGAUUGGACUAGCUAAACGUUCAAAGGAUUCGCCUCUCGAGUUAGAGGAUU
UUAUUCCCAUGGACAGUACAGUUAAAAAUUACUUCAUAACAGAUGCACAAACUGG
AUCUUCAAAAUGUGUGUGUUCUGUUAUAGAUUUAUUACUUGAUGAUUUUGUUGA
AAUAAUAAAAUCUCAAGAUUUAUCUGUGGUUUCUAAAGUUGUCAAAGUGACUAU
UGACUAUACAGAAAUUUCAUUUAUGCUUUGGUGUAAAGAUGGACACGUUGAAAC
AUUUUACCCAAAAUUACAAUCUAGUCAAGCAUGGCAACCGGGAGUGGCUAUGCCA
AACCUUUACAAAAUGCAAAGGAUGCUACUAGAGAAAUGUGACCUUCAGAAUUAU
GGUGAUAGUGCUACAUUACCUAAAGGCAUAAUGAUGAAUGUCGCAAAAUAUACCC
AACUGUGUCAAUAUUUAAAUACAUUAACUUUAGCUGUGCCUUACAAUAUGAGAG
UUAUACAUUUUGGUGCUGGCUCAGAUAAAGGAGUGGCACCUGGUACAGCAGUUU
UGAGACAGUGGUUACCCACGGGUACACUACUUGUUGAUUCAGAUCUUAAUGACUU
UGUCUCUGAUGCAGAUUCAACUUUAAUUGGUGAUUGUGCAACCGUACAUACAGCU
AACAAAUGGGAUCUCAUUAUUAGUGACAUGUACGAUCCUAAGACUAAAAAUGUU
ACAAAAGAAAAUGAUUCCAAAGAAGGAUUUUUCACUUACAUUUGUGGAUUUAUA
CAACAAAAGUUAGCCCUCGGAGGUUCUGUGGCAAUAAAGAUAACGGAGCACUCUU
GGAAUGCUGAUCUUUAUAAGCUCAUGGGACACUUCGCAUGGUGGACCGCUUUUGU
UACUAAUGUGAAUGCCUCAUCUUCAGAAGCAUUUUUAAUUGGAUGUAAUUAUCU
UGGCAAACCGCGUGAACAAAUCGACGGUUAUGUCAUGCAUGCAAAUUACAUAUUU
UGGAGGAACACAAAUCCAAUACAAUUGUCUUCCUAUUCUUUAUUUGACAUGAGUA
AGUUUCCUCUUAAAUUAAGAGGUACUGCUGUAAUGUCUUUAAAAGAAGGCCAAA
UUAAUGAUAUGAUUUUAUCUCUUCUUAGUAAAGGUAGACUUAUUAUUAGAGAGA
ACAACAGAGUUGUUAUUUCUAGUGAUGUUCUUGUUAAUAACUAAACGAACAUGU
UGUUUUUCUUCUUUUUACACUUUGCCUUAGUAAAUUCACAAUGUGUUAAUUUAA
CAGGUAGAGCUGCUAUCCAGCCUUCAUUCACCAAUUCCUCUCAAAGAGGUGUUUA
UUAUCCUGACACCAUAUUUAGAUCAAACACACUUGUGUUGAGUCAGGGUUACUUU
UUACCUUUUUAUUCUAAUGUUAGCUGGUAUUAUGCAUUGACAAAAACUAACAGU
GCUGAAAAGAGAGUUGAUAACCCUGUUUUGGAUUUCAAAGACGGUAUUUACUUU
GCUGCAACUGAAAAAUCUAACAUUGUCAGAGGUUGGAUCUUUGGAACGACUCUUG
ACAACACAUCACAGUCACUUUUGAUAGUUAACAACGCAACUAAUGUUAUCAUCAA
AGUUUGUAAUUUCCAGUUUUGUUAUGACCCUUACCUUAGUGGUUAUUAUCAUAA
CAAUAAAACGUGGAGCACGAGAGAGUUUGCUGUUUAUUCCUCUUAUGCCAAUUGC
ACUUUUGAGUAUGUGUCUAAGUCUUUUAUGCUAGAUAUAGCUGGCAAAAGUGGC
UUAUUUGACACAUUAAGAGAGUUUGUUUUCCGAAAUGUCGACGGAUAUUUCAAG
AUUUACUCAAAAUACACACCUGUUAAUGUAAAUAGUAAUUUACCUAUAGGUUUU
UCAGCACUUGAACCUCUUGUUGAAAUUCCAGCUGGCAUAAAUAUUACUAAAUUUA
GAACACUCCUCACUAUACAUAGGAGACCCCAUGCCU????????????????????????????
???????????????????GGGCUAUUUAGCUCCACGUACAUUUAUGUUAAAUUAUAAUGAA
AAUGGUACAAUAACAGAUGCUGUUGAUUGUGCCCUAGAUCCUCUAUCUGAGGCUA
AAUGCACAUUAAAAUCCUUAACUGUUGAAAAAGGAAUCUAUCAGACUUCUAACUU
UAGAGUUCAACCAACUGAAUCUAUAGUUAGGUUUCCAAAUAUUACAAACUUAUGC
CCUUUUGGUGAAGUUUUCAAUGCAACCACUUUUGCAUCUGUUUAUGCUUGGAAUA
GAAAGAGAAUCAGUAACUGUGUUGCUGAUUACUCUGUUCUUUACAACUCCACUUC
UUUCUCAACAUUCAAAUGUUAUGGAGUUUCACCAACCAAACUAAAUGAUCUCUGC
UUUACUAACGUUUAUGCAGACUCAUUUGUAGUUAGAGGUGAUGAAGUCAGACAA
AUUGCUCCAGGACAAACAGGAAGAAUUGCUGACUAUAAUUAUAAACUCCCUGAUG
AUUUCACAGGUUGUGUAAUAGCUUGGAAUUCUAACAACCUUGAUUCUAAGGUUG
GUGGUAAUUAUAACUACCUUUAUAGAUUGUUUAGAAAGUCCAACCUCAAACCUUU
UGAACGAGACAUUUCUACAGAAAUAUACCAAGCUGGUAGUACACCCUGCAAUGGG

GUUGAAGGUUUUAACUGUUACUUUCCUCUACAAUCUUAUGGUUUCCACCCUACUA
AUGGUGUUGGUUACCAACCUUAUAGAGUAGUAGUAUUGUCAUUUGAACUUUUAA
AUGCACCUGCUACUGUUUGUGGACCUAAACAGUCCACUAACCUAGUUAAAAACAA
AUGUGUCAACUUCAAUUUUAAUGGUCUAACAGGCACAGGUGUUCUUACAGAGUCU
AGCAAAAAGUUUUUGCCUUUCCAACAAUUUGGCAGAGAUAUUGCCGACACUACUG
AUGCUGUCCGUGAUCCACAGACACUUGAAAUUCUUGAUAUCACACCGUGUUCUUU
UGGUGGUGUCAGUGUUAUAACACCAGGAACAAACACUUCUAACCAAGUGGCUGUU
CUUUAUCAGGAUGUUAACUGCACUGAAGUCCCUGUUGCUAUUCAUGCAGAUCAAU
UAACACCAACCUGG?????????????????????????????????????????AGGCUGUUUAAUAGG
GGCUGAACAUGUUAACAACACUUACGAGUGUGACAUACCAAUUGGUGCAGGAAUA
UGUGCCAGUUAUCAGACUCAAACUAAUUCACGUAGUGUUUCAAGUCAAGCUAUUA
UUGCCUACACUAUGUCACUUGGUGCAGAAAAUUCAGUUGCUUAUGCUAAUAACUC
UAUUGCCAUACCUACAAAUUUUACUAUUAGUGUGACCACUGAAAUUCUACCAGUG
UCUAUGACAAAGACAUCAGUAGAUUGUACAAUGUACAUUUGUGGUGACUCAAUA
GAGUGCAGCAACCUUUUGCUCCAAUAUGGUAGUUUUUGCACACAACUUAAUCGUG
CUUUAACUGGAAUUGCUGUUGAACAAGACAAAAACACACAGGAAGUUUUUGCACA
AGUUAAACAAAUUUACAAGACACCACCAAUAAAGGAUUUUGGUGGUUUCAACUU
UUCUCAAAUAUUACCAGAUCCAUCAAAACCAAGCAAGAGGUCAUUUAUUGAAGAU
UUACUCUUCAACAAAGUGACACUUGCUGAUGCUGGCUUCAUCAAACAAUAUGGUG
AUUGCCUUGGUGAUAUUGCCGCUAGAGAUCUUAUUUGUGCACAAAAGUUUAAUG
GCCUUACUGUUCUGCCACCUUUGCUCACAGAUGAAAUGAUUGCUCAAUACACCUC
UGCACUACUUGCAGGGACAAUCACAUCAGGUUGGACCUUUGGUGCUGGUGCAGCA
UUACAGAUACCAUUUGCUAUGCAAAUGGCUUACAGGUUUAAUGGUAUUGGAGUU
ACACAAAAUGUUCUCUACGAGAACCAAAAACUAAUUGCAAACCAAUUCAACAGUG
CAAUUGGCAAAAUUCAAGAUUCACUUUCAUCUACUGCAAGUGCACUUGGAAAACU
UCAAGAUGUUGUCAACCAAAAUGCACAGGCUUUAAACACACUUGUUAAACAACUC
AGCUCUAAUUUUGGAGCCAUUUCGAGUGUGUUAAAUGACAUUCUUUCACGUCUUG
ACAAAGUUGAGGCUGAAGUCCAAAUUGACAGGUUGAUCACUGGCAGAUUACAAA
GUUUGCAGACAUACGUGACUCAACAACUAAUUAGAGCCGCAGAAAUUAGAGCUUC
UGCUAAUCUUGCCGCAACUAAGAUGUCUGAAUGUGUUCUUGGACAAUCUAAAAGA
GUUGACUUUUGUGGUAAAGGCUACCACCUUAUGUCUUUUCCGCAGUCAGCACCUC
AUGGUGUAGUCUUUUUGCAUGUGACUUAUGUUCCAUCUCAAGAAAAGAAUUUUA
CUACUACCCCUGCCAUUUGUCAUGAAGGAAAAGCACACUUUCCUCGUGAAGGUGU
UUUCGUUUCAAACGGCACGCACUGGUUUGUAACACAAAGGAAUUUCUAUGAACCA
CAAAUUAUUACCACGGACAAUACUUUUGUCUCUGGUAGCUGUGAUGUUGUGAUU
GGAAUUGUCAACAACACAGUUUAUGAUCCUUUGCAACCAGAACUUGAUUCAUUCA
AGGAGGAGUUGGACAAAUAUUUUAAAAAUCAUACAUCACCAGAUGUUGAUUUAG
GUGACAUUUCUGGCAUCAACGCUUCAGUUGUCAACAUUCAGAAAGAAAUUGACCG
CCUCAACGAGGUUGCCAAAAAUCUAAAUGAAUCUCUCAUCGACCUCCAAGAACUU
GGAAAGUAUGAGCAGUAUAUAAAAUGGCCAUGGUAUAUUUGGCUAGGAUUUAUU
GCAG?????????????????????????????????????????????????????????????????????????????????
????????????????????????????????????????????????????????????????????????????????????????
??????????????????????????AACUGUAACUUUGAAACAAGGUGAAAUCAAGGAUGCUA
CUCCUUCAGAUUCUGUUCGCGCUACUGCAACGAUACCGAUACAAGCCACACUCCC
UUUCGGAUGGCUUAUUGUUGGCGUUGCACUUCUUGCUGUUUUUCAAAGCGCUUCC
AAAAUAAUAACACUCAAAAAGAGGUGGCAUUUAGCCCUCUCUAAGGGUGUUCACU
UUGUUUGCAACUUGCUGCUGCUGUUUGUAACAGUUUAUUCACAUCUUUUGCUUGU
UGCUGCUGGCCUUGAAGCCCCAUUUCUUUAUCUUUAUGCUUUAGUUUAUUUCUUG
CAAAGUAUAAACUUUGUGAGAAUAAUAAUGAGGCUUUGGUUGUGCUGGAAAUGC
CGUUCCAAAAAUCCUUUACUUUAUGAUGCUAACUACUUCCUGUGUUGGCAUACUA
AUUGUUACGACUAUUGUAUUCCAUACAAUAGUGUAACUUCUUCAAUUGUCAUUAC
CUCCGGUGAUGGCACAACAAAUCCCAUUACAGAACAUGACUACCAAAUUGGUGGU

UAUUUUGAGAAAUGGGAAUCUGGAGUAAAAGACUGUGUUGUAUUACACAGCUAC
UUCACUUCAGAUUACUACCAGCUGUACUCAACUCAAUUGAGCACAGACACUGGUG
UUGAACAUGUAACUUUCUUCAUCUACAAUAAAAUCGUAGAUGAGCCCGAAGAACA
UGUCCAAAUUCACACAAUCGACGGUUCAUCCGGAGUUGUUAAUCCAGCAAUGGAA
CCAAUUUAUGAUGAACCGACGACGACUACUAGCGUGCCUUUGUAAGCACAAGCUG
AUGAGUACGAACUUAUGUACUCAUUCGUUUCGGAAGAGACAGGUACGUUAAUAG
UUAAUAGCGUACUUCUUUUUCUUGCUUUCGUGGUAUUCUUGCUAGUCACACUAGC
CAUCCUUACUGCGCUUCGAUUGUGUGCGUACUGCUGCAAUAUUGUUAACGUGAGU
CUUGUGAAACCUUCUUUUUACGUUUACUCUCGUGUUAAAAAUCUGAAUUCUUCUA
GAGUUCCUGAUCUUCUGGUCUAAACGAACUAAAUAUUAUAUUAGUUUUUCUGUU
UGGAACUUUAAUUUUAGCCAUGUCAGGUGACAACGGUACUAUUACCGUUGAGGA
ACUUAAAAAGCUCCUUGAGCAAUGGAACCUAGUAAUAGGAUUCCUAUUUCUUACA
UGGAUUUGUCUUUUACAAUUUGCCUAUGCCAACAGGAAUAGGUUUUUGUACAUA
AUUAAGUUAAUUUUCCUCUGGCUGCUUUGGCCAGUAACUUUAGCUUGCUUUGUGC
UUGCUGCUGUUUACAGAAUAAAUUGGAUCACAGGUGGAAUUGCCAUUGCAAUGG
CUUGUCUUGUUGGCUUGAUGUGGCUUAGCUACUUCAUUGCUUCAUUCAGGCUGUU
UGCUCGAACGCGUUCCAUGUGGUCCUUCAACCCAGAAACAAACAUUUUGUUGAAU
GUGCCGCUCCACGGUACAAUUUUGACCAGACCGCUUCUAGAGAGUGAACUUGUAA
UUGGAGCUGUGAUCCUUCGAGGUCAUCUUCGAAUUGCUGGACACCAUCUAGGACG
CUGUGACAUCAAGGACCUGCCUAAAGAAAUCACUGUUGCUACAUCACGAACGCUU
UCUUAUUACAAAUUGGGAGCGUCGCAGCGUGUAGCAGGUGACUCAGGUUUUGCUG
CAUACAGUCGCUACAGGAUUGGCAAUUACAAAUUAAACACAGACCAUUCCAGUAG
CAGUGACAAUAUUGCUUUGCUUGUACAGUAAGUGACAACAGAUGUUUCAUCUCGU
UGACUUUCAGGUUACUAUAGCAGAGAUAUUAUUAAUUAUUAUGAGAACUUUUAA
AGUUUCCAUUUGGAAUCUUGACUACAUCAUAAAUCUCAUAAUUAAAAGUUUAUC
UAAGCCACUAACUGAAAAUAAAUAUUCUCAGUUAGAUGAAGAGCAACCAAUGGA
GAUUGAUUAAACGAACAUGAAAAUUAUUCUUUUCUUGGCAUUGAUAACACUUGC
UACUUGUGAGCUUUAUCAUUAUCAAGAGUGUGUUAGAGGUACAACAGUACUUUU
AAAAGAACCUUGCUCUUCUGGAACAUAUGAAGGCAACUCACCUUUUCAUCCUCUA
GCUGAUAACAAAUUUGCACUGACUUGCUUUAGCACUCAAUUUGCUUUUGCUUGUC
CUGACGGUGUUAAACACAUCUACCAGCUACGUGCACGAGCAGUUUCACCUAAACU
GUUCAUCAGACAAGAGGAAGUUCAAGAACUUUACUCACCAAUUUUUCUCAUAGUU
GCGGCGAUAGUGUUUAUAACACUCUGCUUCACACUUAAGAGAAAGAUAGAAUGA
GUGAGCUUUCACUAAUUGACUUCUAUCUGUGCUCUUUAGCCUUUUUGCUAUUCCU
UGUUUUAAUUAUGCUCAUUAUCUUUUGGUUUUCACUUGAACUACAAGAUCAUAA
UGAAACCUGUCAUGCCUAAACGAACAUGAAAUUUCUUGUUUUCUUAGGAAUCCUU
ACAACAGUAACUGCAUUUCAUCAGGAAUGUAGUUUACAGUCAUGUGCUCAACAUC
AACCCUAUGUAGUUGAUGAUCCCUGUCCUAUUCACUUUUACUCUCGAUGGUUUAU
CAGAGUAGGAGCUAGAAAGUCAGCACCUUUAAUUGAACUGUGCGUGGAUGAGGC
UGGUUCUAAAUCACCCAUUCAGUACAUAGAUAUAGGUAAUUACACGGUUUCCUGU
UCACCUUUUACAAUUAAUUGCCAGGAACCUAAAUUAGGCAGUCUCGUAGUACGUU
GUUCGUUCUAUGAGGACUUUUUAGAGUAUCAUGACGUUCGUGUUGUUUUAGAUU
UCAUCUAAACGAACAAACUAAAAUGUCUGAUAAUGGACCACAAAAUCAGCGAAAU
GCACCCCGCAUUACGUUUGGUGGACCCUCAGAUUCAGCUGGCAGUAACCAGAAUG
GAGAACGCAGUGGUGCACGACCUAAACAACGUCGUCCCCAAGGUUUACCCAAUAA
UACUGCGUCUUGGUUCACCGCUCUCACUCAACAUGGCAAGGAAGACCUUAGAUUC
CCUCGAGGACAAGGCGUUCCGAUUAACACCAAUAGCAGUCCAGAUGACCAAAUUG
GCUACUACCGAAGAGCUACCAGACGAAUUCGUGGUGGUGACGGUAAAAUGAAAGA
UCUCAGUCCAAGAUGGUACUUUUACUACCUAGGAACUGGGCCAGAAGCUGGACUU
CCCUAUGGUGCUAACAAAGAAGGCAUCAUAUGGGUUGCAAAUGAGGGAGCCUUGA
AUACACCUAAAGAUCACAUUGGCACCCGAAAUCCUGCUAACAAUGCUGCAAUCGU
GCUACAACUUCCUCAAGGAACAACAUUGCCAAAAGGCUUCUACGCAGAAGGGAGC

AGAGGCGGCAGUCAAGCUUCUUCUCGUUCCUCAUCACGUAGUCGCAACAGUUCAA
GAAACACAACUCCAGGCAGCAGCAGGGGAACUUCUCCUGCUAGGAUGGCUGGCAA
UGGUGGUGAUGCUGCUCUUGCUUUGCUGCUGCUUGACAGGUUGAACCAACUUGAG
AGCAAAAUGUCUGGUAAAGGCCAACAACAACAAGGCCAAACUGUCACUAAGAAAU
CCGCUGCAGAGGCUUCUAAGAAACCUCGCCAAAAACGUACUGCCACCAAACAAUA
CAAUGUAACACAAGCUUUUGGCAGACGUGGUCCAGAACAAACCCAAGGAAACUUU
GGGGAUCAAGAAUUAAUCAGACAAGGAACUGAUUACAAACAAUGGCCGCAAAUU
GCACAAUUUGCUCCUAGCGCUUCUGCAUUCUUCGGAAUGUCGCGCAUUGGCAUGG
AAGUCACACCUUCGGGAACGUGGUUGACCUACACAGGUGCCAUCAAAUUGGACGA
CAAAGAUCCAAAUUUCAAAGAUCAAGUCAUUUUGCUGAAUAAGCACAUUGACGCA
UACAAAACAUUCCCACCAACAGAGCCUAAAAAGGACAAAAAGAAGAAGGCUGAUG
AAACUCAAGCCUUACCGCAGAGACAGAAGAAACAACCCACAGUGACUCUUCUUCC
UGCUGCAGAUUUGGAUGAUUUCUCCAAACAAUUGCAACAAUCCAUGAGCAGUGCU
GAUUCAACUCAGGCUUAAACUCAUGCAGACCACACAAGGCAGAUGGGCUAUAUAA
ACGUUUUCGCUUUUCCGUUUACGAUAUAUAGUCUACUCUUGUGCAGAAUGAAUUC
UCGUAGCUACAUAGCACAAGUAGAUGUAGUUAACUUUAAUCUCACAUAGCAAUCU
UUAAUCAGUGUGUAACAUUAGGGAGGACUUGAAAGAGCCACCACAUUUUCACCGA
GGCCACGCGGAGUACGAUCGAGGGUACAGUGAAUAAUGCUAGGGAGAGCUGCCUA
UAUGGAAGAGCCCUAAUGUGUAAAAUUAAUUUUAGUAGUGCUAUCCCCAUGUGA
UUUUAAUAGCUUCUUAGGAGAAU????????????????????????????????????
