## Supplementary material for "The SARS-CoV-2-like virus found in captive pangolins from Guangdong should be better sequenced": Mitochondrial genomes assembled from SRA data published by Liu et al. (2019) (see Table 1 for details): Supplementary_information_3.pdf

**<u>Mitochondrial genomes assembled from SRA data published by Liu *et al.* (2019)</u>** (see Table 1 for details).

**<u>GD/P7L: *Manis javanica* + *Mus musculus*</u>**

>*Manis_javanica*_GDP7L_mtDNA [consensus reconstructed from 98,226 reads]
GTTAATGTAGCTTAAAACCAAAGCAAAGCATTGAAAATGCTTAGATGAGTCCACAA
ACCTCCATAAACACACAGGTTTGGTCCCAGCCTTTTTATTAATTTTCAATAGGATTAC
ACATGCAAGTATCCGCCCCCCAGTGAAAATGCCCTTCAAGTTATCACAGTACCTGAA
GGAGCTGGCATCAAGCACGCCAATAAACGCAGCTAACGACGCCTTGCAGAGCCACA
CCCCCACGGGAAACAGCAGTGATAAAAATTAGGCTATAAACGAAAGTTCGACCTAG
CCATATTGCATTGGGTTGGTAAATCTCGTGCCAGCCACCGCGGTCATACGATTAACC
CTAGCTAATAAAAAACCGGCGTAAAGCGTGCTAAGATAAATCCAATCCAAATAAAG
TTAAGCCCTGACCAGGCCGTAAAAAGCCGTGGTTGCCGTAAAAATAAACTACGAAA
GTAACTTTAATTCACATCGACACACGATAGCTAAGACCCAAACTGGGATTAGATACC
CCACTATGCTTAGCCCTAAACTTAAATAATTCACCAGACAAAATTATTCGCCAGAGA
ACTACTGGCAACAGCCTAAAACTCAAAGGACTTGGCGGTGCTTTACATCCCCCTAGA
GGAGCCTGTTCTGTAATCGATAAACCCCGATAAACCCTACCAATTCTAGCTAATACA
GCCTATATACCGCCATCTCCAGCAAACCCTAAAAAGGAAGCACAGTAAGCAAGACT
ATGAAAACATAAATACGTTAGGTCAAGGTGTAGCCCATGAATTGGGAAGAAATGGG
CTACATTTTCTAAAATAGAACACAAACGAACGCCCTAATGAAAACGAGGGCCAAAG
GAGGATTTAGCAGTAAGCTGAGAATAGAAAGCTCAACTGAACCCGGCCCTAAAGCA
CGCACACACCGCCCGTCACCCTCTTCAAATCCCCAAGAATACCTAAACATATTAGAC
AAACACCAAGGCATGAGAAGAGACAAGTCGTAACAAGGTAAGCATACTGGAAGGTG
TGCTTGGATCACCAAAGTGTAGCTTAAACAAAGCATCTGGCCTACACCCAGAAGATC
TCAATAACATGACCACTTTGAACAAATCCTAGCCCAACCAACACCCAACATATAACC
AAACAGAGCACATAAACCAAAGCATTTACTAGACTAAAGTATAGGCGATAGAAATT
ACACAACGGCGCTATAGAAAAAGTACCGCAAGGGAAAGATGAAAGATGCATTCAAA
GTACAAAAAAGCAAAGATTACCCCCTGTACCTTTTGCATAATGGACTAACTAGAAAC
ACCCTAGCAAAGAGAACTTAAGCTAGAAACCCCGAAACCAGACGAGCTACCTACGA
GCAGTTTAAAGAACCCACTCATCTATGTGGCAAAATAGTGAGAAGACTTATAGGTAG
AGGTGAAAAGCCTAACGAGCCTGGTGATAGCTGGTTGTCCAAGAAATGAATCTAAG
TTCAACTTGAAGCATACCCAAAAGCCCAAAAACTATAATGTAGGCTTCAAGTATAGT
CTAAAAAGGTACAGCTTTTTAGAAACAGAATTAAATCTTAATTAGTGAGTAAACAAT
ACAACAACCATAGTTGGCCTAAAAGCAGCCATCAATTAAGAAAGCGTCAAAGCTCA
ACAATAAGACATAAATAATACCACAAATAAAAATCAACTCCTAAACCAATATTGGA
CCAATCTATCAATAAATAGAAGAAATACTGTTAGTATGAGTAACAAGAAATAGATCT
CCTTGCACAAGCTTATATCAGAACGGATGACCCACTGATAATTAACAACGAAACAAA
TCAAACCCAAAAATAGAACTTTGTTAAATAGATTGTTAACCCAACACAGGCGTGCAT
ATTCTCAAAGGAAAGGTTAAAACAAATGAAAGGAACTCGGCAAACACAAGCCCCGC
CTGTTTACCAAAAACATCACCTCTAGCATAACCAGTATTAGAGGCACTGCCTGCCCA
GTGACTCGCGTTAAACGGCCGCGRKWYWCTGRSCGKGCAAAGGTAGCATAATCACT
TGTTCTCTAAATAAGGACTAGTATGAACGGCTAGACGAGGGTTTACTGTCTCTCATC
TGTAACCAGTGAAATTGACCTCCCCGTGAAGAGGCGGGGATAATATAATAAGACGA
GAAGACCCTATGGAGCTTTAATTAAACAGCTTAAAATTAACCCAACACCACTCTAAC
AGAGATATAACAAACCAATTAGCCTAAGCTGTCAATTTTGGTTGGGGTGACCTCGGA
GCAAAAAACAACCTACGAGCGGTCAAATCCAGACTAACAAGTCCAGATAATCCGTT
AATTGATCCAACAACTTGATCAACGGAACAAGTTACCCTAGGGATAACAGCGCAATC
CTATTCGAGAGTCCATATCGACAATAGGGTTTACGACCTCGATGTTGGATCAGGACA
TCCCAATGGTGCAGAAGCTATTAATGGTTCGTTTGTTCAACGATTAAAGTCCTACGTG
ATCTGAGTTCAGACCGGAGTAATCCAGGTCGGTTTCTATCTATTAATACACTTCTCCC

AGTACGAAAGGACAAGAGAAGTAGGGCCTACCTCACACAGGCGCCCTCGAATCAAT
AAATGACTACCCTCTTAATTTAGCCAATTCACAACAACCCCAGTCCTAGAACCAGGG
CCRGYKWGGGTAGCAAAGCACGGCAACTGCACAAGACTTAAGCTCTTGTAACAGAG
GTTCAACTCCCCTCCCTAGCAGCAATGTACACAATCAACATAATAATAATGATCATC
CCCATCCTACTAGCCGTGGCATTCCTAACGCTAGTAGAACGCAAAGTGCTAGGATAC
ATGCAACTACGAAAGGGCCYWAACATCGTGGGCCCTTGAGGCCTGCTACAACCAAT
CGCCGACGCAGTAAAACTATTCACTAAAGAACCCCTACGACCCCTAACATCCTCAAT
CACAATATTCATCATGGCACCTATCCTAGCACTGACACTCGCACTCACCATGTGAGT
ACCACTACCAATGCCACACCCACTAGTCAACATAAACCTAGGAGTGCTATTCATACT
GGCCATATCAAGCCTCGCMGTATACTCCATCTTATGATCAGGGTGAGCCTCAAACTC
CAAATACGCCCTCATTGGAGCACTACGTGCAGTGGCCCAAACCATCTCATACGAAGT
GACACTAGCAATTATCCTGCTGTCCTTACTGCTAATAAGCGGGTCCTTCGCCCTCTCC
ACCCTAATCACAACCCAAGAAAAGCTATGATTACTAGTGCCCGCATGACCCCTGGCC
ATAATATGATTCATCTCGACCCTAGCCGAGACAAATCGAGCACCCTTCGACCTAACA
GAAGGAGAATCTGAGCTCGTATCCGGCTTTAACGTTGAATACGCAGCAGGCCCATTC
GCACTCTTCTTCCTTGCCGAATACGCAAACATCATCATAATAAACATCTTGTCAGTGA
CGCTCTTCATAGGAGCCTTTCACGACCCCCACACCCCAAGCCTCTACACAGTTAATTT
TGTAGTAAAAACACTCGCCCTTACCGCCCTATTCCTATGAATCCGAGCATCCTACCCA
CGATTCCGCTACGACCAACTAATGCACCTACTATGAAAAAACTTCCTACCACTAACC
CTAGCACTGTGCATGTGACACGTATCAATACCAATCGCCCTGTCAAGCATTCCCCCA
CAATCATAAGAAATATGTCTGACAAAAGAATTACTTTGATAGAGTAAATAATAGAG
GTTTAAATCCCCTTATTTCTAGAAAAACAGGCATTGAACCTGCACCTGAGAACTCAA
AAATCTCCGTGCTACCAACTTACACCACAATCTACAGTAAGGTCAGCTAAGCAAGCT
ATCGGGCCCATACCCCGAAAATGTTGGATCACAACCTTCCCGTACTAATAAACCTCA
CTATACTAGCCATCCTAACAACCCTATTCCTAGGGACCATGCTCGTCCTAATCAGCTC
TCACTGGCTAATGATTTGAATTGGATTCGAGATAAACATACTAGCAATAGTCCCTAT
CCTAATGAAAAACTTTAGCCCACGAGCCATAGAAGCAGCCACAAAATACTTCCTAAT
CCAAGCCACCGCGTCCATACTACTAATGCTAGCCATTACCCTAGACCTGATATCTTCT
GGACAATGAACCATCACAAAAACACACAACACCCTGCCATCAGCCATCATCACCGT
AGCTATGGCCATAAAACTAGGAATAGCACCATTCCACTTCTGAGTGCCAGAAGTAAC
ACAAGGAAGCCCACTATCATCAGGCATGCTACTCCTAACCTGACAAAAAGTCGCACC
AATATCGATCCTATACCAAATAATACCCACAATCAACACAAATATACTGACAACCAT
AGCCGCACTCTCAATCCTCATCGGCGGTTGAGGAGGGCTAAACCAGACCCAACTACG
AAAAATCATAGCATACTCCTCAATCGCCCACATAGGCTGAATAGCAATAATCATAAC
ATACAACCCGGACATTGCCATTCTAAACCTACTAGTCTACATCATAATGACACTATCT
ATATTCGCAATCCTACTATACAACTCATCAACAACAACCCTATCACTATCCCACCTAT
CAAACAAGACACCACTAATCACAGCCCTCGCACTACTAATCCTACTATCACTAGGAG
GCCTCCCCCCACTGACCGGATTCATGCCCAAATGAATAATCATCCAAGAACTAACCA
AAAACAACATAGTAATAATACCAACCGTGATAGCAATAACAGCACTACTCAACCTAT
ACTTCTACGTACGCCTGGCGTACTCCACAGCACTAACCATACTCCCAACTACCAACA
ACATAAAAATAAAATGACAATTCGAAACAACAAAAATCATAAAACTAACCCCACCT
CTAATCATTTTATCCACAATAGCACTCCCACTCACCCCTATAATATCAATCCTAAACT
AGGGGCTTAGGCTACACAGACCGAGAGCCTTCAAAGCTCTAAGCAAATACAAACTA
TTTAGTCCCTGATATAAAGACTGTAGAAATTCAACCTACATCACCTAAACGCAAATC
AGGCGCTTTAATTAAGCTAAATCCTCACTAGATTGGTGGGATACAAACCCACGAAAC
TTTAGTTAACAGCTAAACACCCTAATCAACTGGCTTCAATCTACTTCTCCCGCC??????
????????????????????????????????????????????????????????????????????????????????????????
????????????????????????????????????????????????????????????????????????????????????????
???????????????ATTGGCACACTATATCTCTTATTTGGCGCCTGGGCCGGAATAGTAGGC
ACCGCACTAAGTCTTCTAATTCGCGCTGAACTAGGTCAGCCTGGGACCCTCTTAGGG
GACGACCAAATTTACAATGTGATCGTTACCGCACATGCATTTGTAATAATCTTCTTCA
TAGTCATGCCAATCATAATCGGGGGCTTTGGAAACTGACTAGTGCCCCTGATAATTG

GAGCCCCTGACATGGCATTCCCCCGTATAAACAACATAAGTTTCTGACTGCTCCCCC
CCTCTTTCCTACTTCTCCTGGCCTCTTCCATAGTCGAAGCAGGGGCTGGAACCGGCTG
AACTGTATACCCCCCCTTTAGCAGGAAATTTAGCACACGCAGGAGCATCTGTAGACCT
AACCATCTTCTCTCTTCACCTAGCAGGTGTCTCGTCAATCCTTGGGGCTATCAATTTT
ATTACAACAATCATCAACATAAAACCCCCTGCAATAAACCAATACCAAACCCCACTA
TTCGTGTGATCAGTCCTAATTACGGCCGTGCTTCTGCTACTATCTCTACCCGTACTAG
CCGCTGGCATTACCATACTGTTAACCGACCGTAATCTAAATACAACCTTTTTTGACCC
CGCAGGAGGAGGTGACCCCATTCTATACCAACACCTATTCTGATTCTTCGGACACCC
CGAAGTGTACATTCTCATCCTTCCTGGGTTTGGAATAATCTCCCACATCGTAACCTAT
TACTCCGGGAAAAAAGAGCCCTTTGGGTACATGGGCATAGTCTGAGCAATAATATCC
ATTGGCTTCCTGGGCTTCATTGTATGGGCACACCACATGTTCACAGTGGGAATAGAC
GTTGATACACGGGCCTACTTCACATCAGCCACCATAATTATTGCTATCCCCACTGGA
GTAAAGGTGTTTAGTTGACTAGCAACCCTGCACGGAGGAAACGTAAAATGGGCCCC
AGCTATACTATGGGCCCTGGGCTTTATCTTCCTGTTTACAGTCGGGGGTCTAACTGGC
ATCGTGTTGGCCAACTCGTCCCTAGATATCGTCCTCCATGATACCTACTACGTAGTAG
CCCACTTCCACTACGTTCTCTCCATGGGAGCAGTTTTCGCCATCATAGGAGGCTTCGT
CCATTGATTCCCCCTGTTCTCAGGATACACGCTCAACAACACATGGGCAAAAGTCCA
CTTTACAATCATATTCGTGGGCGTAAACATGACCTTTTTCCCCCAACACTTCCTTGGA
CTGTCTGGAATACCTCGACGATACTCGGACTACCCAGACGCTTACACAATGTGAAAC
ACTGTGTCCTCCATAGGATCCTTCATCTCATTAACCGCCGTAATACTCATGGCCTTCA
TAATCTGAGAGGCATTCGCCTCTAAACGGGAAGTCCTAATAGTGGAGTCCACAAATA
CCAACCTCGAATGACTACACGGCTGCCCACCACCCTACCACACATTCGAAGAACCTG
CCTTCGTAAATCTGGTCAAAACAAGAGAGGAAGGAATCGAACCCTCAAACAATGGT
TTCAAGCC????????????????????????????????????????????CAAAGTTA
AATTATAGGTTCAAGCCCTTTATACTTCCATGGCGTACCCGCTCCAATTAGGCTTTCA
AGATGCCACATCCCCAATCATAGAAGAATTACTCCACTTCCACGATCACACACTAAT
AATCGTGTTCTTGATCAGCTCCCTAGTACTATACATTATCTCTCTCATACTAACAACC
AAACTCACCCACACAAGTACAATAGACGCTCAAGAAGTAGAAACCATTTGAACCAT
CCTCCCCGCAATTATCCTAATCCTAATCGCCCTGCCCTCCCTACGCATCCTCTACATA
ATGGACGAGATTAACAACCCCGCCCTGACAGTAAAAACAATGGGCCATCAATGATA
CTGAAGCTACGAATACACAGACTACGAAGACCTAAGCTTCGACTCATACATAATCCC
AACTCAAGACCTAAAACCCGGCGAACTCCGACTTCTAGAAGTGGACAACCGACTTGT
ACTACCCATAGACACAACCATCCGCATGCTCATCTCATCTGAAGACGTCCTGCACTC
TTGAGCCATCCCATCCCTGGGCCTAAAAACAGATGCTATCCCGGGGCGTCTAAACCA
AACAACCCTGATATCAACCCGGCCCGGCCTATTCTACGGACAGTGCTCAGAAATCTG
TGGCTCAAATCACAGCTTCATGCCAATTGTCCTCGAACTAGTACCCCTAAAAACATTT
GAAAACTGAACTACGTCCCTACTGTAATTCATTGAGAAGCTAATAGCGCTAGCCTTT
TAAGTTAGAGATTGAGAGCACGCCTCTCCTCAATGACATGCCTCAACTAGACACTAC
AACATGATCCATTACAATTATATCCATGATCCTGACCCTCTTTGTCCTATTCCAATTA?
???????????????????????????????????????????????????????AAAAAT
GAACGAAAATCTATTCACCTCTTTCATTACCCCAGTAATAATAGGGATCCCTATTGTA
ACAATTATCATTATGTTCCCAGTAATCCTCTTCCCAACATCAAACCGACTAATCAACA
ACCGCATTGTATCCATACAACAATGACTCCTAAAACAAACATCCAAACAAATAATAA
GCATCCACAACTACAAAGGACAGACCTGAACCCTGATATTAATAACACTAATCATTT
TCATCGCATCTACTAACCTACTAGGCCTGCTACCCCACTCATTCACCCCCACAGCCCA
ACTGTCAATAAACCTGAGCATAGCCGTTCCTCTATGGGCAGCCACCGTAGTCACAGG
TTTTCGACACAATACAAAAACATCTTTAGCCCACTTCCTACCCCAGGGAACACCAAC
CCCCCTTATCCCAGTGCTAGTGATCATTGAAACAATCAGCCTGCTAATCCAACCCAT
AGCGCTCGCAGTACGACTGACAGCCAACATCACCGCCGGCCACCTGTTAATACACCT
AATCGGAAGCGCAACCCTTGCCCTAATATCAATCAACCTCACCGTGGCCACAACTAC
CTTCATCGTCCTAGTCCTGCTTACAATCCTTGAATTCGCAGTCGCACTCATTCAGGCC
TACGTCTTCACTCTTCTAATCAGCCTCTACTTACATGATAACACATAATGACCCACCA

AACCCACTCATACCACATAGTAAACCCAAGCCCTTGACCCCTAACCGGGGCCCTATC
TGCCCTCCTAATAACATCGGGCCTAGCAATATGATTCCACTTCAACTCTACAATACTA
CTCCTCCTAGGACTAACAACAAACCTCTTAACAATATATCAATGATGACGTGACATT
GTACGAGAAAGCACCTTTCAAGGCCACCACACACCCACAGTCCAAAAAGGACTACG
ATACGGCATAATCCTATTCATTGTCTCAGAAGTATTTTTCTTCGCTGGCTTCTTCTGA
GCATTCTACCACTCAAGCCTAGCACCTACCCCCGAACTAGGAGGATGCTGACCCCCC
ACAGGCATCAACCCGCTAAACCCACTGGAAGTACCTCTACTCAACACATCCGTTCTT
TTAGCCTCAGGAGTATCAATTACATGGGCACACCATAGCTTAATAGAAGGCAGCCGA
AACCACATAACCCAGGCCCTGCTCATCACAATCCTCCTAGGCATTTACTTCACACTAC
TGCAGGTCTCAGAATATTACGAAGCACCCTTCACAATCTCCGACGGCGTGTACGGCT
CTACCTTCTTTGTGGCAACTGGGKTCCACGGCCTACACGTCATCATCGGCACCTCCTT
CCTAACTGTGTGCCTACTACGACAACTAAAATACCACTTTACATCAAACCACCACTT
CGGATTCGAAGCTGCCGCCTGATACTGACACTTCGTAGATGTAGTGTGACTGTTCCT
GTATGTCTCCATTTACTGATGAGGTTCCTATTTTCTAAGTATGCACAGTACAGTTGAC
TTCCAATCAACAAGCTCTGGCCACAGCCCAGAAGAAAATAATAAACCTACTCCTAGC
AATAACAACAAACACCATACTAGCATGCCTGCTCATACTAATCGCCTTCTGACTCCC
TCAATTAAACACGTACTCGGAAAAAATCACCCCCTACGAATGCGGATTCGACCCCAT
GGGATCGGCACGACTACCATTCTCCATAAAATTCTTCCTAATCGCCATCACGTTCCTA
CTATTCGACCTAGAAATCGCACTACTCCTGCCACTCCCCTGAGCATCACAAACAAAC
AACCTAGGCACCATAATCACCGTAGCCTTGGTCCTCATCCTACTGCTAGCAATCAGC
CTGGCCTACGAATGAACACAAAAAGGCCTAGAATGAACTGAGTATGGTAGCTAGTTT
ACACAAAACAAATGATTTCGACTCATTAAACTATGACTTACTCATAGCTACCAAATG
TCCCTAATTTACATTAACACAACACTGGCATTCACTATCTCCCTAATGGGGATATTAA
TGTACCGATCACACCTAATGTCCTCACTACTATGTCTAGAGGGCATAATACTCTCCCT
ATTCGTAATAATCACCATTACAATCCTAACTAACCACTTCACACTAGCCAACATAGC
ACCCATCATCCTCCTCGTACTAGCAGCATGCGAGGCTGCCCTAGGCCTCTCTCTACTA
GTACTTGTATCTAACACATACGGCACGGACCACGTACAAAACCTAAACCTGCTACAA
TGCTAAAACTAATTGTCCCCACAGCCATACTAATCCCCCTAACATGACTATCGAACA
AAAACATAGTATGAATCAACGTAACATCACACAGCATGCTAATCAGCCTAACATCTC
TAGGCATCTTGGGCCAGCATGACCACAACAACATAAGCCTCTCAGCCATCTTCTTCT
CTGACCCCCTATCAGCACCCTTAATCGTATTAACAACATGACTCCTCCCCCTAATACT
AACAGCCAGCCAAGCCCACCTCTCCAATGAGCCCCTGGGCCTAAAAAAACTATACAT
TACCATACTAATCACCCTCCAAGCACTACTAATCATAACATTCGCCTCCTCTGAGTTC
ATTATATTCTACATCCTGTTTGAAGCAACACTCGTACCAACCCTAATCATCATCACAC
GATGAGGAAACCAAGCAGAGCGACTAAACGCAGGGTCCTACTTCCTCTTCTACACAA
TAGTAGGGTCACTCCCCCTTCTAGTAGTACTCACATACACCCAAAACATAACAGGAA
CCCTAAACATACTAGTACTACAATACTGAGCGAAACCCATAAACGACTCTTGATCCA
ACATGCTAATATGACTAGCATGCATAATAGCATTCATAGTAAAAATACCTCTATACG
GACTACACCTCTGACTGCCAAAAGCCCACGTAGAGGCCCCCATTGCAGGCTCCATAG
TACTTGCAGCCGTACTACTAAAACTAGGCGGATACGGCATAATACGCATCACTATCA
TGCTAGAACCAATAACAACATTCATAGCATACCCCTTCCTAATACTATCCCTGTGAG
GAATAATCATAACAAGCTCCATCTGCCTTCGCCAAACCGACCTAAAATCACTAATCG
CCTACTCCTCCGTAAGCCACATAGCACTGGTAATCGTCGCAATCCTAATCCAAACCC
CATGAAGCTACATAGGGGCCACCGCTCTCATAATCGCCCACGGTCTAACATCCTCCA
TACTATTTTGCCTAGCAAACACAAACTATGAGCGAACCCACAGCCGAACTATAATAC
TAGCACGAGGACTGCAAACCCTCCTACCCCTAATAGCTGCCTGATGGCTCCTAGCAA
GCCTAACCAACCTGGCCCTCCCCCCCAGCATCAACCTAATCGGAGAACTATTCGTAG
TAGTATCAACATTCTCGTGGTCTAACACCACCATCATCCTTACTGGAACAAACATTAT
CATCACAGCCACCTACTCCCTATACATACTAATATCCACCCAACGTGGAAAATACAC
CCACCACATCAACAACATCTGCCCCTCCTTCACCCGAGAAAACGCTCTAATGGCACT
CCACATACTGCCCCTACTAATGCTATCAACCAACCCCAAAATCATCCTAGGGTGCCT
TTACTGTAAGTATAGTTTAATAAAACCTCAGATTGTGGATCTGACAATAGAAGACCA

CAACTTCTTACTTACCAAAAAAGCATGCAAGAACTGCTAATTCCTGCCCCCATGTAT
AAAAACATGGCTTTTTAAACTTTTAGAGGATGACAGACACCCGTTGGTCTTAGGAA
CCAAAAGACTTGGTGCAACTCCAAGTAAAAGTAATTAACACCCTAACCTGCACCTCC
CTTATAACACTAACAACACTAATCATACCAATCATAATCACCCCCACAAATGCCTAC
ACAAGCAAAAACTACCCCCACCACGTAAAAAACATAGTCGCACTAGCCTTCGCCGTC
AGCATAATCCCAGCAA??????????????????????????????????????????GACACTGAACAAC
AATCCAAACAATAAAAATAACAATAAGCTTCAAACTAGACTATTTCTCCACAACATT
CATACCAGTGGCCCTATTCGTCACATGGTCTATCATAGAATTCTCTCTATGATATATA
AAATCAGACCCCCACATCAACCGATTCTTCAAGTACCTACTAATCTTCCTAATCACA
ATAATAATCCTAGTATCCGCCAACAACATATTCCAACTCTTCATCGGCTGAGAAGGA
GTAGGAATCATATCTTTCCTCCTAATCGGCTGATGACACGGACGAACAGATGCAAAC
ACAGCAGCCATACAGGCCATCCTATACAACCGCATCGGAGACATTGGACTAATCCTG
TCAATGGCATGATTCCTCACAAACCTAAACTCATGGGACCTCCAACAAATCCTCATG
CTAAACCCCGAGAACACAAACATCCCCCTAGCGGGCCTACTACTAGCAGCAACTGG
AAAATCCGCACAATTTGGGCTACACCCATGACTCCCCTCAGCCATAGAAGGCCCAAC
CCCAGTATCAGCCCTACTCCACTCCAGCACAATAGTAGTCGCAGGGGTATTCCTACT
AATCCGATTCCACCCCCTAATAGAAAACAACAAAACAATCCAAACAACAACCCTAT
GCCTAGGGGCCATTACAACCCTGTTCACAGCCATCTGTGCTCTCACCCAAAACGACA
TCAAAAAGATCGTGGCCTTCTCAACCTCAAGCCAACTAGGTCTAATAATAGTCACCA
TCGGAATCAACCAACCACACCTAGCCTTCATACACATCTGCACCCACGCCTTCTTCA
AAGCCATACTATTCATGTGCTCCGGATCGATCATCCACAGCCTAAACGACGAACAAG
ACATCCGCAAAATAGGAGGACTTTTCCACGCCCTCCCCACTACCACCTCCGCTCTCAT
TATTGGCAGCCTAGCCCTAACAGGAACCCCATTCCTAACAGGCTTCTACTCTAAAGA
CCTAATCATCGAAACCGCTAACATATCCCACACAAACGCCTGAGCCCTGCTCATCAC
ACTAGTAGCCACCTCCCTCACAGCCGCCTACAGCACACGAATCATCTTCTTCACACT
ATTAGGACAACCCCGATCCAACACCCTAATCAACATCAACGAAAACAACCCCTCACT
AACAAACCCAATCAAACGCCTAATGCTAGGGAGCATCTTCGCCGGATTCCTACTATA
CAACAACATCCCCCCAACAACCATGCCCCAAACAACTATACCACAATACCTAAAATA
CACCGCCCTCGCCACCACAATCCTAGGCCTAATCATAGCCCTAG??????????CCTGTCC
CTAAACCTAAGCCACAAACCCCCCTCAAGTGCCTTCAAATTCTCAACCTTACTAGGC
TACTTCCCCACAATCATTCACCGCTCAGGACCCCTCTGATCACTAACAACAAGCCAA
AAACTAGCATCCCTTATTCTAGACCTAATCTGACTAGAAAATATTATACCAAAGTCA
ATCTCACACTTCCACATAAAAGCCTCCCTAATAACATCCAACCAAAAAGGCTCAATC
AAACTATACTTCCTGTCCTTCGCAGTAACAATAATCCTCGCTCTCCTCATATTCTATT
CCCCCGGGTAACCTCCATAACAACAACAACACAAATAAACAAAGACCACCCGGTCA
TAATCACAAACCAAACACCATGGCTATAAAGAGCAGCAACACCCGCAGCTTCCTCAC
TAAAAAAACTAGACCCCCCAACATCATAAGTCGCCCAATCACCAGGACCACCAAGA
TCAAAAACCACACTCAACCCCTCACCCTTCAACATATAAAGCACTAAAACCAACTCT
ATGACCACACCCAAAACAAAAGCACCGAGCACAACCACATTAGACACCCAAACTTC
AGGATACTCCTCAATGGCTATGGCCGCCGTATAACCAAAAACAACCAACATCCCCCC
AAGATAAATCAAAAAAACCATTAAACCCAAAAAAGACCCACCAAAATTAACTACAA
CCCCACAACCTGCACCGCCAGCTACAACCAGACTAAACCCCCCATAAATAGGAGAA
GGCTTTGAAGAAAACCCTAAAAAACTAATCACAAAAATAACACTCAAAATAAACAC
AACATATGTCATTATTCCCACATGGACTCAAACCATGACCTATGACACGAAAAATCA
TCGTTGTAATTAAACTACGGAAACCTAATGACAAACATCCGAAAAACACACCCCCTA
CTCAAAATCATCAACAACTCCTTTATTGATCTCCCCACCCCCTCTAACATCTCAGCAT
GATGAAACTTCGGATCACTGCTAGGAATCTGCTTAGTCCTACAAATTCTCACAGGAC
TATTTCTAGCCATACACTACACAGCAGACACAGCAACCGCATTCTCATCAGTGACTC
ACATCTGCCGAGATGTAAACTACGGATGAATTATCCGATACATACACGCTAACGGAG
CTTCCCTGTTCTTCATCTGCCTGTTCGCACACATCGGACGAGGCATCTACTACGGATC
CTTTGCCTACAAAGAGACATGAAACATCGGTATCCTGCTCCTGTTTGCAGTAATAGC
AACAGCCTTTATGGGATACGTCCTACCATGAGGACAAATGTCCTTCTGAGGTGCTAC

AGTAATTACAAACCTTTTATCCGCAATACCCTACATCGGGTCTAGCCTAGTAGAGTG
AGTCTGAGGGGGATTCTCGGTAGACAAAGCAACTCTCACTCGATTCTTCGCTCTTCA
CTTCATCCTTCCCTTCGTAATTCTTGCCCTAGTACTAGTACACTTACTATTCTTACACG
AAACCGGATCCAACAACCCAATAGGAATCGTATCCAACCCCGACGTAATCCCCTTCC
ACCCATACTACACAGCCAAGGACACCCTTGGCCTATTCATCATGCTCACAGCACTAA
TATCCTTAGCCCTATTCTTCCCCGACCTACTAGGAGACCCAGACAATTATACACCCGC
AAACCCCCTAAACACACCACCCCACATCAAACCAGAATGGTACTTCCTATTCGCATA
CGCAATCCTACGCTCAATTCCCAACAAATTGGGAGGAGTACTAGCACTAATCCTATC
CATCCTCGTACTAGCACTAATCCCACTACTACACACATCAAAACAACGAACTATAAT
GTTCCGACCCCTGAGCCAAACCATCTTCTGACTCCTGGTGGCCGACCTACTAGTTCTC
ACATGAATCGGAGGACAACCCGTGGAACACCCCTTCATCCTAATCGGACAAGTGGCC
TCTATCCTTTATTTCACACTAATCCTAGCAGCAATGCCAATCGCAGGTATCATCGAAA
ACAATCTCATAAAATGAAGAGTCTTTGTAGTATATGTCAATACACTGGTCTTGTAAA
CCAGCAAAGGAATTAACCCTCCCCAAGACTCAGGAAGAAGACAAAAGCCCTACCAT
CAGCACCCAAAGCTGAAATTCTCAATAAACTACTTCCTGCAAACAATTAAAGTCCAA
CAAGCTTTATATAGTATATGCCATGTATAATCGTGCATTAATGGTTTGCCCCATGCAT
ATAAGCAGGTACATTATATTATTATAGTACATAGGACATATTATGTATAATCGTGCA
TTATTGATCTAGTACATACATATAAGCAGGTACATTATATTATTATAGTACATAGGAC
ATATTATGTATAATCGTGCATTATTGATCTAGTACATGCATATAAGCAGGTACATAAT
ATCCTTAGAGTACATAGTACATATTATTATTGATCGGACATAGCACATCAAGTCAAA
TCATTTCCAGTCAACATGCGTATCCCTACCACTGAAGGCCGCCTAATCACCATGCCG
CGTGAAATCATCAACCCGCTCATAGTCGTGTCCCTCTTCTCGCTCCGGGCCCATATGG
ACTGTGGGGTAGTTAAAGTGAAACTATACCTGGCATCTGGTTCTTACTTCATGTTCAT
CTTATATCTAGCCGCTCACTCGTTCCTCTTAAATAAGACATCTCGATGGATTAATTAC
TAATCAGCCCATGCCTAACATAACTGTGCTGTCATGCCTTTGGTATCTTTTAATTTTC
GGGGTGCGAGGTTCAACTGGGTCACCGACCTTCGAGCAGGTGGATAACTTGTAGATA
GACATTCATTGAATATTATTGGTCGTACATACTAACTCCTAGGTGTTATTCAGTCAAT
GGTTACAGGACATAGAGAATTTTACAACAAAATTTGAGTTGAGCCTACAGAAATCA
TAGATCACAATCGCAGACACCAACAAATACTAAAGTAATTAATTTAAATTTATAATT
TTAGGTTTGATATATCAAACCCCCCCTTACCCCCCCAAAGCCTCCACGTACTAAACAT
CTTGTCAAACCCCAAAAGCAAGAATACACACGTACACAAAGGCGCTATAAGCACAA
AGTATTACGTAAGTATATAGATATACAAAACTACTAGCGCAGCTTTCTGTCCACTAG
TTAAATACTTAGGTGCTTGAACATATATGGCTGGAAGGCCATGTAGTGACTACGTAA
TGATTTCTATTAGCCACAATAAAAACAATATACAATAAATACAAATATAGATCAATT
CTTATAGGCGCCGATAGCATAAACTATCTGCCCCCTATGTACAATTGAATTAAG

>Mus_musculus_GDP7L_mtDNA [consensus reconstructed from 1,634 reads]
?????????????????????????????????????????????????????????????????????????????GTCCTG
GCCTTATAATTAATTAGAGGTAAAATTACACATGCAAACCTCCATAGACCGGTGTAA
AATCCCTTAAACATTTACTTAAAATTTAAGGAGAGGGTATCAAGCACATTAAAATAG
CTTAAGACACCTTGCCTAGCCACACCCCCACGGGACTCAGCAGTGATAAATATTAAG
CAATAAACGAAAGTTTGACTAAGTTATACCTCTTAGGGTTGGTAAATTTCGTGCCAG
CCACCGCGGTCATACGATTAACCCAAACTAATTATCTTCGGCGTAAAACGTGTCAAC
TATAAATAAATAAATAGAATTAAAATCCAACTTATATGTGAAAATTCATTGTTAGGA
CCTAAACTCAATAACGAAAGTAATTCTAGTCATTTATAATACACGACAGCTAAGACC
CAAACTGGGATTAGATACCCCACTATGCTTAGCCATAAACCTAAATAATTAAATTTA
ACAAAACTATTTGCCAGAGAACTACTAGCCATAGCTTAAAACTCAAAGGACTTGGCG
GTACTTTATATCCATCTAGAGGAGCCTGTTCTATAATCGATAAACCCCGCTCTACCTC
ACCATCTCTTGCTAATTCAGCCTATATACCGCCATCTTCAGCAAACCCTAAAAAGGT
ATTAAAGTAAGCAAAAGAATCAAACATAAAAACGTTAGGTCAAGGTGTAGCCAATG
AAATGGGAAGAAATGGGCTACATTTTCTTATAAAAGAACATTACTATACCCTTTATG
AAACTAAAGGACTAAGGAGGATTTAGTAGTAAATTAAGAATAGAGAGCTTAATTGA

ATTGAGCAATGAAGTACGCACACACCGCCC??????????????????????????????????????????
????????????????????????????????????????????????????????????????????????????????????????
?????????????????????????????????????????????????????????????CCCTAGCCCTACACA
AATATAATTATACTATTATATAAATCAAAACATTTATCCTACTAAAAGTATTGGAGA
AAGAAATTCGTACATCTAGGAGCTATAGAACTAGTACCGCAAGGGAAAGATGAAAG
ACTAATTAAAAGTAAGAACAAGCAAAGATTAAACCTTGTACCTTTTGCATAATGAAC
TAACTAGAAAACTTCTAACTAAAAGAATTACAGCTAGAAACCCCGAAACCAAACGA
GCTACCTAAAAACAATTTTATGAATCAACTCGTCTATGTGGCAAAATAGTGAGAAGA
TTTTTAGGTAGAGGTGAAAAGCCTAACGAGCTTGGTGATAGCTGGTTACCCAAAAAA
TGAATTTAAGTTCAATTTTAAACTTGCTAAAAAAACAACAAAATCAAAAAGTAAGTT
TAGATTATAGCCAAAAGAGGGACAGCTCTTCTGGAACGGAAAAAACCTTTAATAGT
GAATAATTAACAAAACAGCTTTTAACCATTGTAGGCCTAAAAGCAGCCACCAATAAA
GAA-
AGCGTTCAAGCTCAACATAAAATTTCAATTAATTCCATAATTTACACCAACTTCCTAA
ACTTAAAATTG--
GGTTAATCTATAACTTTATAGATGCAACACTGTTAGTATGAGTAACAAGAATTCCAA
TTCTCCAGGCATACGCGTATAACAACTCGGATAACCATTGTTAGTTAATCAGACTAT
AGGCAATAATCACACTATAAATAATCCACCTATAACTTCTCTGTTAACCCAACACCG
GAATGCCTAAAGGAAAGATCCAAAAAGATAAAAGGAACTCGGCAAACAAGAACCC
CGCCTGTTTACCAAAAACATCACCTCTAGCATTACAAGTATTAGAGGCACTGCCTGC
CCAGTGACTAAAGTTTAACGGCCGCGGTATCCTGACCGTGCAAAGGTAGCATAATCA
CTTGTTCCTTAATTAGGGACTAGCATGAACGGCTAAACGAGGGTCCAACTGTCTCTT
ATCTTTAATCAGTGAAATTGACCTTTCAGTGAAGAGGCTGAAATATAATAATAAGAC
GAGAAGACCCTATGGAGCTTAAATTATATAACTTATCTATTTAATTTATTAAACCTAA
TGGCCCAAAAACTATAGTATAAGTTTGAAATTTCGGTTGGGGTGACCTCGGAGAATA
AAAAATCCTCCGAATGATTATAACCTAGACTTACAAGTCAAAGTAAAATCAACATAT
CTTATTGACCCAGATATATTTTGATCAACGGACCAAGTTACCCTAGGGATAACAGMG
CAATCCTATTTAAGAGTTCATATCGACAATTAGGGTTTACGACCTCGATGTTGGATCA
GGACATCCCAATGGTGTAGAAGCTATTAATGGTTCGTTTGTTCAACGATTAAAGTCC
TACGTGATCTGAGTTCAGACCGGAGCAATCCAGGTCGGTTTCTATCTATTTACGATTT
CTCCCAGTACGAAAGGACAAGAGAAATAGAGCCACCTTACAAATAAGCGC?????????
????????????????????????????????????????????????????????????????????????????????????????
??????????????????????????????????????????????????CTTTATTAATATCCTAACACTCCTC
GTCCCCATTCTAATCGCCATAGCCTTCCTAACATTAGTAGAACGCAAAATCTTAGGG
TACATACAACTACGAAAAGGCCCTAACATTGTTGGTCCATA-
CGGCATTTTACAACCATTTGCAGACGCCATAAAATTATTTATAAAAGAACCAATACG
CCCTTTAACAACCTCTATATCCTTATTTATTATTGCACCTACCCTATCACTCACACTAG
CATTAAGTCTATGAGTTCCCCTACCAATACCACACCCATTAATTAATTTAAACCTAGG
GATTTTATTTATTTTAGCAACATCTAGCCTATCAGTTTACTCCATTCTATGATCAGGA
TGAGCCTCAAACTCCAAATACTCACTATTCGGAGCTTTACGAGCCGTAGCCCAAACA
ATTTCATATGAAGTAACCATAGCTATTATC-
CTTTTATCAGTTCTATTAATAAATGGATCCTACTCTCTACAAACACTTATTACAACCC
AAGAACACATATGATTACTTCTGCCAGCCTGACCCATAGCCATAATATGATTTATCTC
AACCCTAGCAGAAACAAACCGGGCCCCCTTCGACCTGACAGAAGGAGAATCAGAAT
TAGTATCAGGGTTTAACGTAGAATACGCAGCCGGCCCATTCGCGTTATTCTTTATAGC
AGAGTACACTAACATTATTCTAATAAACGCCCTAACAACTATTATCTTCCTAGGACC
CCTATACTATATCAATTTACCAGAACTCTACTCAACTAACTTCATAATAGAAGCTCTA
CTACTATC-
ATCAACATTCCTATGGATCCGAGCATCTTATCCACGCTTCCGTTACGATCAACTTATA
CATCTTCTATGAAAAAACTTTCTACCCCTAACACTAGCATTATGTATGTGACATATTT
CTTTACCAATTTTTACAGCGGGAGTACCACC?????????????????????????????????????????
????????????????????????????????????????????????????????????????????????????????????????

??????????????????????????????????????TATCGGGCCCATACCCCGAAAACGTTGGTTTA
AATCCTTCCCGTACTAATAAATCCTATCACCCTTGCCATC-
ATCTACTTCACAATCTTCTTAGGTCCTGTAATCACAATATCCAGCACCAACCTAATAC
TAATATGAGTAGGCCTGGAATTCAGCCTACTAGCAATT--ATC-
CCCATACTAATCAACAAAAAAAA-
CCCACGATCAACTGAAGCAGCAACAAAATACTTCGTCACACAAGCAACAGCCTCAA
TAATTATCCTCCTGGCCATCGTACTCAACTATAAACAACTAGGAACATGAATATTTC
AACAACAAACAAACGGTCTTATCCTTAACATAACATTAATAGCCCTATCCATAAAAC
TAGGCCTCGCCCCATTCCACTTCTGATTACCAGAAGTAACTCAAGGGATCCCACTGC
ACATAGGACTTATTCTTCTTACATGACAAAAAATTGCTCCCCTATCAATTTTAATTCA
AATTTACCCGCTACTCAACTCTACTAT-
CATTTTAATACTAGCAATTACTTCTATTTTCATAGGGGCATGAGGAGGACTTAACCA
AACACAAATACGAAAAATTATAGCCTATTCATCAATTGCCCACATAGGATGAATATT
AGCAATTCTTCCTTACAACCCATCCCTCACTCTACTCAACCTCATAATCTATATTATT
CTTACAGCCCCTATATTCATAGCACTTATACTAAATAACTCTATAACCATCAACTCAA
TCTCACTTCTATGAAATAAAACTCCAGCAATACTAACTATAATCTCACTGATATTACT
ATCCCTAGGAGGCCTTCCACCACTAACAGGATTCTTACCAAAATGAATTATCATCAC
AGAACTTATAAAAAACAACTGTCTAATTATAGCAACACTCATAGCAATAATAGCTCT
ACTAAACCTATTCTTTTATACTCGCCTAATTTATTCCACTTCACTAACAATATTTCCAA
CCAACAATAACTCAAAAATAATAACTCACCAAACAAAAACTAAACCCAACCTAATA
TTTTCCA??????????????????????????????????????????????????????????????????????
?????????????????????????????????????????????????????????????????????????????????
??????????????????????????????????????CAGGAATTAAACCTACGAAAATTTAGTTAACAG
CTAAATACCCTATTACTGGCTTCAATCTACTTCTACCGCCGAAAAAAAAAAAATGGC
GGTAGAAGTCTTAGTAGAGATTTCTCTACACCTTCGAATTTGCAATTCGACATGAAT
ATCACC???????????????????????????????????????????????????????????????????????
???????????GTTGATTATTCTCAACCAATCACAAAGATATCGGAAC-
CCTCTATCTACTATTCGGAGCCTGAGCGGGAATAGTGGGTACTGCACTAAGTATTTT
AATTCGAGCAGAATTAGGTCAACCAGGTGCACTTTTAGGAGATGACCAAATTTACAA
TGTTATCGTAACTGCCCATG-
CTTTTGTTATAATTTTCTTCATAGTAATACCAATAATAATTGGAGGCTTTGGAAACTG
ACTTGTCCCACTAATAATCGGAGCCCCAGATATAGCATTCCCACGAATAAATAATAT
AAGTTTTTGACTCCTACCACCATCATTTCTCCTTCTCCTAGCATCATCAATAGTAGAA
GCAGGAGCAGGAACAGGATGAACAGTCTACCCACCTCT--
AGCCGGAAATCTAGCCCATGCAGGAGCATCAGTAGACCTAACAATTTTCTCCCTTCA
TTTAGCTGGAGTGTCATCTATTTTAGGTGCAATTAATTTTATTACCACTAT-
TATCAACATGAAACCCCCAGCCATAACACAGTATCAAACTCCACTATTTGTCTGATC
CGTACTTATTACAGCCGTACTGCTCCTATTATCACTACCAGTGCTAGCCGCAGGCATT
ACTATACTACTAACAGACCGCAACCTAAACACAACTTTCTTTGATCCCGCTGGAGGA
GGGGACCCAATTCTCTACCAGCATCTGTTCTGATTCTTTGGGCACCCAGAAGTTTATA
T-TCTTATCCTCCCAGG-
ATTTGGAATTATTTCACATGTAGTTACTTACTACTCCGGAAAAAAAGAACCTTTCGGC
TATATAGGAATAGTATGAGCAATAATGT-
CTATTGGCTTTCTAGGCTTTATTGTATGAGCCCACCACATATTCACAGTAGGATTAGA
TGTAGACACACGAGCTTACTTTACATCAGCCACTATAATTATCGCAATTCCTA---
CCGGTG--
TCAAAGTATTTAGCTGACTTGCAACCCTACACGGAGGTAATATTAAATGATCTCCAG
CTATACTATGAGCCTTAGGCTTTATTTTCTTATTTACAGTTGGTGGTCTAACCGGAAT
TGTTTTATCCAACTCATCCCTTGACATCGTGCT-TCACGAT-
ACATACTATGTAGTAGCCCATTTCCACTATGTTCTATCAATGGGAGCAGTGTTTGCTA
TCATAGCAGGATTTGTTCACTGATTCCCATTATTTCAGGCTT---CACCCTA-
GATGACACATGAGCAAAAGCCCACTTCGCCATCATATTCGTAGGAGTAAACATAACA

TTCTTCCCTCAACATTTCCTGGGCCTTTCAGGAATACCACGACGCTACTCAGACTACC
CAGATGCTTACACCACATGAAACACTGTCTCTTCT-ATA-
GGATCATTTATTTCACTAACAGCTGTTCTCATCATGATCTTTATAATTTGAGAGGCCT
TTGCTTCAAAACGAGAAGTAATATCAGTATCGTATGCTTCAACAAATTTAGAATGAC
TTCATGGCTGCCCTCCACCAT-ATCACACAT-TCG-
AGGAACCAACCTATGTAAAAGTAAAATAAGAAAGGAAGGAATCGAACCCCCTAAAA
TTGGTTTCAAGCCAATCTCATATCCTATA??????????????????????????????TTACATAA
CTTTGTCAAAGTTAAATTATAGATCAATAATCTATATATCTTATATGGCCTACCCATT
CCAACTTGGTCT-
ACAAGACGCCACATCCCCTATTATAGAAGAGCTAATAAATTTCCATGATCACACACT
AATAATTGTTTTCCTAATTAGCTCCTTAGTCCTCTATATCATCTCGCTAATATTAACA
ACAAAACTAAC-ACATACAAG-
CACAATAGATGCACAAGAAGTTGAAACCATTTGAACTATTCTACCAGCTGTAATCCT
TATCATAATTGCTCTCCCCTCTCTACGCATTCTATATATAATAGACGAAATCAACAAC
CCCGTATTAACCGTTAAAACCATAGGGCACCAATGATACTGAAGCTACGAATATACT
GACTATGAAGACCTATGCTTTGATTCATATATAATCCCAACAAACGACCTAAAACCT
GGTGAACTACGACTGCTAGAAGTTGATAACCGAGTCGTTCTGCCAATA-
GAACTTCCAATCCGTATATTAATTTCATCTGAAGACGTCCTCCACTCATGAGCAGTCC
CCTCCCTAGGACTTAAAACTGATGCCATCCCAGGCCGACTAAATCAAGCAACAGTAA
CATCAAACCGACCAGGGTTATTCTATGGCCAATGCTCTGAAATTTGTGGATCTAACC
ATAGCTTTATGCCCATTGTCCTAGAAATGGTTCCACTAAAATATTTCGAAAACTGATC
TGCTTC???????????????????????????????????????????????????????TAAAATCTCCAT
AGTGATATGCCACAACTAGATA-
CATCAACATGATTTATCACAATTATCTCATCAATAATTACCCTATTTATCTTATTTCA
ACTAAAAGTCTCATCACAAACATTCCCA---
CTGGCACCTTCACCAAAATCACTAACAACCATAAAAGTAAAAACCCCTTGAGAATTA
AAATGAACGAAAATCTATTTGCCTCATTCATTACCCCAACAATAATAGGATTCCCAA
TCGTTGTAGCCATCATTATATTTCCTTCAATCCTATTCCCATCCTCAAAACGCCTAAT
CAACAACCGTCTCCATTCTTTCCAACACTGACTAGTTAAACTTATTATCAAACAAATA
ATGCTAATCCACACACCAAAAGGACGAACATGAACCCTAATAATTGTTTCCCTAATC
ATATTTATTGGATCAACAAATCTCCTAGGCCTTTTACCACATACATTTACACCTACTA
CCCAACTATCCATAAATCTAAGTATAGCCATTCCACTATGAGCTGGAGCCGTAATTA
CAGGCTTCCGACACAAACTAAAAAGCTCACTTGCCCACTTCCTTCCA--
CAAGGAACTCCAATT-
TCACTAATTCCAATACTTATTATTATTGAAACAATTAGCCTATTTATTCAACCAATGG
CATTAGCAGTCCGGCTTACAGCTAACATTACTGCAGGACACTTATTA-
ATACACCTAATCGGAGGAGCTACTCTAGTATTAATAAATATTAGCCCACCAACAGCT
ACCATTACATTTATTATTTTACTTCTACTCACAATTCTAGAATTTGCAGTAGCATTAA
TTCAAGCCTACGTATTCACCCTCCTAGTAAGCCTATATCTACATKATAATACATAATG
ACCCACMAAACTCATGCATATCACATAGTTAATCCAAGTCCATGACCATTAACTGGA
GCCTTTTCAGCCCTCCTTCTAACATCAGGTCTAGTAATATGATTTCACTATAATTCAA
TTACACTATTAACCCTTGGCCTACTCACCAATATCCTCACAATATATCAATGATGACG
AGACGTAATTCGTGAAGGAACCTACCAAGGCCACCACACTCCTATTGTACAAAAAG
GACTACGATATGGTATAATTCTATTCATCGTCTCGGAAGTATTTTTCTTTGCAGGATT
CTTCTGAGCGTTCTATC-
ATTCTAGCCTCGTACCAACACATGATCTAGGAGGCTGCTGACCTCCAACAGGAATTT
CACCACTTAACCCTCTAGAAGTCCCACTACTTAATACTTCAGTACTTCTAGCATCAGG
TGTTTCAATTACATGAGCTCATCATAGCCTTATAGAAGGTAAACGAAACCACATAAA
TCAAGCCCTACTAATTACCATTATACTAGGACTTTACTTCACCATCCTCCAAGCTTCA
GAATACTTTGAAACATCATTCTCCATTTCAGATGGTATCTATGGTTCTACATTCTTC--
ATGGCTACTGGATTCCATGGACTCCATGTAATTATTGGATCAACATTCCTTATTGTTT
GCCTACTACGACAACTAAAATTTCACTTCACATCAAAACATCACTTCGGATTTGAAG

CCGCAGCATGATACTGACATTTTGTAGACGTAGTCTGACTTTTCCTATACGTC????????
???????????????????????????????????????????????????????????????????????????????????????????????????????????????
???????????????????????????????ATTTATTATCCCTAACGCTAATTCTAGTTGCATTCTGACTCC
CCCAAATAAATCTGTACTCAGAAAAAGCAAATCCATATGAATGCGGATTCGACCCTA
CAAGCTCTGCACGTCTACCATTCTCAATAAAATTTTTCTTGGTAGCAATTACATTTCT
ATTATTTGACCTAGAAATTGCTCTTCTACTTCCACTACCATGAGCAATTCAAACAATT
AAAACCTCTA???????????????????????????????????????????????????????????????????????????????????????????
???????????????????????????????????????????????????????????????????????????????????????????????????????????????
?????????????????????TCTTCAACCTCACCATAGCCTTCTCACTATCACTTCTAGGGACACTTA
TATTTCGCTCTCACCTAATATCCACATTACTATGCCTGGAAGGCATAGTATTATCCTT
ATTTATTATAACTTCAGTAACTTCCCTAAACTCCAACTCCATAAGCTCCATACCAATC
CCCATCACCATCTTAGTTTTCGCAGCCTGCGAAGCAGCTGTAGGACTAGCCCTACTA
GTAAAAGTTTCAAACACGTACGGAACAGATTACGTCCAAAATCTCAACCTACTACAA
TGCTAAAAATTATTCTTCCCTCACTAATGCTACTACCACTAACCTGACTATCAAGCCC
TAAAAAAACCTGAACAAACGTAACCTCATATAGTTTTCTAATTAGTTTAACCAGCCT
AACACTTCTATGACAAACCGACGAAAATTATAAAAACTTTTCAAATATATTCTCCTC
AGACCCCCTATCCACACCATTAATTATTTTAACAGCCTGATTACTGCCACTAATATTA
ATAGCTAGCCAAAACCACCTAAAAAAAGATAATAACGTACTACAAAAACTCTACAT
CTCAATACTAATCAGCTTACAAATTCTCCTAATCATAACCTTTTCAGCAACTGAACTA
ATTATATTTTATATTTTATTTGAAGCAACCTTAATCCCAACACTTATTATTATTACCCG
ATGAGGGAACCAAACTGAACGCCTAAACGCAGGGATTTATTTCCTATTTTATACCCT
AATCGGTTCTATTCCACTGCTAATTGCCCTCATCTTAATCCAAAACCATGTAGGAACC
CTAAACCTCATAATTTTATCATTCACAACACACACCTTAGACGCTTCATGATCTAACA
ACTTACTATGGTTGGCATGCATAATAGCATTTCTTATTAAAATACCATTATATGGAGT
TCACCTATGACTACCAAAAGCCCATGTTGAAGCTCCAATTGCTGGGTCAATAATTCT
AGCAGCTATTCTTCTAAAATTAGGTAGTTACGGAATAATTCGCATCTCCATTATTCTA
GACCCACTAACAAAATATATAGCATACCCCTTCATCCTTCTCTCCCTATGAGGAATA
ATTATAACTAGCTCAATCTGCTTACGCCAAACAGATTTAAAATCACTAATCGCCTACT
CCTCAGTTAGCCACATAGCACTTGTTATTGCATCAATCATAATCCAAACTCCATGAA
GCTTCATAGGAGCAACAATACTAATAATCGCACATGGCCTCACATCATCACTCCTAT
TCTGCCTAGCAAACTCCAACTACGAACGGATCCACAGCCGTACTATAATCATGGCCC
GAGGACTTCAAATGGTCTTCCCACTTATAGCCACAT-
GATGACTGATAGCAAGTCTAGCTAATCTAGCTCTACCCCCTTCAATCAATCTAATAG
GAGAATTATTCATTACCATATCATTATTTCTTGATCAAACTTTACCATTATTCTTATA
GGAATTAACATTATTATTACAGGTATATACTCAATATACATAATTATTACCACCCAAC
GCGGCAAACTAACCAACCATATAATTAACCTCCAACCCTCACA-
CACACGAGAACTAACACTAATAGCCCTTCACATAATTCCACTTATTCTTCTAACTACC
AGTCCA????????????????????????????????????????????????????????????????????????????????????????????????????
???????????????????????????????????????????????????????????????????????????????????????????????????????????????
??????????????????????????????????????????????????????????????????????????????????????????CTCAATCTTAT
TAATCTTCATTCTTCTACTATCCCCAATCCTAATTTCAATATCAAACCTAATTAAACA
CATCAACTTCCCACTGTACACCACCACATCAATCAAATTCTCCTTCATTATTAGCCTC
TTACCCCTATTAATATTTTTCCACAATAATATAGAATATATAATTACAACCTGGCACT
GAGTCACCATAAATTCAATAGAACTTAAAATAAGCTTCAAAACTGACTTTTTCTCTAT
CCTGTTTACATCTGTAGCCCTTTTTGTCACATGATCAATTATACAATTCTCTTCATGAT
ATATACACTCAGACCCAAACATCAATCGATTCATTAAATATCTTACACTATTCCTGAT
TACCATGCTTATCCTCACC??????????????????????????????????????????????????????????????????????????
??????????????????????GTACGGACGAACAGACGCAAATACTGCAGCCCTACAAGCAATCCT
CTATAACCGCATCGGAGACATCGGATTCATTTTAGCTATAGTTTGATTTTCCCTAAAC
ATAAACTCATGAGAACTTCAACAGATTATATTCTCCAACAACAACGACAATCTAATT
CCACTTATAGGCCTATTAATCGCAGCTACAGGAAAATCAGCACAATTTGGCCTCCAC
CCATGACTACCATCAGCAATAGAAGGCCCTACACCAGTTTCAGCACTACTACACTCA

AGTACAATAGTAGTTGCAGGAATTTTCCTACTGGTCCGATTCCACCCCCTCACGACTA
ATAATAACTTTATTTTAACAACTATACTTTGCCTCGGAGCCCTAACCACATTATTTAC
AGCTATTTGTGCTCTCACCCAAAACGACATCAAAAAAATCATTGCCTTCTCTACATCA
AGCCAACTAGGCCTGATAATAGTGACGCTAGGAATAAACCAACCACACCTAGCATTC
CTACACATCTGTACCCACGCATTCTTCAAAGCTATACTCTTTATATGCTCTGGCTCAA
TCATTCATAGCCTGGCAGACGAACAAGACATCCGAAAAATAGGAAACATCACAAAA
ATCATACCATTCACATCATCATGCCTAGTAATCGGAAGCCTCGCCCTCACAGGAATA
CCATTCCTAACAGGGTTCTACTCAAAAGACCTAATTATTGAAGCAATTAATACCTGC
AACACCAACGCCTGAGCCCTACTAATTACACTAATCGCCACTTCTATAACAGCTATG
TACAGCATACGAA-
TCATTTACTTCGTAACAATAACAAAACCGCGTTTTCCCCCCCTAATCTCCATTAACGA
AAATGACCCAGACCTCATAAACCCAATCAAACGCCTAGCATTCGGAAGCATCTTTGC
AGGATTTGTCATCTCATATAATATTCCACCAACCAGCATTCCAGTCCTCACAATACCA
TGATTTTTAAAAACCACAGCCCTAATTATTTCAGTATTAGGATTCCTAATCGCACTAG
AACTAAACAACCTAACCATAAAACTATCAATAAATAAAGCAAATCCATATTCATCCT
TCTCAACTTTACTGGGGTTTTTCCCATCTATTATTCACCGCATTACACCCATAAAATC
TCTCAACCTAAGCCTAAAAACATCCCTAACTCTCCTAGACTTGATCTGGTTAGAAAA
AACCATCCCAAAATCCACCTCAACTCTTCACACAAACATAACCACTTTAACAACCAA
CCAAAAAGGCTTAATTAAATTGTACTTTATATCATTCCTAATTAACATCATCTTAATT
ATTATCTTATACTCAATTAATCTCGAGTAATCTCGATAATAATAAAAATACCCGCAA
ACAAAGATCACCCAGCTACTACCATCATTCAAGTAGCACAACTATATATTGCCGCTA
CCCCAATCCCTCCTTCCAACATAACTCCAACATCATCAACCTCATACATCAACCAATC
TCCCAAACCATCAAGATTAATTACTCCAACTTCATCATAATAATTAAGCACACAAAT
TAAAAAAACCTCTATAATCACCCCCAATACTAAAAAACCCAAAATTAATCAGTTAGA
TCCCCAAGTCTCTGGATATTCCTCAGTAGCTATAGCAGTCGTATATCCAAACACAAC
CAACATCCCCCCTAAATAAATTAAAAAAACTATTAAACCTAAAAACGATCCACCAAA
CCCTAAAACCATTAAACAACCAACAAACCCACTAACAATTAAACCTAAACCTCCATA
AATAGGTGAAGGCTTTAATGCTAACCCAAGACAACCAACCAAAAATAATGAACTTA
AAACAAAAATATAATTATTCATTATTTCTACACAGCATTCAACTGCGACCAATGACA
TGAAAAATCATCGTTGTAATTCAACTA--
CAGAAACAC????????????????GAAAAACACACCCATTATTTAAAATTATTAACCACTC
ATTCATTGACCTACCTGCCCCATCCAACATTTCATCATGATGAAACTTTGGGTCCCTT
CTAGGAGTCTGCCTAATAGTCCAAATCATTACAGGTCTTTTCTTAGCCATACACTACA
CATCAGATACAATAACAGCCTTTTCATCAGTAACACACATTTGTCGAGACGTAAATT
ACGGGTGACTAATCCGATATATACACGCAAACGGAGCCTCAATATTTTTTATTTGCTT
ATTCCTTCATGTCGGACGAGGCTTATATTATGGATCATATACATTTATAGAAACCTGA
AACATTGGAGTACTTCTACTGTTCGCAGTCATAGCCACAGCATTTATAGGCTACGTCC
TTCCATGAGGACAAATATCATTCTGAGGTGCCACAGTTATTACAAACCTCCTATCAG
CCATCCCATATATTGGAACAACCCTAG-
TCGAATGAATTTGAGGGGGCTTCTCAGTAGACAAAGCCACCTTGACCCGATTCTTCG
CTTTCCACT-
TCATCTTACCATTTATTATCGCGGCCCTAGCAATCGTTCACCTCCTCTTCCTCC-
ACGAAACAGGATCAAACAACCCAACAGGATTAAACTCAGATGCAGATAAAATTCCA
TTTCACCCCTACTATACAATCAAAGATATCCTAGGTATCCTAATCATATTCTTAATTC
TCATAACCCTAGTATTATTTTTCCCAGACATACTAGGAGACCCAGACAACTACATAC
CAGCTAATCCACTA-AACACCC--
CACCCCATATTAAACCCGAATGATATTTCCTATTTGCATACGCCATTCTACGCTCAAT
CCCCAATAAACTAGGAGGTGTCCTAGCCTTA-
ATCTTATCTATCCTAATTTTAGCCCTAATACCTTTCCTTCATACCTCAAAGCAACGAA
GCCTAATATTCCGCCCAATCACACAAATTTTGTACTGAATCCTAGTAGCCAACCTACT
TATCTTAACCTGAATTGGGGGCCAACCAGTAGAACACCCATTTATTATCATTGGCCA
ACTAGCCTCCATCTCATACTTCTCAATCATCTTAATTCTTATACCAATC??????????????

?????????????????????????????????????????????????????????????????????????????????????????
?????????????????????????????????????????????????????????????????????????????????TTTAC
ATAGTACAACAGTACATTTATGTATATCGTACATTAAACTATTTTCCCCAAGCATATA
AGCTAGTACATTAAATCAATGKTTCAGGTCATAAAATAATCATCAACATAAATCAAT
ATATATACCATGAATATTATCTTAAACACATTAAACTAATKTTATAAGGACATATCT
GTGTTATCTGACATACACCATACAGTCATAAACTCTTCTCTTCCATATGACTAT??????
?????????????????????????????????????????????????????????????????????????????????????????
??GGGGTAGCTAAACTGAAACTTTATCAGACATCTGGTTCTTACTTCAGGGCCATCAA
ATGCGTTATCGCCCATACGTTCCCCTTAAATAAGACATCTCGATGGTATCGGGTCTAA
TCAGCCCATGACCAACATAACTGTGGTGTCATGCATTTGGTATCTTTTTATTTTGGCC
TACTTTCATCAACATAGCCGTCAAGGCATGAAAGGAC??????????????????????????????
?????????????????????????????????????????????????????????????????????????????????????????
?????????????????????????????????????????????????????????????????????????????????????????
?????????????????????????????????????????????????????????????????????????????????????????
????????????????????????????????????

**GD/P8L: *Manis javanica* + *Homo sapiens***

>Manis_javanica_GDP8L_mtDNA [consensus reconstructed from 7,727 reads]
?????????????????CCAAAGCAAAGCATTGAAAATGCTTAGATGAGTCCACAAACCTCC
ATAAACACACAGGTTTGGTCCCAGCCTTTTTATTAATTTTCAATAGGATTACACATGC
AAGTATCCGCCCCCCAGTGAAAATGCCCTTCAAGTTATCACAGTACCTGAAGGAGCT
GGCATCAAGCACGCCAATAAACGCAGCTAACGACGCCTTGCAGAGCCACACCCCCA
CGGGAAACAGCAGTGATAAAAATTAGGCTATAAACGAAAGTTCGACCTAGCCATAT
TGCATTGGGTTGGTAAATCTCGTGCCAGCCACCGCGGTCATACGATTAACCCTAGCT
AATAAAAAACCGGCGTAAAGCGTGCTAAGATGAATCCAATCCAAATAAAGTTAAGC
CCTGACCAGGCCGTAAAAAGCCGTGGTTACCGTAAAAATAAACTACGAAAGTAACT
TTAATTCACATCGACACACGATAGCTAAGACCCAAACTGGGATTAGATACCCCACTA
TGCTTAGCCCTAAACCTAAATAATTCACCAGACAAAATTATTCGCCAGAGAACTACT
GGCAACAGCCTAAAACTCAAAGGACTTGGCGGTGCTTTACATCCCCCTAGAGGAGCC
TGTTCTGTAATCGATAAACCCCGATAAACCCTACCAATTCTAGCTAATACAGCCTAT
ATACCGCCATCTCCAGCAAACCCTAAAAAGGAAGCACAGTAAGCAAGACTATGAAA
ACATAAATACGTTAGGTCAAGGTGTAGCCCATGAATTGGGAAGAAATGGGCTACATT
TTCTAAAATAGAACACAAACGAACGCCCTAATGAAAACGAGGGCCAAAGGAGGATT
TAGCAGTAAGCTGAGAATAGAAAGCTCAACTGAACCCGGCCCTAAAGCACGCACAC
ACCGCCCGTCACCCTCTTCAAATCCCCAAGAATACCTAAACATATTAGACAAACACC
AAGGCATGAGAAGAGACAAGTCGTAACAAGGTAAGCATACTGGAAGGTGTGCTTGG
ATCACCAAAGTGTAGCTTAAACAAAGCATCTGGCCTACACCCAGAAGATCTCAATAA
CATGACCACTTTGAACAAATCCTAGCCCAACCAACACCCAACATACAACCAAACAG
AGCACATAAACCAAAGCATTTACTAGACTAAAGTATAGGCGATAGAAATTACACAA
CGGCGCTATAGAAAAAGTACCGCAAGGGAAAGATGAAAGATGCATTCAAAGTACAA
AAAAGCAAAGATTACCCCTTGTACCTTTTGCATAATGGACTAACTAGAAACACCCTA
GCAAAGAGAACTTAAGCTAGAAACCCCGAAACCAGACGAGCTACCTACGAGCAGTT
TAAAGAACCCACTCATCTATGTGGCAAAATAGTGAGAAGACTTGTAGGTAGAGGTG
AAAAGCCTAACGAGCCTGGTGATAGCTGGTTGTCCAAGAAATGAATCTAAGTTCAAC
TTGAAGCATACCCAAAAGCCCAAAAACTATAATGTAGGCTTCAAGTATAGTCTAAAA
AGGTACAGCTTTTTAGAAACAGAATTAAATCTTAATTAGTGAGTAAACAATACAACA
ACCATAGTTGGCCTAAAAGCAGCCATCAATTAAGAAAGCGTCAAAGCTCAACAATA
AGACATAAATAATACCACAAATAAAAATCAACTCCTAAACCAATATTGGACCAATCT
ATCAATAAATAGAAGAAATACTGTTAGTATGAGTAACAAGAAATAGATCTCCTTGCA
CAAGCTTATATCAGAACGGATGACCCACTGATAATTAACAACGAAACAAATCAAAC
CCAAAAATAGAACTTTGTTAAATAGATTGTTAACCCAACACAGGCGTGCATATTCTC
AAAGGAAAGGTTAAAACAAATGAAAGGAACTCGGCAAACACAAGCCCCGCCTGTTT

ACCAAAAACATCACCTCTAGCATAACCAGTATTAGAGGCACTGCCTGCCCAGTGACT
CGCGTTAAACGGCCGCGKTWYTCTGACCGTGCAAAGGTAGCATAATCACTTGTTCTC
TAAATAAGGACTAGTATGAACGGCTAGACGAGGGTTTACTGTCTCTCATCTGTAAC
CAGTGAAATTGACCTCCCCGTGAAGAGGCGGGAATAATATAATAAGACGAGAAGAC
CCTATGGAGCTTTAATTAAACAGCTTAAAATTAACCCAACACCACTCTAACAGAGAT
ATAACAAACCAATTAGCCTAAGCTGTCAATTTTGGTTGGGGTGACCTCGGAGCAAAA
AACAACCTACGAGCGGTCAAATCCAGACTAACAAGTCCAGATAGTCCGTTAATTGAT
CCAACAACTTGATCAACGGAACAAGTTACCCTAGGGATAACAGCGCAATCCTATTCG
AGAGTCCATATCGACAATAGGGTTTACGACCTCGATGTTGGATCAGGACATCCCAAT
GGTGCAGAAGCTATTAATGGTTCGTTTGTTCAACGATTAAAGTCCTACGTGATCTGA
GTTCAGACCGGAGTAATCCAGGTCGGTTTCTATCTATTAATACACTTCTCCCAGTACG
AAAGGACAAGAGAAGTAGGGCCTACCTCACACAGGCGCCCTCGAATCAATAAATGA
CTACCCTCTTAATTTAGCCAATTCACAACAACCCCCAGTCCTAGAACCAGGGCCAGC
TAGGGTAGCAAAGCACGGCAACTGCACAAGACTTAAGCTCTTGTAACAGAGGTTCA
ACTCCCCTCCCTAGCAGCAATGTACACAATCAACATAATAATAATGAT?????????????
?????????????????????????????????????????????????????????????????????????????????????????????????
?????????????????????????????????????????????????????????????????????????????????????????????????
?????????????????????????????????????????????????????????????????????????????????????????????????
?????????????????????????????????????????????????????????????????????????????????????????????????
?????????????????????????????????????????????????????????????????????????????????????????????????
?????????????????????????????????????????????????????????????????????????????????????????????????
?????????????????????????????????????????????????????????????????????????????????????????????????
?????????????????????????????????????????????????????????????????????????????????????????????????
?????????????????????????????????????????????????????????????????????????????????????????????????
?????????????????????????????????????????????????????????????????????????????????????????????????
?????????????????????????????????????????????????????????????????????????????????????????????????
?????????????????????????????????????????????????????????????????????????????????????????????????
?????????????????????????????????????????????????????????????????????????????????????????????????
?????????????????????????????????????????????????????????????????????????????????????????????????
?????????????????????????????????????????????????????????????????????????????????????????????????
?????????????????????????????????????????????????????????????????????????????????????????????????
?????????????????????????????????????????????????????????????????????????????????????????????????
?????????????????????????????????????????????????????????????????????????????????????????????????
???????????????????????????????????????????????????CCAGACCCAACTACGAAAAATCATAG
CATACTCCTCAATCGCCCACATAGGCTGAATAGCAATAATCATAACATACAACCCGG
ACATTGCCATTCTAAACCTACTAGTCTACATCATAATGACACTATCTATATTCGCAAT
CCTACTATA????????????????????????????????????AAACAAGACACCACTAATCACA
GCCCTCGCACTACTAATCCTACTATCACTAGGAGGCCTCCCCCCACTGACCGGATTC
ATGCCCAAATGAATAATCATCCAAGAACTAACCAAAAACAACATAGTAATAATACC
AACCGTGATAGCAAT???????????????????????????????????????????????????????????????????????
?????????????????????????????????????????????????????????????????????????????????????????????????
?????????????????????????????????????????????????????????????????????????????????????????????????
?????????????????????????????????????????????????????????????????????????????????????????????????
?????????????????????????????????????????????????????????????????????????????????????????????????
?????????????????????????????????????????????????????????????????????????????????????????????????
?????????????????????????????????????????????????????????????????????????????????????????????????
?????????????????????????????????????????????????????CTGGGCCGGAATAGTAGGCACCGCA
CTAAGTCTTCTAATTCGCGCTGAGCTAGGTCAGCCTGGGACCCTCTTAGGGGACGAC
CAAATTTACAATGTGATCGTTACCGCACATGCATTTGTAATAATCTTCTTCATAGTCA
TGCCAATCATAATCGGGGGCTTTGGAAACTGACTAGTACCCCTGATAATTGGAGCCC
CTGACATGGCATTCCCCCGTATAAACAACATAAGTTTCTGACTGCTCCCCCCCTCTTT
CCTACTTCTTCTGGCCTCTTCCATAGTCGAAGCAGGGGCTGGAACCGGCTGAACTGT

ATACCCCCCTTTAGCAGGAAATTTAGCACACGCAGGAGCATCTGTAGACCTAACCAT
CTTCTCTCTTCATCTAGCGGGTGTCTCATCAATCCTTGGGGCTATCAATTTTATTACA
ACAATCATCAAC???????????????????????????????????????????????????????????????????????
??????????????????????????????????????????????????????CCGACCGTAATCTAAATACAACC
TTTTTTGACCCCGCAGGAGGAGGTGACCCCATTCTATACCAACACCTATTCTGATTCT
TCGGACACCCCGAAGTGTACATTCTCATCCTTCCTGGATTTGGAATAATCTCCCACAT
CGTAACCTATTACTCCGGGAAAAAAGAGCCCTTTGGATACATGGGCATAGTCTGAGC
AATAATATCCATTGGCTTCCTGGGCTTCATTGTATGGGCACACCACATGTTCACAGTG
GGAATAGACGTTGATACACGGGCCTACTTCACATCAGCCACCATAATTATTGCTATC
CCCACTGGAGTAAAGGTGTTTAGTTGACTAGCAACCCTGCACGGAGGAAACGTAAA
ATGGGCCCCAGCTATACTATGGGCCCTGGGCTTTATCTTCCTGTTTACAGTCGGGGGT
CTAACTGGCATCGTGCTGGCCAACTCGTCCCTAGATATCGTCCTCCATGATACCTACT
ACGTAGTAGCCCACTTCCACTACGTTCTCTCCATGGGAGCAGTTTTCGCCATCATAGG
AGGCTTCGTCCATTGATTCCCCCTGTTCTCAGGATACACGCTCAACAACACATGGGC
AAAAGTTCACTTTACAATCATATTCGTGGGCGTAAACATGACCTTTTTCCCCCAACAC
TTCCTTGGACTGTCTGGAATACCTCGACGATACTCGGACTACCCAGACGCTTACACA
ATGTGAAACACTGTGTCCTCCATAGGATCCTTCATCTCATTAACCGCCGTAATACTCA
TGGCCTTCATAATCTGAGAGGCATTCGCCTCTAAACGGGAAGTCCTAATAGTGGAGT
CCACAAATACCAACCTCGAATGACTACACGGCTGCCCACCACCCTACCACACATTCG
AAGAACCTGCTTTCGTAAATCTGGTCAAAACAAGAGAGGAAGGAATCGAACCCTCA
AACAATGGTTTCAAGCCAATATCATAGCCCCTATG???????????????????????????????????
???????????????????????????????????????????????????????????????????????????????????????????
???????????????????????????????????????????????????????????????????????????????????????????
???????????????????????????????????????????????????????????????????????????????????????????
???????????????????????????????????????????????????????????????????CCTGACAGTAAAAA
CAATGGGCCATCAATGATACTGAAGCTACGAATACACAGACTACGAAGACCTAAGC
TTCGACTCATACATAATCCCAACTCAAGACCTAAAACCCGGCGAACTCCGACTTCTA
GAAGTGGACAACCGACTTGTACTACCCATAGACACAACCATCCGCATGCTCATCTCA
TCTGAAGACGTCCTGCACTCTTGAGCCRTCCCATCCCTGGGCCTAAAAACAGATGCT
ATCCCGGGGCGTCTAAACCAAACAACCCTGAT???????????????????????????????????????
???????????????????????????????????????????????????????????????????????????????????????????
???????????????????????????????????????????????????????????????????????????????????????????
???????????????????????????????????????????????????????????????????????????????????????????
???????????????????????????????????????????????????????????????????????????????????????????
?????????????????TCCCTATTGTAACAATTATCATTATGTTCCCAGTAATCCTCTTCCCA
ACATCAAACCGACTAATCAACAACCGCATTGTATCCATACAACAATGACTCCTAAAA
CAAACATCCAAACAAATAATAAGCATCCACAACTACAAAGGACAGACCTGAACCCT
GATATTAATAACACTAATCATTTTCATCGCATCTACTAACCTACTAGGCCTGCTACCC
CACTCATTCACCCCCACAGCCCAACTGTCAATAAACCTGAGCATAGCCGTTCCTCTAT
GGGCAGCCACCGTAGTCACAGGTT?????????????????????????????????????????????????????
???????????????????????????????????????????????????????????????????????????????????????????
???????????????????????????????????????????????????????????????????????????????????????????
???????????????????????????????????????????????????????????????????????????????????????????
???????????????????????????????????????????????????????????????????????????????????????????
??????????????GGGCCTAGCAATATGATTCCACTTCAACTCTACAACACTACTCCTCCT
AGGACTAACAACAAACCTCTTAACAATATATCAATGWTGACGTGACATTGTACGAG
AAAGCACCTTTCAAGGCCACCACACACCCACAGTCCAAAAAGGACTACGATAC?????
????????????????????????TATTTTTCTTCGCTGGCTTCTTCTGAGCATTCTACCACTCAAG
CCTAGCACCTACCCCCGAACTAGGAGGATGCTGACCCCCCACAGGCATCAACCCGCT
AAACCCACTGGAAGTACCTCTACTCAACACATCCGTTCTTTTAGCCTCAGGAGTATC
AATTACATGGGCACACCATAGCTTAATAGAAGGCAGCCGAAACCACATAATCCAGG
CCCTGCTCATCACAATYCTCCTAGGCATTTACTTCACACTACTGCAGGTCTCAGAATA

TTACGAAGCACCCTTCACAATCTC???????????????????????????????????????????????????
????????????????????????????????????????????????????????????????????????????????????????
????????????????????????????????????????????????????????????????????????????????????????
????????????????????????????????????????????????????????????????????????????????????????
????????????????????????????????????????????????????????????????????????????????????????
???????????????GGGATCGGCACGACTACCATTCTCCATAAAATTCTTCCTAATCGCCAT
CACGTTCCTACTATTCGACCTAGAAATCGCACTACTCCTG???????????????????????????
????????????????????????????????????????????????????????????????????????????????????????
????????????????????????????????????????????????????????????????????????????????????????
????????????????????????????????????????????????????????????????????????????????????????
????????????????????????????????????????????????????????????????????????????????????????
????????????????????????????????????????????????????????????????????????????????????????
????????????????????????????????????????????????????????????????????????????????????????
????????????????????????????????????????????????????????????????????????????????????????
????????????????????????????????????????????????????????????????????????????????????????
????????????????????????????????????????????????????????????????????????????????????????
??????????????????????????????????????????????????????????????????????ATGAGGAAACC
AAGCAGAGCGACTAAACGCAGGGTCCTACTTCCTCTTCTACACAATAGTAGGGTCAC
TCCCCCTTCTAGTAGTACTCACATACACCCAAAACATAACAGGAACCCTAAACATAC
TAGTACTACAATACTGAGCGAAACC?????????????????????????????????????????????????
????????????????????????????????????????????????????????????????????????????????????????
????????????????????????????????????????????????????????????????????????????????????????
????????????????????????????????????????????????????????????????????????????????????????
????????????????????????????????????????????????????????????????????????????????????????
????????????????????????????????????????????????????????????????????????????????????????
????????????????????????????????????????????????????????????????????????????????????????
????????????????????????????????????????????????????????????????????????????????????????
????????????????????????????????????????????????????????????????????????????????????????
????????????????????????????????????????????????????????????????????????????????????????
????????????????????????????????????????????????????????????????????????????????????????
????????????????????????????????????????????????????????????????????????????????????????
????????????????????????????????????????????????????????????????????????????????????????
????????????????????????????????????????????????????????????????????????????????????????
????????????????????????????????????????????????????????????????????????????????????????
????????????????????????????????????????????????????????????????????????????????????????
????????????????????????????????????????????????????????????????????????????????????????
????????????????????????????????????????????????????????????????????????????????????????
????????????????????????????????????????????????????????????????????????????????????????
????????????????????????????????????????????????????????????????????????????????????????
????????????????????????????????????????????????????????????????????????????????????????
????????????????????????????????????????????????????????????????????????????????????????
???????????????????????????????????????????????????????????CTAATAATAGTCACCATCG
GAATCAACCAACCACACCTAGCCTTCATACACATCTGCACCCAYGCCTTTTTCAAAG
CCATACTATTCATGTGCTCCGGATCGATCATCCACAGCCTAAACGACGAACAAGACA
TCCGCAAAATAGGAGG?????????????????????????????????????????????????????????????G
GAACCCCATTCCTAACAGGCTTCTACTCTAAAGACCTAATCATCGAAACCGCTAACA
TATCCCACACAAACGCCTGAGCCCTGCTCATCWCACTAGTAGCCACMTCCCTCACAG
CCGCCTACAGCACACGAATCATCTTCTTCACACTA???????????????????????????????????
????????????????????????????????????????????????????????????????????????????????????????
????????????????????????????????????????????????????????????????????????????????????????
????????????????????????????????????????????????????????????????????????????????????????
????????????????????????????????????????????????????????????????????????????????????????

?????????????????????????????????????????????????????????????????????????????????????????
?????????????????????????????????????????????????????????????????????????????????????????
?????????????????????????????????????????????????????????????????????????????????????????
?????????????????????????????????????????????????????????????????????????????????????????
?????????????????????????????????????????????????????????????????????????????????????????
?????????????????????????????????????????????????????????????????????????????????????????
?????????????????????????????????????????????????????????????????????????????????????????
?????????????????????????????????????????????????????????????????????????????????????????
?????????????????????????????????????????????????????????????????????????????????????????
?????????????????????????????????????????????????????????????????????????????????????????
??????????????????????????????????????TAACTCACATCTGCCGAGATGTAAACTACGGAT
GAATTATCCGATACATACACGCTAACGGAGCTTCCCTGTTCTTCATCTGCCTGTTCGC
ACACATCGGACGAGGCATCTACTACGGATCCTTTGCCTACAAAGAGACATGAAACAT
CGGTATCCTGCTCCTGTTTGCAGTAATAGCAACAGCCTTTATGGGATACGTCCTACCA
TGAGGACAAATGTCCTTCTGAGGTGCTACGGTAATTACAAACCTTTTATCCGCAATA
CCCTACATCGGGTCTAGCCTAGTAGAGTGAGTCTGAGGGGGATTCTCGGTAGACAAA
GCAACTCTCACTCGATTCTTCGCTCTTCACTTCATCCTTCCCTTCGTAATTCTTGCCCT
AGTACTAGTACACTTACTATTCTTACACGAAACCGGATCCAACAACCCAATAGGAAT
CGTATCCAACCCCGACGTAATCCCCTTCCACCCATACTACACAGCCAAGGACACCCT
TGGCCTATTCATCATGCTCACAGCACTAATATCCTTAGCCCTATTCTTCCCCGACCTA
CTAGGAGACCCAGACAATTATACAYCCGCAAACCCC?????????????????????????????????
?????????????????????????????????????????????????????????????????????????????????????????
???????TCCCACTACTACACACATCAAAACAACGAACTATAATGTTCCGACCCCTGAG
CCAAACCATCYTCTGACTCCTGGTGGCYGACCTACTAGTTCTCACATGAATCGGAGG
ACAACCCGTGGAACACCCCTTCATCCTAATCGGACAAGTRGCCTCTATCCTTTATTTC
ACACTAATCCTAGCAGCAATGCCAATCGCAGGTATCATCGAAAACAATCTCATAAAA
TGAAGAGTCTTTGTAGTATATGTCAATACACTGGTCTTGTAAACCAGCAAAGGAATT
AACCCTCCCCAAGACTCAGGAAGAAGACAAAAGCCCTACCATCAGCACCCAAAGCT
GAAATTCTCAATAAACTACTTCCTGCAA??????????????????????????????????????????????
?????????????????????????????????????????????????????????????????????????????????????????
?????????????????????????????????????????????????????????????????????????????????????????
?????????????????????????????????????????????????????????????????????????????????????????
?????????????????????????????????????????????????????????????????????????????????????????
?????????????????????????????????????????????????????????????????????????????????????????
?????????????????????????????????????????????????????????????????????????????????????????
????????????AATTTTCGGGGTGCGAGGTTCAACTGGGTCACCGACCTTCGAGCAGGTK
GATAACTTGTAGATAGACATTCATTGAATATTATTGGTCGTACATACTAACTCCTAGG
TGTTATTCAGTCAATGGTTACAGGACATAGAGAATTTACAAC??????????????????????
?????????????????????????????????????????????????????????????????????????????????????????
?????????????????????????????????????????????????????????????????????????????????????????
?????????????????????????????????????????????????????????????????????????????????????????
?????????????????????????????????????????????????????????????????????????????????????????
???????????????????????????????????????????????????

>Homo_sapiens_GDP8L_mtDNA [consensus reconstructed from 1,333 reads]
??????????????CACCCTATTAACCACTCACGGGAGCTCTCCATGCATTTGGTATTTTCGT
CTGGGGGGTGTGCACGCGATAGCATTGCGAGACGCTGGAGCCGGAGCACCCTATGT
CGCAGTATCTGTCTTTGATTCCTGCCTCATTCTATTATTTATCGCACCTACGTTCAATA
TTACAGGCGAACATACTTACCAAAGTGTGTTAATTAATTAATGCTTGTAGGACATAA
TAATAACAATTGAATGTCTGCACASCCGCTTTCCACACAGACATCATAACAAAAAAT
TT???????????????????????????????????????????????????????????????????????????????????????
?????????????????????????????????????????????????????????????????????????????????????????

????????????????????????????????????????????????????????????????????????????????????????????
????????????????????????????????????????????????????????????????????????????????????????????
???????????????CCTAGCCTTTCTATTAGCTCTTAGTAAGATTACACATGCAAGCATCC
CCGTTCCAGTGAGTTCACCCTCTAAATCACCACGATCAAAAGGACAAGCATCAAGC
ACGCAGCAATGCAGCTCAAAACGCTTAGCCTAGCCACACCCCCACGGGAAACAGCA
GTGATTAACCTTTAGCAATAAACGAAAGTTTAACTAAGCTATACTAACCCCAGGGTT
GGTCAATTTCGTGCCAGCCACCGCGGTCACACGATTAACCCAAGTCAATAGAAGCCG
GCGTAAAGAGTGTTTTAGATCACCCCCTCCCCAATAAAGCTAAAACTCACCTGAGTT
GTAAAAAACTCCAGTTGACACAAAATAGACTACGAAAGTGGCTTTAACATATCTGAA
CACACAATAGCTAAGACCCAAACTGGGATTAGATACCCCACTATGCTTAGCCCTAAA
CCTCAACAGTTAAATCAACAAAACTGCTCGCCAGAACACTACGAGCCACAGCTTAAA
ACTCAAAGGACCTGGCGGTGCTTCATATCCCTCTAGAGGAGCCTGTTCTGTAATCGA
TAAACCCCGATCAACCTCACCACCTCTTGCTCAGCCTATATACCGCCATCTTCAGCAA
ACCCTGATGAAGGCTACAAAGTAAGCGCAAGTACCCACGTAAAGACGTTAGGTCAA
GGTGTAGCCCATGAGGTGGCAAGAAATGG????????????????????????????????????????????
????????????????????????????????????????????????????????????????????????????????????????????
????????????????????????????????????????????????????????????????????????????????????????????
????????????????????????????????????????????????????????????????????????????????????????????
??????????????CCCCAAACCCACCCCACCTTACTACCAGACAACCTTAGCCAAACCATT
TACCCAAATAAAGTATAGGCGATAGAAATTGAAACCTGGCGCAATAGATATAGTAC
CGCAAGGGAAAGATGAAAAATTATAACCAAGCATAATATAGCAAGGACTAACCCCT
ATACCTTCTGCATAATGAATTAACTAGAAATAACTTTGCAAGGAGAGCCAAAGCTAA
GACCCCCGAAACCAGACGAGCTACCTAAGAACAGCTAAAAGAGCACACCCGTCTAT
GTAGCAAAATAGTGGGAAGATTTATAGGTAGAGGCGACAAACCTACCGAGCCTGGT
GATAGCTGGTTGTCCAAGATAGAATCTTAGTTCAACTTTAAATTTGCCCACAGAACC
CTCTAAATCCCCTTGTAAATTTAACTGTTAGTCCAAAGAGGAACAGCTCTTTGGACA
CTAGGAAAAAACCTTGTAGAGAGAGTAAAAAATTTAACACCCATAGTAGGCCTAAA
AGCAGCCACCAATTAAGAAAGCGTTCAAGCTCAACACCCACTACCTAAAAAATCCC
AAACATATAACTGAACTCCTCACACCCAATTGGACCAATCTATCACCCTATAGAAGA
ACTAATGTTAGTATAAGTAACATGAAAACATTCTCCTCCGCATAAGCCTGCGTCAGA
TTAAAACACTGAACTGACAATTAACAGCCCAATATCTACAATCAACCAACAAGTCAT
TATTACCCTCACTGTCAACCCAACACAGGCATGCTCATAAGGAAAGGTTAAAAAAAG
TAAAAGGAACTCGGCAAATCTTACCCCGCCTGTTTACCAAAAACATCACCTCTAGCA
TCACCAGTATTAGAGGCACCGCCTGCCCAGTGACACATGTTTAACGGCCGCGGTACC
CTAACCGTGCAAAGGTAGCATAATCACTTGTTCCTTAAWTAGGGACCTGTATGAATG
GCTCCACGAGGGTTCAGCTGTCTCTTACTTTTAACCAGTGAAATTGACCTGCCCGTGA
AGAGGCGGGCATGACACAGCAAGACGAGAAGACCCTATGGAGCTTTAATTTATTAA
TGCAAACAGTACCTAACAAACCCACAGGTCCTAAACTACCAAACCTGCATTAAAAAT
TTCGGTTGGGGCGACCTCGGAGCAGAACCCAACCTCCGAGCAGTACATGCTAAGACT
TCACCAGTCAAAGCGAACTACTATACTCAATTGATCCAATAACTTGACCAACGGAAC
AAGTTACCCTAGGGATAACAGCGCAATCCTATTCTAGAGTCCATATCAACAATAGGG
TTTACGACCTCGATGTTGGATCAGGACATCCCGATGGTGCAGCCGCTATTAAAGGTT
CGTTTGTTCAACGATTAAAGTCCTACGTGATCTGAGTTCAGACCGGAGTAATCCAGG
TCGGTTTCTATCTACTTCAAATTCCTCCCTGTACGAAAGGACAAGAGAAATAAGGCC
TACTTCACAAAGCGCCTTCCCCCGTAAATGATATCATCTCAACTTAGTATTATACCCA
CACCCACCCAAGAACAGGGTTTGTTAAGATGGCAGAGCCCGGTAATCGCATAAAAC
TTAAAACTTTACAGTCAGAGGTTCAATTCCTCTTCTTAACAACATACCCATGGCCAAC
CTCCTACTCCTCATTGTACCCATTCTAATCGCAATGGCATTCCTAATGCTTACCGAAC
GAAAAATTCTAGGCTATATACAACTACGCAAAGGCCCCAACGTTGTAGGCCCCTACG
GGCTACTACAACCCTTCGCTGACGCCATAAAACTCTTCACCAAAGAGCCCCTAAAAC
CCGCCACATCTACCATCACCCTCTACATCACCGCCCCGACCTTAGCTCTCACCATCGC
TCTTCTACTATGAACCCCCCTCCCCATACCCAACCCCCTGGTCAACCTCAACCTAGGC

CTCCTATTTATTCTAGCCACCTCTAGCCTAGCCGTTTACTCAATCCTCTGATCAGGGT
GAGCATCAAACTCAAACTACGCCCTGATCGGCGCACTGCGAGCAGTAGCCCAAACA
ATCTCATATGAAGTCACCCTAGCCATCATTCTACTATCAACATTACTAATAAGTGGCT
CCTTTAACCTCTCCACCCTTATCACAACACAAGAACACCTCTGATTACTCCTGCCATC
ATGACCCTTGGCCATAATATGATTTATCTCCACACTAGCAGAGACCAACCGAACCCC
CTTCGACCTTGCCGAAGGGGAGTCCGAACTAGTCTCAGGCTTCAACATCGAATACGC
CGCAGGCCCCTTCGCCCTATTCTTCATAGCCGAATACACAAACATTATTATAATAAA
CACCCTCACCACTACAATCTTCCTAGGAACAACATATAACGCACTCTCCCCTGAACT
CTATACAACATATTTTGTCACCAAGACCCTACTTCTAACCTCCCTGTTCTTATGAATT
CGAACAGCATACCCCCGATTCCGCTACGACCAACTCATRCACCTCCTATGAAAAAAC
TTCCTACCACTCACCCTAGCATTACTTATATGATATGTCTCCATACCCATTACAATCT
CCAGCATTCCCCC??????????????????????TAAAAGAGTTACTTTGATAGAGTAAATAA
TAGGAGCTTAAACCCCCTTATTTCTAGGACTATGAGAATCGAACCCATCCCTGAGAA
TCCAAAATTCTCCGTGCCACCTATCACACCCCATCCTAAAGTAAGGTCAGCTAAATA
AGCTATC???????????????????TTGGTTATACCCTTCCCGTACTAATTAATCCCCTGGCC
CAACCCGTCATCTACTCTACCATCTTTGCAGGCACACTCATCACAGCGCTAAGCTCGC
ACTGATTTTTTACCTGAGTAGGCCTAGAAATAAACATGCTAGCTTTTATTCCAGTTCT
AACCAAAAAAATAAACCCTCGTTCCACAGAAGCTGCCATCAAGTATTTCCTCACGCA
AGCAACCGCATCCATAATCCTTCTAATAGCTATCCTCTTCAACAATATACTCTCCGGA
CAATGAACCATAACCAATACTACCAATCAATACTCATCATTAATAATCATAATGGCT
ATAGCAATAAAACTAGGAATAGCCCCCTTTCACTTCTGAGTCCCAGAGGTTACCCAA
GGCACCCCTCTGACATCCGGCCTGCTTCTTCTCACATGACAAAAACTAGCCCCCATCT
CAATCATATACCAAATCTCTCCCTCACTAAACGTAAGCCTTCTCCTCACTCTCTCAAT
CTTATCCATCATAGCAGGCAGTTGAGGTGGATTAAACCAAACCCAGCTACGCAAAAT
CTTAGCATACTCCTCAATTACCCACATAGGATGAATAATAGCAGTTCTACCGTACAA
CCCTAACATAACCATTCTTAATTTAACTATTTATATTATCCTAACTACTACCGCATTC
CTACTACTCAACTTAAACTCCAGCACCACGACCCTACTACTATCTCGCACCTGAAAC
AAGCTAACATGACTAACACCCTTAATTCCATCCACCCTCCTCTCCCTAGGAGGCCTGC
CCCCGCTAACCGGCTTTTTGCCCAAATGGGCCATTATCGAAGAATTCACAAAAAACA
ATAGCCTCATCATCCCCACCATCATAGCCACCATCACCCTCCTTAACCTCTACTTCTA
CCTGCGCCTAATCTACTCCACCTCAATCACACTACTCCCCATATCTAACAACGTAAAA
ATAAAATGACAGTTTGAACATACAAAACCCACCCCATTCCTCCCCACACTCATCACC
CTTACCACGCTACTCCTACCTATCYCCCCTTTTATACTAATAATCTTAT???????????????
?????????????????????????????????????????????????????????GGACTGCAAAACCCCA
CTCTGCATCAACTGAACGCAAATCAGCCACTTTAATTAAGCTAAGCCCTTACTAGAC
CAATGGGACTTAAACCCACAAACACTTAGTTAACAGCTAAGCACCCTAATCAACTGG
CTTCAATCTACTTCTCCCGCCGCCGGGAAAAAAGGCGGGAGAAGCCCCGGCAGGTTT
GAAGCTGCTTCTTCGAATTTGCAATTCAATATGAAAATCACCTCGGAGCTGGTAAAA
AGAGGCCTAACCCCTGTCTTTAGATTTACAGTCCAATGCTTCACTCAGCCATTTTACC
TCACCCCSACTGATGTTCGCCGACCGTTGACTATTCTCTACAAACCACAAAGACATTG
GAACACTATACCTATTATTCGGCGCATGAGCTGGAGTCCTAGGCACAGCTCTAAGCC
TCCTTATTCGAGCCGAGCTGGGCCAGCCAGGCAACCTTCTAGGTAACGACCACATCT
ACAACGTTATCGTCACAGCCCATGCATTTGTAATAATCTTCTTCATAGTAATACCCAT
CATAATCGGAGGCTTTGGCAACTGACTAGTTCCCCTAATAATCGGTGCCCCCGATAT
GGCGTTTCCCCGCATAAACAACATAAGCTTCTGACTCTTACCTCCCTCTCTCCTACTC
CTGCTCGCATCTGCTATAGTGGAGGCCGGAGCAGGAACAGGTTGAACAGTCTACCCT
CCCTTAGCAGGGAACTACTCCCACCCTGGAGCCTCCGTAGACCTAACCATCTTCTCCT
TACACCTAGCAGGTGTCTCCTCTATCTTAGGGGCCATCAATTTCATCACAACAATTAT
CAATATAAAACCCCCTGCCATAACCCAATACCAAACGCCCCTCTTTGTCTGATCCGTC
CTAATCACAGCAGTCCTACTTCTCCTATCTCTCCCAGTCCTAGCTGCTGGCATCACTA
TACTACTAACAGACCGCAACCTCAACACCACCTTCTTCGACCCCGCCGGAGGAGGAG
ACCCCATTCTATACCAACACCTATTCTGATTTTTCGGTCACCCTGAAGTTTACATTCTT

ATCCTACCAGGCTTCGGAATAATCTCCCATATTGTAACCTACTACTCCGGAAAAAAA
GAACCATTTGGATACATAGGTATGGTCTGAGCTATGATATCAATTGGCTTCCTAGGG
TTTATCGTGTGAGCACACCATATATTTACAGTAGGAATAGACGTAGACACACGAGCA
TATTTCACCTCCGCTACCATAATCATCGCTATCCCCACCGGCGTCAAAGTATTTAGCT
GACTCGCCACACTCCACGGAAGCAATATGAAATGATCTGCTGCAGTGCTCTGAGCCC
TAGGATTCATCTTTCTTTTCACCGTAGGTGGCCTGACTGGCATTGTATTAGCAAACTC
ATCACTAGACATCGTACTACACGACACGTACTACGTTGTAGCTCACTTCCACTATGTC
CTATCAATAGGAGCTGTATTTGCCATCATAGGAGGCTTCATTCACTGATTTCCCCTAT
TCTCAGGCTACACCCTAGACCAAACCTACGCCAAAATCCATTTCACTATCATATTCAT
CGGCGTAAATCTAACTTTCTTCCCACAACACTTTCTCGGCCTATCCGGAATGCCCCGA
CGTTACTCGGACTACCCCGATGCATACACCACATGAAACATCCTATCATCTGTAGGC
TCATTCATTTCTCTAACAGCAGTAATATTAATAATTTTCATGATTTGAGAAGCCTTCG
CTTCGAAGCGAAAAGTCCTAATAGTAGAAGAACCCTCCATAAACCTGGAGTGACTAT
ATGGATGCCCCCCACCCTACCACACATTCGAAGAACCCGTATACATAAAATCTAGAC
AAAAAAGGAAGGAATCGAACCCCCCAAAGCTGGTTTCAAGCCAACCCCATGGCCTC
CAT??????????????????????????????????????????????????????????????????????????????
??????????????GTAGGTCTACAAGACGCTACTTCCCCTATCATAGAAGAGCTTATCAC
CTTTCATGATCACGCCCTCATAATCATTTTCCTCATCTGCTTCCTAGTCCTGTATGCCC
TTTTCCTAACACTCACAACAAAACTAACTAATACTAACATCTCAGACGCTCAGGAAA
TAGAAACCGTCTGAACTATCCTGCCCGCCATCATCCTAGTCCTCATCGCCCTCCCATC
CCTACGCATCCTTTACATAACAGACGAGATCAACGATCCCTCCCTTACCATCAAATC
AATTGGCCACCAATGGTACTGAACCTACGAGTACACCGACTACGGCGGACTAATCTT
CAACTCCTACATACTTCCCCCATTATTCCTAGAACCAGGCGACCTGCGACTCCTTGAC
GTTGACAATCGAGTAGTACTCCCGATTGAAGCCCCCATTCGTATAATAATTACATCA
CAAGACGTCTTGCACTCATGAGCTGTCCCCACATTAGGCTTAAAAACAGATGCAATT
CCCGGACGTCTAAACCAAACCACTTTCACCGCTACACGACCGGGGGTATACTACGGT
CAATGCTCTGAAATCTGTGGAGCAAACCACAGTTTCATGCCCATCGTCCTAGAATTA
ATTCCCCTAAAAATCTTTGAAATAGGGCCCGTATTTACCCTATAGCACCCCCTCTACC
CCCTCTAGA?????????????????????????????????????????????????????????????????????????
??????????TAAATACTACCGTATGGCCCACCATAATTACCCCCATACTCCTTACACTAT
TCCTCATCACCCAACTAAAAATATTAAACACAAACTACCACCTACCTCCCTCACCAA
AGCCCATAAAAATAAAAAATTATAACAAACCCTGAGAACCAAAATGAACGAAAATC
TGTTCGCTTCATTCATTGCCCCCACAATCCTAGGCCTACCCGCCGCAGTACTGATCAT
TCTATTTCCCCCTCTATTGATCCCCACCTCCAAATATCTCATCAACAACCGACTAATC
ACCACCCAACAATGACTAATCAAACTAACCTCAAAACAAATGATAGCCATACACAA
CACTAAAGGACGAACCTGATCTCTTATACTAGTATCCTTAATCATTTTTATTGCCACA
ACTAACCTCCTCGGACTCCTGCCTCACTCATTTACACCAACCACCCAACTATCTATAA
ACCTAGCCATGGCCATCCCCTTATGAGCGGGCGCAGTGATTATAGGCTTTCGCTCTA
AGATTAAAAATGCCCTAGCCCACTTCTTACCACAAGGCACACCTACACCCCTTATCC
CCATACTAGTTATTATCGAAACCATCAGCCTACTCATTCAACCAATAGCCCTGGCCGT
ACGCCTAACCGCTAACATTACTGCAGGCCACCTACTCATGCACCTAATTGGAAGCGC
CACCCTAGCAATATCAACCATTAACCTTCCCTCTACACTTATCATCTTCACAATTCTA
ATTCTACTGACTATCCTAGAAATCGCTGTCGCCTTAATCCAAGCCTACGTTTTCACAC
TTCTAGTAAGCCTCTACCTGCACGACAACACATAATGACCCACCAATCACATGCCTA
TCATATAGTAAAACCCAGCCCATGGCCCCTAACAGGGGCCCTCTCAGCCCTCCTAAT
GACCTCCGGCCTAGCCATGTGATTTCACTTCCACTCCATAACGCTCCTCATACTAGGC
CTACTAACCAACACACTAACCATATACCAATGATGGCGCGATGTAACACGAGAAAG
CACATACCAAGGCCACCACACACCACCTGTCCAAAAAGGCCTTCGATACGGGATAAT
CCTATTTATTACCTCAGAAGTTTTTTTCTTCGCAGGATTTTTCTGAGCCTTTTACCACT
CCAGCCTAGCCCCTACCCCCCAACTAGGAGGGCACTGGCCCCCAACAGGCATCACCC
CGCTAAATCCCCTAGAAGTCCCACTCCTAAACACATCCGTATTACTCGCATCAGGAG
TATCAATCACCTGAGCTCACCATAGTCTAATAGAAAACAACCGAAACCAAATAATTC

AAGCACTGCTTATTACAATTTTACTGGGTCTCTATTTTACCCTCCTACAAGCCTCAGA
GTACTTCGAGTCTCCCTTCACCATTTCCGACGGCATCTACGGCTCAACATTTTTTGTA
GCCACAGGCTTCCACGGACTCCACGTCATTATTGGCTCAACTTTCCTCACTATCTGCT
TCATCCGCCAACTAATATTTCACTTTACATCCAAACATCACTTTGGCTTCGAAGCCGC
CGCCTGATACTGGCATTTTGTAGATGTGGTTTGACTATTTCTGTATGTCTCCATCTATT
GATGAGGGTCTTA????????????????????????????????????????????????????????????????????
???????????????TTTTAATAATCAACACCCTCCTAGCCTTACTACTAATAATTATTACAT
TTTGACTACCACAACTCAACGGCTACATAGAAAAATCCACCCCTTACGAGTGCGGCT
TCGACCCTATATCCCCCGCCCGCGTCCCTTTCTCCATAAAATTCTTCTTAGTAGCTATT
ACCTTCTTATTATTTGATCTAGAAATTGCCCTCCTTTTACCCCTACCATGAGCCCTAC
AAACAACTAACCTGCCACTAATAGTTATGTCATCCCTCTTATTAATCATCATCCTAGC
CCTAAGTCTGGCCTATGAGTGACTACAAAAAGGATTAGACTGAGCTGAATTGGTATA
TTGTTTAAACAAAACGAATGATTTCGACTCATTAAATTATGATAATCATATTTACCAA
ATGCCCC?????????TAAATATTATACTAGCATTTACCATCTCACTTCTAGGAATACTAG
TATATCGCTCACACCTCATATCCTCCCTACTATGCCTAGAAGGAATAATACTATCGCT
GTTCATTATAGCTACTCTCATAACCCTCAACACCCACTCCCTCTTAGCCAATATTGTG
CCTATTGCCATACTAGTCTTTGCCGCCTGCGAAGCAGCGGTGGGCCTAGCCCTACTA
GTCTCAATCTCCAACACATATGGCCTAGACTACGTACATAACCTAAACCTACTCCAA
TGCTAAAACTAATCGTCCCAACAATTATATTACTACCACTGACATGACTTTCCAAAA
AACACATAATTTGAATCAACACAACCACCCACAGCCTAATTATTAGCATCATCCCCC
TACTATTTTTTAACCAAATCAACAACAACCTATTTAGCTGTTCCCCAACCTTTTCCTC
CGACCCCCTAACAACCCCCCTCCTAATACTAACTACCTGACTCCTACCCCTCACAATC
ATGGCAAGCCAACGCCACTTATCCAGTGAACCACTATCACGAAAAAAACTCTACCTC
TCTATACTAATCTCCCTACAAATCTCCTTAATTATAACATTCACAGCCACAGAACTAA
TCATATTTTATATCTTCTTCGAAACCACACTTATCCCCACCTTGGCTATCATCACCCG
ATGAGGCAACCAGCCAGAACGCCTGAACGCAGGCACATACTTCCTATTCTACACCCT
AGTAGGCTCCCTTCCCCTACTCATCGCACTAATTTACACTCACAACACCCTAGGCTCA
CTAAACATTCTACTACTCACTCTCACTGCCCAAGAACTATCAAACTCCTGAGCCAAC
AACTTAATATGACTAGCTTACACAATAGCTTTTATAGTAAAGATACCTCTTTACGGAC
TCCACTTATGACTCCCTAAAGCCCATGTCGAAGCCCCCATCGCTGGGTCAATAGTAC
TTGCCGCAGTACTCTTAAAACTAGGCGGCTATGGTATAATACGCCTCACACTCATTCT
CAACCCCCTGACAAAACACATAGCCTACCCCTTCCTTGTACTATCCCTATGAGGCAT
AATTATAACAAGCTCCATCTGCCTACGACAAACAGACCTAAAATCGCTCATTGCATA
CTCTTCAATCAGCCACATAGCCCTCGTAGTAACAGCTATTCTCATCCAAACCCCCTGA
AGCTTCACCGGCGCAGTCATTCTCATAATCGCCCACGGACTTACATCCTCATTACTAT
TCTGCCTAGCAAACTCAAACTACGAACGCACTCACAGTCGCATCATAATCCTCTCTC
AAGGACTTCAAACTCTACTCCCACTAATAGCTTTTTGATGACTTCTAGCAAGCCTCGC
TAACCTCGCCTTACCCCCCACTATTAACCTACTGGGAGAACTCTCTGTGCTAGTAACC
ACGTTCTCCTGATCAAATATCACTCTCCTACTTACAGGACTCAACATACTAGTCACAG
CCCTATACTCCCTCTACATATTTACCACAACACAATGGGGCTCACTCACCCACCACAT
TAACAACATAAAACCCTCATTCACACGAGAAAACACCCTCATGTTCATACACCTATC
CCCCATTCTCCTCCTATCCCTCAACCCCGACATCATTACCGG?????????????????????????
?????????????????????????????????????????????????????????????????????????????????????????
?????????????????????????????????????????????????????????????????????????????????????????
????????????????????CACTACTATAACCACCCTAACCCTGACTTCCCTAATTCCCCCCAT
CCTTACCACCCTTGTTAACCCTAACAAAAAAAACTCATACCCCCATTATGTAAAATC
CATTGTCGCATCCACCTTTATTATCAGTCTCTTCCCCACAACAATATTCATGTGCCTA
GACCAAGAAGTTATTATCTCGAACTGACACTGAGCCACAACCCAAACAACCCAGCTC
TCCCTAAGCTTCAAACTAGACTACTTCTCCATAATATTCATCCCTGTAGCATTGTTCG
TTACATGGTCCATCATAGAATTCTCACTGTGATATATAAACTCAGACCCAAACATTA
ATCAGTTCTTCAAATATCTACTCATTTTCCTAATTACCATACTAATCTTAGTTACCGCT
AACAACCTATTCCAACTGTTCATCGGCTGAGAGGGCGTAGGAATTATATCCTTCTTG

CTCATCAGTTGATGACACGCCCGAGCAGATGCCAACACAGCAGCCATTCAAGCAATC
CTATACAACCGTATCGGCGATATCGGTTTCATCCTCGCCTTAGCATGATTTATCCTAC
ACTCCAACTCATGAGACCCACAACAAATAGCCCTTCTAAACGCTAATCCAAGCCTCA
CCCCACTACTAGGCCTCCTCCTAGCAGCAGCAGGCAAATCAGCCCAATTAGGTCTCC
ACCCCTGACTCCCCTCAGCCATAGAAGGCCCCACCCCAGTCTCAGCCCTACTCCACT
CAAGCACTATAGTTGTAGCAGGAATCTTCTTACTCATCCGCTTCCACCCCCTAGCAGA
AAATAGCCCACTAATCCAAACTCTAACACTATGCTTAGGCGCTATCACCACTCTGTTC
GCAGCAGTCTGCGCCCTTACACAAAATGACATCAAAAAAATCGTAGCCTTCTCCACT
TCAAGTCAACTAGGACTCATAATAGTTACAATCGGCATCAACCAACCACACCTAGCA
TTCCTGCACATCTGTACCCACGCCTTCTTCAAAGCCATACTATTTATGTGCTCCGGGT
CCATCATCCACAACCTTAACAATGAACAAGATATTCGAAAAATAGGAGGACTACTCA
AAACCATACCTCTCACTTCAACCTCCCTCACCATTGGCAGCCTAGCATTAGCAGGAA
TACCTTTCCTCACAGGTTTCTACTCCAAAGACCACATCATCGAAACCGCAAACATAT
CATACACAAACGCCTGAGCCCTATCTATTACTCTCATCGCTACCTCCCTGACAAGCGC
CTATAGCACTCGAATAATTCTTCTCACCCTAACAGGTCAACCTCGCTTCCCCACCCTT
ACTAACATTAACGAAAATAACCCCACCCTACTAAACCCCATTAAACGCCTGGCAGCC
GGAAGCCTATTCGCAGGATTTCTCATTACTAACAACATTTCCCCCGCATCCCCCTTCC
AAACAACAATCCCCCTCTACCTAAAACTCACAGCCCTCGCTGTCACTTTCCTAGGACT
CCTAACAGCCCTAGACCTCAACTACCTAACCAACAAACTTAAAATAAAATCCCCACT
ATGCACATTTTATTTCTCCAACATACTCGGATTCTACCCTAGCATCACACACCGCACA
ATCCCCTATCTAGGCCTTCTTACGAGCCAAAACCTGCCCCTACTCCTCCTAGACCTAA
CCTGACTAGAAAAGCTATTACCTAAAACAATTTCACAGCACCAAATCTCCACCTCCA
TCATCACCTCAACCCAAAAAGGCATAATTAAACTTTACTTCCTCTCTTTCTTCTTCCC
ACTCATCCTAACCCTACTCCTAATCACATAACCTATTCCCCCGAGCAATCTCAATTAC
AATATATACACCAACAAACAATGTTCAACCAGTAACTACTACTAATCAACGCCCATA
ATCATACAAAGCCCCCGCACCAATAGGATCCTCCCGAATCAACCCTGACCCCTCTCC
TTCATAAATTATTCAGCTTCCTACACTATTAAAGTTTACCACAACCACCACCCCATCA
TACTCTTTCACCCACAGCACCAATCCTACCTCCATCGCTAACCCCACTAAAACACTCA
CCAAGACCTCAACCCCTGACCCCCATGCCTCAGGATACTCCTCAATAGCCATCGCTG
TAGTATATCCAAAGACAACCATCATTCCCCCTAAATAAATTAAAAAAACTATTAAAC
CCATATAACCTCCCCCAAAATTCAGAATAATAACACACCCGACCACACCGCTAACAA
TCAATACTAAACCCCCATAAATAGGAGAAGGCTTAGAAGAAAACCCCACAAACCCC
ATTACTAAACCCACACTCAACAGAAACAAAGCATACATCATTATTCTCGCACGGACT
ACAACCACGACCAATGATATGAAAAACCATCGTTGTATTTCAACTACAAGAACACC?
???????????CGCAAAATTAACCCCCTAATAAAACTAATTAACCACTCATTCATCGACC
TCCCCACCCCATCCAACATCTCCGCATGATGAAACTTCGGCTCACTCCTTGGCGCCTG
CCTGATCCTCCAAATCACCACAGGACTATTCCTAGCCATGCACTACTCACCAGACGC
CTCAACCGCCTTTTCATCAATCGCCCACATCACTCGAGACGTAAATTATGGCTGAATC
ATCCGCTACCTTCACGCCAATGGCGCCTCAATATTCTTTATCTGCCTCTTCCTACACA
TCGGACGAGGCCTATATTACGGATCATTTCTCTACTCAGAAACCTGAAACATCGGCA
TTATCCTCCTGCTTGCAACTATAGCAACAGCCTTCATAGGCTATGTCCTCCCGTGAGG
CCAAATATCATTCTGAGGGGCCACAGTAATTACAAACTTACTATCCGCCATCCCATA
CATTGGGACAGACCTAGTTCAATGAATCTGAGGAGGCTACTCAGTAGACAGTCCCAC
CCTCACACGATTCTTTACCTTTCACTTCATCTTACCCTTCATTATTGCAGCCCTAGCAG
CACTCCACCTCCTATTCTTGCACGAAACGGGATCAAACAACCCCCTAGGAATCACCT
CCCATTCCGATAAAATCACCTTCCACCCTTACTACACAATCAAAGACGCCCTCGGCTT
ACTTCTCTTCCTTCTCTCCTTAATGACATTAACACTATTCTCACCAGACCTCCTAGGC
GACCCAGACAATTATACCCTAGCCAACCCCTTAAACACCCCTCCCCACATCAAGCCC
GAATGATATTTCCTATTCGCCTACACAATTCTCCGATCCGTCCCTAACAAACTAGGAG
GCGTCCTTGCCCTATTACTATCCATCCTCATCCTAGCAATAATCCCCATCCTCCATAT
ATCCAAACAACAAAGCATAATATTTCGCCCACTAAGCCAATCACTTTATTGACTCCT
AGCCGCAGACCTCCTCATTCTAACCTGAATCGGAGGACAACCAGTAAGCTACCCTTT

TACCATCATTGGACAAGTAGCATCCGTACTATACTTCACAACAATCCTAATCCTAAT
ACCAACTATC????????????????????????????????????????????????????????????????????????????????
????????????????????????????????????????????????????????????????????????????????????????????????
????????????????????????????????????????????????????????????????????????????????????????????????
????????????????????????????????????????????????????????????????????????????????????????????????
????????????????????????????????????????????????????????????????????????????????????????????????
????????????????????????????????????????????????????????????????????????????????????????????????
?????????????????ATAGGGGTCCCTTGACCACCATCCTCCGTGAAATCAATATCCCGC
ACAAGAGTGCTACTCTCCTCGCTCCGGGCCCATAACACTTGGGGGTAGCTAAAGTGA
ACTGTATCCGACATCTGGTTCCTACTTCAGGGCCATAAAGCCTAAATAGCCCACACG
TTCCCCTTAAATAAGACATCACGATG

**GD/P9L: *Manis javanica* + *Panthera tigris***

>Manis_javanica_GDP9L_mtDNA [consensus reconstructed from 13,770 reads]
GTTAATGTAGCTTAAAACCAAAGCAAAGCATTGAAAATGCTTAGATGAGTCCACAA
ACTCCATAAACACACAGGTTTGGTCCCAGCCTTTTTATTAATTTTCAATAGGATTACA
CATGCAAGTATCCGCCCCCCAGTGAAAATGCCCTTCAAGTTATCACAGTACCTGAAG
GAGCTGGCATCAAGCACGCCAATAAACGCAGCTAAMGACGCCTTGCAGAGCCACAC
CCCCACGGGAAACAGCAGTGATAAAAATTAGGCTATAAACGAAAGTTCGACCTAGC
CATATTGCATTGGGTTGGTAAATCTCGTGCCAGCCACCGCGGTCATACGATTAACCC
TAGCTAATAAAAAACCGGCGTAAAGCGTGCTAAGATRAATCCAATCCAAATAAAGT
TAAGCCCTGACCAGGCCGTAAAAAGCCGTGGTTRCCGTAAAAATAAACTACGAAAG
TAACTTTAATTCACATCGACACACGATAGCTAAGACCCAAACTGGGATTAGATACCC
CACTATGCTTAGCCCTAAACCTAAATAATTCACCAGACAAAATTATTCGCCAGAGAA
CTACTGGCAACAGCCTAAAACTCAAAGGACTTGGCGGTGCTTTACATCCCCCTAGAG
GAGCCTGTTCTGTAATCGATAAACCCCGATAAACCCTACCAATTCTAGCTAATACAG
CCTATATACCGCCATCTCCAGCAAACCCTAAAAAGGAAGCACAGTAAGCAAGACTA
TGAAAACATAAATACGTTAGGTCAAGGTGTAGCCCATGAATTGGGAAGAAATGGGC
TACATTTTCTAAAATAGAACACAAACGAACGCCCTAATGAAAACGAGGGCCAAAGG
AGGATTTAGCAGTAAGCTGAGAATAGAAAGCTCAACTGAACCCGGCCCTAAAGCAC
GCACACACCGCCCGTCACCYTCTTCAAATCCCCAAGAATACCTAAACATATTAAACA
AACACCAAGGCATGAGAAGAGACAAGTCGTAACAAGGTAAGCATACTGGAAGGTGT
GCTTGGATCACCAAAGTGTAGCTTAAACAAAGCATCTGGCCTACACCCAGAAGATCT
CAATAACATGACCACTTTGAACAAATCCTAGCCCAACCAACACCCAACATACAACCA
AACAGAGCACATAAACCAAAGCATTTACTAGACTAAAGTATAGGCGATAGAAATTA
CACAACGGCGCTATAGAAAAAGTACCGCAAGGGAAAGATGAAAGATGCATTCAAAG
TACAAAAAAGCAAAGATTACCCCTTGTACCTTTTGCATAATGGACTAACTAGAAACA
CCCTAGCAAAGAGAACTTAAGCTAGAAACCCCGAAACCAGACGAGCTACCTACGAG
CAGTTTAAAGAACCCACTCATCTATGTGGCAAAATAGTGAGAAGACTTGTAGGTAGA
GGTGAAAAGCCTAACGAGCCTGGTGATAGCTGGTTGTCCAAGAAATGAATCTAAGTT
CAACTTGAAGCATACCCAAAAGCCCAAAAACTATAATGTAGGCTTCAAGTATAGTCT
AAAAAGGTACAGCTTTTTAGAAACAGAATTAAATCTTAATTAGTGAGTAAACAATAC
AACAACCATAGTTGGCCTAAAAGCAGCCATCAATTAAGAAAGCGTCAAAGCTCAAC
AATAAGACATAAATAATACCACAAATAAAAATCAACTCCTAAACCAATATTGGACC
AATCTATCAATAAATAGAAGAAATACTGTTAGTATGAGTAACAAGAAATAGATCTCC
TTGCACAAGCTTATATCAGAACGGATGACCCACTGATAATTAACAACGAAACAAATC
AAACCCAAAAATAGAACTTTGTTAAATAGATTGTTAACCCAACACAGGCGTGCATAT
TCTCAAAGGAAAGGTTAAAACAAATGAAAGGAACTCGGCAAACACAAGCCCCGCCT
GTTTACCAAAAACATCACCTCTAGCATAACCAGTATTAGAGGCACTGCCTGCCCAGT
GACTCGCGTTAAACGGCCGCGGTATTCTGACCGTGCAAAGGTAGCATAATCACTTGT
TCTCTAAATAAGGACTAGTATGAACGGCTAGACGAGGGTTTTACTGTCTCTCATCTGT
AACCAGTGAAATTGACCTCCCCGTGAAGAGGCGGGAATAATATAATAAGACGAGAA

GACCCTATGGAGCTTTAATTAAACAGCTTAAAATTAACCCAACACCACTCTAACAGA
GATATAACAAACCAATTAACCTAAGCTGTCAATTTTGGTTGGGGTGACCTCGGAGCA
AAAAACAACCTACGAGCGGTCAAATCCAGACTAACAAGTCCAGATAATCCGTTAATT
GATCCAACAACTTGATCAACGGAACAAGTTACCCTAGGGATAACAGCGCAATCCTAT
TCGAGAGTCCATATCGACAATAGGGTTTACGACCTCGATGTTGGATCAGGACATCCC
AATGGTGCAGAAGCTATTAATGGTTCGTTTGTTCAACGATTAAAGTCCTACGTGATCT
GAGTTCAGACCGGAGTAATCCAGGTCGGTTTCTATCTATTAATACACTTCTCCCAGTA
CGAAAGGACAAGAGAAGTAGGGCCTACCTCACACAGGCGCCCTCGAATCAATAAAT
GACTACCCTCTTAATTTAGCCAATTCACAACAACCCCCAGTCCTAGAACCAGGGCCD
GSTRGGGTAGCAAAGCACGGCAACTGCACAAGACTTAAGCTCTTGTAACAGAGGTTC
AACTCCCCTCCCTAGCAGCAATGTACACAATCAACATAATAATAATGATCATCCCCA
TCCTACTAGCCGTG???????????????????????????????????????????????????????????????????
????????????????????????????????????????????????????????????????????????????????????????
????????????????????????????????????????????????????????????????????????????????????????
????????????????????????????????????????????????????????????????????????????????????????
????????????????????????????????????????????????????????????????????????????????????????
????????????????????????????????????????????????????????????????????????????????????????
????????????????????????????????????????????????????????????????????????????????????????
????????????????????????????????????????????????????????????????????????????????????????
????????????????????????????????????????????????????????????????????????????????????????
????????????????????????????????????????????????????????????????????????????????????????
????????????????????????????????????????????????????????????????????????????????????????
????????????????????????????????????????????????????????????????????????????????????????
????????????????????????????????????????????????????????????????????????????????????????
????????????????????????????????????????????????????????????????????????????????????????
??????????????????TAGTCCCTATCCTAATGAAAAACTTTAGCCCACGRGCCATAGAAG
CAGCCACAAAATACTTCCTAATCCAAGCCACCGCGTCCATACTACTAATGCTAGCCA
TTACCCTAGACCTGATATCTTCTGGACAATGAACCATCACAAAAACAC??????????????
????????????????????????????????????????????????????????????????????????????????????????
?????????????????????????????????????????????????????CTATACCAAATAATACCCACAAT
CAACACAAATATACTGACAACCATAGCCGCACTCTCAATCCTCATCGGCGGTTGAGG
AGGGCTAAACCAGACCCAACTACGAAAAATCATAGCATACTCCTCAATCGCCC??????
????????????????????????????????????????????????????????????????????????????????????????
????????????????????????????????????????????????????????????????????????????????????????
????????????????????????????????????????????????????????????????????????????????????????
????????????????????????????????????????????????????????????????????????????????????????
????????????????????????????????????????????????????????????????????????????????????????
????????????????????????????????????????????????????????????????????????????????????????
????????????????????????????????????????????????????????????????????????????????????????
????????????????????????????????????????????????????????????????????????????????????????
????????????????????????????????????????????????????????????????????????????????????????
????????????????????????????????????????????????????????????????????????????????????????
????????????????????????????????????????????????????????????????????????????????????????
???????????????????????????????????????????????????????????????CGTTACCGCACATGCA
TTTGTAATAATCTTCTTCATAGTCATGCCAATCATAATCGGGGGCTTTGGAAACTGAC
TAGTGCCCCTGATAATTGGAGCCCCTGACATGGCATTCCCCCGTATAAACAACATAA
GTTTCTGACTRCTCCCCCCCTC??TCCTACTTCTTCTGGCCTCTTCCATAGTCGAAGCA
GGGGCTGGAACCGGCTGAACTGTATACCCCCCTTTAGCAGGAAATTTAGCACACGCA
GGAGCATCTGTAGACCTAACCATCTTCTCTCTTCATCTAGCGGGTGTC???????????????
????????????????????????????????????????????????????????????????????????????????????????
????????????????????????????????????????????????????????????????????????????????????????
????????????????????????????????????????????????????????????????????????????????????????

?????????????????????????????????TATTACTCCGGGAAAAAGAGCCCTTTGGATACAT
GGGCATAGTCTGAGCAATAATATCCATTGGCTTCCTGGGCTTCATTGTATGGGCACA
CCACATGTTCACAGTGGGAATAGACGTTGATACACGGGCCTACTTCACATCAGCCAC
C???????????????????????????????????????????TAGCAACCCTGCACGGAGGAAACGTAA
AATGGGCCCCAGCTATACTMTGGGCCCTGGGCTTTATYTTCCTGTTTACAGTCGGGG
GTCTAACTGGCATCGTGCTGGCCAACTCGTCCCTAGATATCGTCCTCCATGATACCTA
CTACGTAGTAGCCCACTTCCACTACGTTCTCTCCATGGGAGCAGTTTTCGCCATCATA
GGAGGCTTCGTCCATTGATTCCCCCTGTTCTCAGGMTACACGCTCAACAACACATGG
GCAAAAGTTCACTTTACAATCATATTCGTGGGCGTAAACATGACCTTTTTCCCCCAAC
ACTTCCTTGGRCTGTCTGGAATACCTCGACGATACTCGGACTACCCAGACGCTTACA
CAATGTGAAACACTGTGTCCTCCATAGGATCCTTCATCTCATTAACCGCCGTAATACT
CATGGCCTTCATAATCTGAGAGGCATTCGCCTCTAAACGGGAAGTCCTAATAGTGGA
GTCCACAAATACCAACCTCGAATGACTACACGGCTGCCCACCACCCTACCACACATT
CGAAGAACCTGCTTTCGTAAATCTGGTCAAAACAAGAGAGGAAGGAATCGAACCCT
CAAACAATGGTTTCAAGCCAATATCATARCCCCTA?????????????????????????????????
????????????????????????????????????????????????????????????????????????????????????????
????????????????????????????????????????????????????????????????????????????????????????
????????????????????????????????????????????????????????????????????????????????????????
????????????????????????????????????????????????????????????????CGCCCTGACAGTAAA
AACAATGGGCCATCAATGATACTGAAGCTACGAATACACAGACTACGAAGACCTAA
GCTTCGACTCATACATAATCCCAACTCAAGACCTAAAACCCGGCGAACTCCGACTTC
TAGAAGTGGACAACCGACTTGTACTACCCATAGACACAACCATCCGCATGCTCATCT
CATCTGAAGACGTCCTGCACTCTTGAGCCRTCCCATCCCTGGGCCTAAAA????????????
?????????????ACCAAACAACCCTGATATCAACCCGGCCCGGCCTATTCTACGGACAGT
GCTCAGAAATCTGTGGCTCAAATCACAGCTTCATGCCAATTGTCCTCGAACTAGTAC
CCCTAAAAACATTTGAAAACTGAACTACRTCCCTACTGTAATTCA???????????????????
????????????????????????????????????????????????????????????????????????????????????????
????????????????????????????????????????????????????????????????????????????????????????
????????????????????????????????????????????????????????????????????????????????????????
????????????????????????????????????????????????????????????????????????????????????????
????????????????????????????????????????????????????????????????????????????????????????
????????????????????????????????????????????????????????????????????????????????????????
????????????????????????????????????????????????????????????????????????????????????????
????????????????????????????????????????????????????????????????????????????????????????
????????????????????????????????????????????????????????????????????????????????????????
?????????????????????????????TCGCACTCATTCAGGCCTACGTCTTCACYCTTCTAATCA
GCCTCTACTTACATGATAACACATAATGACCCACCAAACCCACTCATACCACATAGT
AAACCCAAGCCCTTGACCCCTAACCGGGGCCCTATCTGCCCTCCTAATAACATC?????
????????????????????????????????????????????????????????????????????????????????????????
????????????????????????????????????????????????????????????????????????????????????????
????????????????????CTTCTTCTGAGCATTCTACCACTCAAGCCTAGCACCTACCCCCGA
ACTAGGAGGATGCTGACCCCCCACAGGCATCAACCCGCTAAACCCRCTGGAAGTACC
TCTACTCAACACATCCGTTCTTTTAGCCTCAGGAGTATCAATTACATG???????????????
????????????????????????????????????????????????????????????????????????????????????????
????????????????????????????????????????????????????????????????????????????????????????
????????????????????????????????????????????????????????????????????????????????????????
????????????????????????????????????????????????????????????????????????????????????????
????????????????????????????????????????????????????????????????????????????????????????
????????????????????????????????????????????????????????????????????????????????????????
????????????????????????????????????????????????????????????????????????????????????????
????????????????????????????????????????????????????????????????????????????????????????
????????????????????????????????????????????????????????????????????????????????????????

???????????????????????????????????????????????????????????????????????????????ATTAAT
GTACCGATCACACCTAATGTCCTCACTACTATGTCTAGAGGGCATAATACTCTCCCTA
TTCGTAATAATCACCATTACAATCCTAACTAACCACTTCACACTAGCCAACATAGCA
CCCATCATCCTCCTCGTACTAGCAGCATG????????????????????????????????????????????
??????????????????????????????????????????????????????????????????????????????????ACAT
GACTATCGAACAAAAACATAGTGTGAATCAACGTAACATCACACAGCATGCTAATC
AGCCTAACATCTCTAGGCATCTTGGGCCAGCATGACCACAACAACATAAGCCTCTCA
GCCATCTTCTTCTCTGACCCCCTATCAGCACCC??????????????????????????????????????
????????????????????????????????????????????????????????????????????????????????????????
????????????????????????????????????????????????????????????????????????????????????????
????????????????????????????????????????????????????????????????????????????????????????
????????????????????????????????????????????????????????????????????????????????????????
???????????????????TAGCATTCATAGTAAAAATACCCCTATACGGACTACACCTTTGAC
TGCCAAAAGCCCACGTAGAGGCCCCCATTGCAGGCTCCATAGTACTTGCAGCCGTAC
TACTAAAACTAGGCGGATACGGCATAATACGCATCACTATCATGCTAGAACCAATAA
CAACATTCATAGCATACCCCTTCCTAATACTATCCCTGTGAGGAATAATCATAACAA
GCTCCATCTGCCTTCGCCAAACCGACCTAAA????????????????????????????????????????
????????????????????????????????????????????????????????????????????????????????????????
????????????????????????????????????????????????????????????????????????????????????????
???????????GACTCCTAGCAAGCCTAACCAACCTGGCCCTCCCCCCCAGCAWCAACCT
AATCGGAGAACTATTCGTAGTAGTATCAACATTCTCGTGGTCYAACACCACCATCAT
CCTTACTGGAACAAACATTATCATCACAGCCACCTACTCCCTAT?????????????????????
????????????????????????????????????????????????????????????????????????????????????????
????????????????????????????????????????????????????????????????????????????????????????
????????????????????????????????????????????????????????????????????????????????????????
????????????????????????????????????????????????????????????????????????????????????????
????????????????????????????????????????????????????????????????????????????????????????
????????????????????????????????????????????????????????????????????????????????????????
????????????????????????????????????????????????????????????????????????????????????????
????????????????????????????????????????????????????????????????????????????????????????
????????????????????????????????????????????????????????????????????????????????????????
??????????TGGCACGGACGAACAGATGCAAACACAGCAGCCATACAGGCCATCCTATA
CAACCGCATCGGAGACATTGGACTAATCCTGTCAATGGCATGATTCCTCACAAACCT
AAACTCATGGGACCTCCAACAAATCCTCATGCTAAACCCCGAAAACACAAACATCCC
CCTAGCGGGCCTACTACTAGCAGCAACTGGAAAGTCCGCACAATTTGGGCTACACCC
ATGACTCCCCTCAGCCATAGAAGGCCCAACCCCAGTATCAGCCCTACTCCA??????????
????????????????????????????????????????????????????????????????????????????????????????
?????????????????????????????????????????????????????????????????????????????????GCCA
ACTAGGTCTAATAATAGTCACCATCGGAATTAACCAACCACACCTAGCCTTCATACA
CATCTGCACCCACGCCTTTTCAAAGCCATACTATTCATGTGCTCCGGATCGATCATC
CACAGCCTAAC??????????????????????????????????????????????????????????????????TCA
TTATTGGCAGCCTAGCCCTAACAGGAACCCCATTCCTAACAGGCTTCTACTCTAAAG
ACCTAATCATCGAAACCGCTAACATATCCCACACAAACGCCTGAGCCCTGCTCATCA
CACTAGTAGCCACCTCCCT???????????????????????????????????????????????????????????
???????????????????????????????CTCACTAACAAACCCAATCAAACGCCTAATGCTAGGG
AGCATCTTCGCCGGATTCCTACTATACAACAACATCCCCCCAACAACCATGCCCCAA
ACAACTATACCACAATACCTAAAATACACCGCCCTCGCCACTACAATCCTAGG??????
????????????????????????????????????????????????????????????????????????????????????????
???????????????????????????????????????????????????????????AGCATCCCTTATTCTAGAC
CTAATCTGACTAGAAAATATYATACCAAAGTCAATCTCACACTTCCACATAAAAGCC
TCCCTAATAACATCCAACCAAAAAGGCTCAATCAAACTATACTTCCTGTCCTTCGCA
GTAACAATAATCCTCGCTCTCCTCATATTCTATTCCCCCGAGTAACCTCCATAACAAC

AACAACACAAATAAACAAAGACCACCCGGTCATAATCACAAACCAAGCACCATGGC
TATAAAGAGCAGCAACACCCGCAGCTTCCTCA?????????????????????????????????????????
??????????????????????????????????????????????????????????????????????????????????????????????
??????????????????????????????????????????????????????????????????????????????????????????????
??????????????????????????????????????????????????????????????????????????????????????????????
??????????????????????????????????????????????????????????????????????????????????????????????
?????????????????????????????????????CCATGACCTATGACACGAAAAATCATCGTTGTA
ATTAAACTACGGAAACCTAATGACAAACATCCGAAAAACACACCCCCTACTCAAAA
TCATCAACAACTCCTTTATTGATCTCCCCACCCCCTCTAACATCTCAGCATGATGAAA
CTTCGGATCACTACTAGGAATCTGCTTAGTCCTACAAATTCTCACAGGACTATTTCTA
G?CATACACTACACAGCAGACACAGCAACCGCATTCTCATCAGTAACTCACATCTGC
CGAGATGTAAACTACGGATGAATTATCCGATACAYACACGCTAACGGAGCTTCCCTG
TTCTTCATCTGCCTATTCGCA????????????????????????????????????????????????????????????
??????????????????????????????????????????????????????????????????????????????????????????????
??????????????????????????????????????????????????????????????????????????????????????????????
??????????????????????????????????????????????????????????????????????????????????????????????
??????????????????????????????????????????????????????????????????????????????????????????????
??????????????????????????????????????????????????????????????????????????????????????????????
??????????????????????????????????????????????????????????????????????????????????????????????
??????????????????????????????????????????????????????????????????????????????????????????????
??????????????????????????????????????????????????????????????????????????????????????????????
??????????????????????????????????????????????????????????????????????????????????????????????
?GAGTCTTTGTAGTATATGTCAATACACTGGTCTTGTAAACCAGCAAAGGAATTAACC
CTCCCCAAGACTCAGGAAGAAGACAAAAGCCCTACCATCAGCACCCAAAGCTGAAA
TTCTCAATAAACTACTTCCTG??????????????????????????????????????????????????????????
??????????????????????????????????????????????????????????????????????????????????????????????
??????????????????????????????????????????????????????????????????????????????????????????????
??????????????????????????????????????????????????????????????????????????????????????????????
??????????????????????????????????????????????????????????????????????????????????????????????
???GCCCATACGGACTGTGKGGTAGTTAAAGTGWAACTATASCTGKCATCTGRTTCTT
ACTTCATGTTCATCTTATATCTAGCCGCTCACTCGTTCCTCTTAAATAAGACATCTCG
ATGGATTAATTACTAATCAGCCCATGCCTAACATAAC???????????????????????????????
??????????????????????????????????????????????????????????????????????????????????????????????
??????????????????????????????????????????????????????????????????????????????????????????????
??????????????????????????????????????????????????????????????????????????????????????????????
??????????????????????????????????????????????????????????????????????????????????????????????
??????????????????????????????????????????????????????????????????????????????????????????????
?????????????????????????????????????AATAAAAASAATATACAATAAATACAAATATA
GATCAATTCTTATAGGCGCCGATAGCATAAACTATCTGCCCCCTATGTACAATTGAA
TTAAAGTTAATGTAGCTTAAAACCAAAGCAAAGCATTGAAAATGCTTAGATGAGTCC
ACAAACTCCATAAACACACAGGTTTGGTCCCAGCCTTTTTATTAATTTTCAATAGGAT
TACACATGCAAGTATCCGCCCCCAGTGAAAATGC

>Panthera_tigrus_GDP9L_mtDNA [consensus reconstructed from 3,447 reads]
??????????????????????????????????????????????????????????????????????????????????????????????
??????????????????????????????????????????????????????????????????????????????????????????????
??????????????????????????????????????????????????????????????????????????????????????????????
??????????????????????????????????????????????????????????????????????????????????????????????
??????????????????????????????????????????????????????????????????????????????????????????????
??????????????????????????????????????????????????????????????????????????????????????????????
??????????????????????????????????????????????????????????????????????????????????????????????
??????????????????????????????????????????????????????????????????????????????????????????????

??????????????????????????????????????????????????????TATCTAGGCACAAGGCCTAAGA
TTAACGTTTACAAACTCTACCAACCCCATCATTACCAATTATTAATACTAAATCATAA
CTTCGTTCGCAGTTATCTATAGATACGACAACCCGATCTCTAATTCGTCCCTATCGAA
CAATATTTACATACCCAACAACCCTATGTCTTGGTTAATGTAGCTTAAATATATTAAA
GCAAGGCACTGAAAATGCCTAGATGGGCCGCCAGGCTCCATAAACATAAAGGTTTG
GTCCTAGCCTTTCCATTAGTTGTTAATAAAATTACACATGCAAGCCTCCGCATCCCGG
TGAAAATGCCCTCTAAATCACCCAGTGATCCAAAGGAGCCGGTATCAAGTACACAAC
CATTGTAGCTCATGACACCTTGCTCAGCCACACCCCCACGGGACACAGCAGTGATAA
AAATTAAGCCATGAATGAAAGTTCGACTAAGCTATATTAAATTAGGGTTGGTAAATT
TCGTGCCAGCCACCGCGGTCATACGATTAACCCAAACTAATAGACCCACGGCGTAAA
GCGTGTTACAGAAAAAAGTATACTAAAGTTAAGCCTTAACTAGGCTGTAAAAAGCC
ACAGTTAACGTAAAAATACAGCACGAAAGTAACTTTAATATTTCTGACCACACGATA
GCTAAGACCCAAACTGGGATTAGATACCCCACTATGCTTAGCCCTAAACCTAGATAG
TTAACCCAAACAAAACTATCCGCCAGAGAACTACTAGCAACAGCTTAAAACTCAAA
GGACTTGGCGGTGCTTTATATCCCTCTAGAGGAGCCTGTTCCATAATCGATAAACCC
CGATAAACCTCACCATCTCTTGCTAATTCAGCCTATATACCGCCATCTTCAGCAAACC
CTAAAAAGGAAGAAAAGTAAGCACAAGTATCTTAACATAAAAAAGTTAGGTCAAGG
TGTAGCCCATGAGATGGGGAAGTAATGGGCTACATTTTCTATAACTAGAACATCCAC
GAAAATCCTTATGAAATTAAGTATTAAAGGAGGATTTAGTAGTAAATTCGAGAATAG
AGAGCTCGATTGAATCGGGCCATGAAGCACGCACACACCGCCCGTCACCCTCCTCAA
GTGATTAGACCCCAAAGAAACCTATTCAAACCACTACACCCACAAGAGGAGACAAG
TCGTAACAAGGTAAGCATACTGGAAAGTGTGCTTGGATAACAAGATGTAGCTTAAAC
AAAGCATCTGGCCTACACCCAGAAGATTTCATATTAAACTGACCATCTTGAGCTAGA
GCTAGCCCAACTACCCATAAACACAACTAACATTAGAAAGTAAAACAAAACATTTA
GTTACTCTAAAAAGTATAGGAGATAGAAATTTAACTCGGCGCTATAGAAAAAGTACC
GCAAGGGAATGATGAAAGAAAAAACTAAAAGCACTATACAGCAAAGATTACCCCTT
GTACCTTTTGCATAATGAATTAGCTAGAATAACCTAACAAAGAGAACTTCAGCTAGG
CCCCCCGAAACCAGACGAGCTACCCATAAACAATCTATTACAGGATGAACTCGTCTA
TGTTGCAAAATAGTGAGAAGATTTATGGGTAGAGGTGAAAAGCCTAACGAGCCTGG
TGATAGCTGGTTGCCCAGAACAGAATCTTAGTTCAGCTTTAAACTTACCTCAAAAAC
CCTAAAATTCCAATGTAAGTTTAAAATATAGTCTAAAAAGGTACAGCTTTTTAGAAT
CTAGGATACAGCCTTAATTAGAGAGTAAGCATATAACACAAACCATAGTTGGCCTAA
AAGCAGCCACCAATTAAGAAAGCGTTCAAGCTCAACAATCAAAACATCTCAATGTC
AAAAAACGCAACCAACTCCTAATCTAAAACTGGGCTAATCTATTTAACAATAGAAGC
AATAATGCTAATATGAGTAACAAGAAATACTTCTCCCGCGCATAAGCTTATATCAGA
ACGGATAACCACTGATAGTTAACAACAAGATATATATAACCTAACTACAAGCAAAA
TATCAGACTAATTGTTAACCCAACACAGGCATGCAATTTAGGGAAAGATTAAAAGA
AGTAAAAGGAACTCGGCAAACACAAGCCCCGCCTGTTTACCAAAAACATCACCTCTA
GCATTTCCAGTATTAGAGGCACTGCCTGCCCAGTGACATTAGTTAAACGGCCGCGGT
ATCCTGACCGTGCAAAGGTAGCATAATCATTTGTTCCTTAAATAGGGACTTGTATGA
ATGGCCACACGAGGGCTTTACTGTCTCTTACTTCCAATCCGTGAAATTGACCTTCCCG
TGAAGAGGCGGGAATACGACAATAAGACGAGAAGACCCTGTGGAGCTTTAATTAAT
CGACCCAAAGAGACCTTAATAACCAACCGACAGGAACAACAGACCTCTGCCATGGG
CCGACAATTTAGGTTGGGGTGACCTCGGAGAATAAAACAACCTCCGAGTGATTTAAA
TCTAGACTGACCAGTCGAAAGTATTACATCACTTATTGATCCAAAGCTTGATCAACG
GAACAAGTTACCCCAGGGATAACAGCGCAATCCTATTTCAGAGTCCATATCGACAAT
AGGGTTTACGACCTCGATGTTGGATCAGGACATCCCGATGGTGCAGCAGCTATCAAA
GGTTCGTTTGTTCAACGATTAAAGTCCTACGTGATCTGAGTTCAGACCGGAGTAATC
CAGGTCGGTTTCTATCTATTAAATAATTTCTCCCAGTACGAAAGGACAAGAGAAATA
AGGCCCACTTTACCAAAGCGCCTTTAACCAAATAGATGATATAATCTCAATCTAAAC
AGTTTATCTAAACATATCACCCGTAGAGCTCGGGTTTGTTAGGGTGGCAGAGCCCGG
CAATTGCATAAAACTTAAGCTTTTACTATCAGAGGTTCAACTCCTCTCCCTAACAGCA

TGTTTATAATCAATATTCTCTCATTAATTATCCCTATTCTTCTCGCCGTAGCCTTCCTA
ACCCTAGTTGAACGTAAAGTACTGGGCTACATACAACTCCGTAAAGGACCAAACGTC
GTAGGACCATACGGCCTACTTCAGCCTATTGCAGACGCCATGAAACTCTTCACTAAA
GAACCCCTCCGACCCCTCACATCCTCCACATTCATATTCATCACAGCACCCA????????
????????????????????????????????????????????????????????????????????????????????????????????????
????????????????????????????????????????????????????????????????????????????????????????????????
???????????????????????????????????????????????????ACTACTAATAAATGGATCCTTCACATT
AGCTGCACTAATTACCACCCAAGAATACATCTGGCTCATCATCCCTGCATGACCCCT
AGCCATAATATGATTCATCTCCACACTAGCAGAAACCAACCGAGCTCCATTTGATCT
AACAGAAGG???????????????????????????????????????????????????????????????????????????????
????????????????????????????????????????????????????????????????????????????????????????????????
????????????????????????????CTAACAACCACCTTCCTATGGATCCGAGCATCTTATCCAC
GATTCCGATATGACCAATTAATGCACCTCCTATGAAAAAACTTCCTACCCCTTACTCT
AGCCCTATGCATATGACACGTCTCCCTACCCATCATTACAGCAAGCATTCCACCCCA
AACATAAGAAATATGTCTGACAAAAGAATTACTTTGATAGAGTAAAACATAGAGGT
TTAAGCCCTCTTATTTCTAGAATTATAGGAATCGAACCTAATCCTAAGAATCCAAAA
ATCTTCGTGCTACCAATATTACACCACATTCTAAGTAAGGTCAGCTAAATAAGCTAT
CGGGCCCA??????????????????????????????????????????????????????????????????????????????????
????????????????????????????????????????????????????????????????????????????????????????????????
????????????????????????????????????????????????????????????????????????????????????????????????
????????????????????????????????????????????????????????????????????????????????????????????????
????????????????????????????????????????????????????????????????????????????????????????????????
??????????????????????????????????????????????????????????????????????????????CTAGCAATAG
CCATTATATCAGTTATAATCGGAGGCTGAGGGGGACTTAACCAAACCCAGCTACGAA
AAATCATAGCATACTCCTCAATCGCCCATATAGGTTGAATAACAGCCATCATAATAT
ATAGCCCCACAATAATAATTTTAAAC????????????????????????????????????????????????????
????????????????????????????????????????????????????????????????????????????????????????????????
???????????????????????????????????????????????????????????????????????????????????????????????
CCACACTACTAGCCATAACAGCACTACTTAACCTRTACTTCTACATACGACTAACAT
ATACCACTGCACTAACTATATTCYCCTCAAACAACTGTATAAAAATAAAATGACGGT
TCAAATGCACAAAAAAAATAATCTTTTTACCCCCC?????????????????????????????????????
????????????????????????????????????????????????????????????????????????????????????????????????
????????????????????????????????????????????????????????????????????????????????????????????????
????????????????????????????????????????????????????????????????????????????????????????????????
????????????????????????????????????????????????????????????????????????????????????????????????
????????????????????????????????????????????????????????????????????????????????????????????????
??????????????????????????????????????????????????????????????????????????????????TCAGTCTC
CTAATTCGAGCCGAACTGGGTCAACCTGGCACACTACTAGGAGATGACCAAATTTAT
AATGTAGTAGTTACTGCCCATGCCTTTGTGATAATCTTTTTTATAGTAATGCCTATTA
TAATTGGAGGATTCGGAAACTGGCTAGTTCCGTTAATAATCGGAGCCCCCGATATGG
CATTCCC?????????????????????????????????????????????????????????????????????????????????????
????????????????????????????????????????????????????????????????????????????????AACTATTTT
TCACTACACCTAGCAGGCGTCTCCTCAATCTTAGGTGCTATTAATTTTATTACTACTA
TTATTAATATAAAACCGCCCGCTATGTCCCAATACCAAACACCCCTGTTTGTTTGATC
GGTTCTAATTACTGCTGTGTTGCT??????????????????????????????????????????????????????AG
ATCGAAATCTAAATACCACATTTTTTGATCCTGCCGGGGGAGGAGACCCCATCTTAT
ATCAACACCTATTCTGATTCTTCGGTCACCCAGAAGTCTATATCTTAATCCTGCCCGG
GTTTGGAATAATTTCACATATTGTCACCTACTA??????????????????????TGGCTACATGG
GGATAGTCTGAGCCATAATGTCAATTGGCTTTCTGGGCTTTATCGTATGGGCCCATCA
CATGTTTACTGTAGGGATAGATGTGGATACACGAGCATACTTTACGTCAGCTACTAT
AATTATCGCTATTCCTACTGGGGTAAAAGTATTTAGCTGATTGGCCACTCTTCACGGG
GGTAATATTAAATGGTCTCCCGCTATACTATGGGCTTTGGGATTCATTTTCCTATTCA

CCGT???????????????????????????????????????????????????????????????????????????????????????????
?????????????????????????????????????????????????????????????????????????????????????????????????
?????????????????????????????????????????????????????????????????????????????????????????????????
??CTCGACGTTATTCTGACTATCCAGACGCGTATACAACTTGAAACACAATCTCCTCA
ATAGGCTCTTTTATTTCACTAACAGCAGTAATATTAATAGTCTTTATAATGTGAGAAG
CTTTCGCATCAAAGCGAGAAGTAGCCACAGTGGAACTAACCACAACTAATCTCGAAT
GACTTCACGGATGTCCTCCTCCGTATCACACATTTGAAGAGCCAGCC????????????????
???????????????????????TCCTTAGACTGGTTTCAAGCCAATACCATAACCACTATGTCTTT
CTCAATTAAGAAGTATTAGTAAAATAATTACATAACTTTGTCAAAGTTAAATTATAG
GTTTAAGCCCTATGTGCTTCCATGGCATACCCCTTCCAACTAGGTTTTC??????????????C
CCCATTATAGAAGAGCTTTTACACTTCCATGATCATACATTAATAATTGTATTCCTAA
TTAGCTCCCTAGTCCTCTACATTATCTCATTAATACTGACAACTAAACTTACGCATAC
AAGCACAATAGATGCCCAAGAAGTAGAAACTA?????????????????????????????????????????
?????????????????????????????????????????????????????????????????????????????????????????????????
??????????????????????????????????TAAGCTTCGACTCCTACATAATCCCCACTCAAGAGTT
AAAGCCCGGAGAGCTCCGACTACTAGAAGTTGATAACCGAGTAGTGTTGCCAATAG
AAGTGACTATTCGCATGTTAGTCTCATCAGAGGACGTACTGCACTCGTGAGCCATCC
CATCCCTGGGCCTAAAAACTGACGCTATCCCAGGCCGACTAAACCAAACAACCCTAA
TAGGCACACGGCCTGGGCTATATTATGGTCAGTGCTCAGAAATCTGCGGCTCAAATC
ACAGTTTTATGCCCATTGTCCTTGAACTAGTCCCGCTGTCATATTTCGAAAAATGA???
??????ATGCTGTAATTTCACTAAGAAGCTAAATTAGCGTTAACCTTTTAAGTTAAAAA
CTGGGAGTTCAAACCTCCCCTTAGTGACATGCCACAGTTAGACACATCAACCTGGTT
TATTACTATTATTTCAATAAT???????????????????????????????????????????????????????????????
?????????????????????????????????????????????????????????????????????????????????????????????????
?????????????????????????????????????????????????????????????????????????????????????????????????
?????????????????????????????????????????????????????????????????????????????????????????????????
?????????????????????????????????????????????????????????????????????????????????????????????????
?????????????????????????????????????????????????????????????????????????????????????????????????
???????????????????????????????????????????????????????CTATCAGCCTTTTATCCAGCCCGT
AGCCCTAGCCGTACGACTCACAGCTAATATTACTGCAGGCCATTTATTAATACACCT
AATCGGAGGAGCTGCTTTAGCCCTAACAAATATTAGTGCCCMTACTGCTTTAATTAC
CTTTATCATCC???????????????????????????????????????????????????????????TTTACCCTACTTG
TGAGCCTGTATTTACATGATAATACTTAATGACCCACCAAACCCACGCATATCACAT
GGTTAATCCCAGCCCATGGCCACTTACAGGGGCCCTTTCGGCCCTACTATTAACCTCA
GGCCTGGCTATATGATTTCACTATAACTCAATACTACTATTAACTCTAGGAATAACCA
CTAACCTATTGACTATATACCAATGGTGACGAGACATCATTCGGGAGAGCACATTCC
AAGGCCACCACACACCCATTGTTCAAAAAGGCCTCCGATACGGAATAATCCTTTT???
?????????????????????????????????????????????????????????????????????????????????????????????????
?????????????????????????????????????????????????????????????????????????????????????????????????
?????????????????????????????????????????????????????????????????????????????????????????????????
??????????????????CTATGAAACATCATTTACAATCTCGGACGGAGTTTATGGGTCCAC
CTTTTTCATAGCCACAGGGTTCCACGGACTACACGTAATTATTGGCTCTACCTTCCTA
ATCGTATGTTTCTTGCGCCAACTAAAATACCACTTCACATCGAGCCACCATTTTGGAT
TCGAAGCCGCTGCTTGATATTGACATTTCGTAGACGTGGTTTGACTGTTCTTATACGT
TTCCATTTATTGATGAGGATCCTATTCCCTTAGTATCAACAAGTACAGTTGACTTCCA
ATCAAC?????????????????????????????????????????????????????????????????????????????????????????
?????????????????????????????????????????????????????????????????????????????????????????????????
?????CTGCCCTTCTCCATAAAATTCTTCCTAGTAGCTATCACATTCTTGCTATTCGACC
TAGAAATTGCACTACTGCTCCCTCTTCCCTGAGCCTCACAAACAAACAAACTGTCAA
CCATGCTTATCACAGCCCTTCTACTAATCTCCCTATTAGCCGCAAGCCTAGCCTACGA
GTGAACCCAAAAAGGATTAGAATGAACTGAATATGATAATTAGTTT??????????????
?????????????????????????????????????????????????????????????????????????????????????????????????

?????????????????????????????????????????????????????CCTAGAAGGCATAATACTATCCCTATT
TATTATGATAACCATGGCAGTTCTAAACAATCACTTTACACTAGCTAGCATGACCCC
CATTATCCTGCTAGTATTTGCAGCCTGCGAGGCAGCACTGGGCTTGTCCCTACTAGTA
ATGGTATCAAATACATATGGTACCGACTATGTACAAAACCTAAACCTCTTGCAATGC
TAAAAATTATTATCCCCACTGCCATACTCATACCAATAACATGATTATCAAAACCCA
GCATAATCTGAATTAACTCAACCACCTATAGTTTTCTGATCAGCCTTGTTAGCCTGTC
CTACTTAAAT????????????????????????????????????????????????????????????????????????????????????
?????????????????????????????????????????????????????????????????????????????????????????????????????
?????????????????????????????????????????????????????????????????????????????????????????????????????
?????????????????????????????????????????????????????????????????????????????????????????????????????
?????????????????????????????????????????????????????AGGAACTCTGAACTTCCTAATTATTCA
ATACTGAGCCAAACCAATTTCAGCCACCTGATCTAATATCTTTCTCTGACTAGCATGC
ATAATAGCATTCATAGTAAAAATACCTCTATATGGGCTCCACCTGTGACTACCAAAA
GCACATGTCGAAGCTCCCATTGCCGGCTCAATAGTCCTTGCTGCTGTACTGTTGAAGC
TAGGAGGATATGGAATGATACGCATTACAATCCTACTCAACCCCACAAC??????????????
?????????????TGCTATCCCTATGGGGAATAATTATAACAAGCTCTATTTGTCTACGCC
AGACAGACCTAAAATCCCTAATCCCATATTCATCCGTAAGCCATATGGCCCTAGTAA
TCGTGGCCGTACTAATTCAAACACCTTGGAGTTACATAGGAGCCACAGCTCTTATAA
TCGCCCACGGACTAACTTCCTCAGTGCTATTTTGCCTTGCAAACTCAAACTACGAACG
AATCCATAGCCGAACAATAATTCTCGCACGAGGCCTACAAACCATCCTCCCCCTAAT
AGCTGCTTGATGACTACTGGCCAGCCTCGCAAACCTGGCCCTACCTCCTACTATTAAC
CTAATTGCAGAGCTATTTGTAGTAGTGGCCTCCTTTTCATGATCTAACATAACCATTA
CTCTCATGGGCACAAATATCATCATCACAGCCCTATATACCCTCTACATACTCATTAC
AACCCAACGAGGCAAATATACACACCACATTAAAAACATCAATCCATCATTCACACG
AGAAAATGCCCTAATAACACTTCATCTGCTCCCACTTTTTCTCTTATCTCTCAACCCC
AAAATCGTMCTAGGTCCTATTTACTGTAAATATAGTTTAATAAAAACATTAGATTGT
GAATC????????????????????????????????????????????????????????????????????????????????????????????
?????????????????????????????????????????????????????????????????????????????????????????????????????
??????????????????????????????????????????????????????????????????????????????ATTATATAAAGT
AACCTATACCCTCACTATGTAAAAACCACAATCTCTTACGCCTTTACCATTAGTATAA
TCCCAGCCATAATATTCATTTCCTCCGGACAAGAGATAACCATCTCAAACTGATGTT
GACTATCAATTCAAACCCTTAAATTATCACTAAGCTTCAAACTAGAT???????????????
?????????????????????????????ATGGTCGATCATAGAATTCTCAATGTGATACATACACACAG
ATCCCTATATTAACCAGTTCTTTAAGTACCTCCTTATATTCCTAATCACTATAATGAT
CTTAGTGACCGCCAATAATCTATTTCAGCTGTTTATTGGATGGGAGGGAGTAGGAAT
TATATCTTTCCTACTTATCGGATGATGATATGGTCGAGCAGACGCAAACACTGCCGC
CCTGCAAGCAATTCTCTACAACCGTATTGGAGATGTAGGATTTATCATGGCCATAGC
ATGATTCCTTACCAACCTAAATGCATGAAACCTCCAACAAATCTTTATCACTCAACAT
GAAAGCC?????????????????????????????????????????????????AGTCCGCCCAATTTGGCCTACAC
CCATGATTGCCATCAGCCATAGAAGGTCCAACTCCCGTCTCCGCCCTACTCCACTCA
AGCACAATAGTTGTAGCCGGAGTCTTCTTATTAATCCGCTTCCACCCACTCATAGAAC
AAAATAAAGCCATACAAACCCTCACTCTATGCCTGGGGGC?????????????????????????????
?????CTCACACAAAATGATATTAAAAAAATTGTTGCTTTCTCAACTTCAAGCCAATTA
GGCCTGATAATCGTTACTATCGGAATTAACCAACCCTACCTTGCATTCCTGCATATCT
GCACACACGCATTTTTTAAAGCCATATTATTCATGTGCTCCGGATCAATTATCCACAG
TCTAAACGACGAGCAAGATATTCGAAAAATAGGCGGACTATATAAACCAATACCCTT
TACTACCACCTCCCTTATTATCGGAAGCCTCGCATTAACAGGCATGCCATTCCTAACA
GGCTTTTACTCCAAAGACCTAATCATCGAGACAGCCAATACGTCGTATACCAACGCC
TGAGCCCTATTGGTCACTCTCATTGCTACATCCCTCACAGCCGCCTATAGTACTCGAA
TCATATTCTTTGCACTCCTGGGGCAACCCCGATTCAACTCCCTAAGCCCAATCAATGA
AAACAACCCCCACCTCATCAACTCCATTAAACGTCTCTTAATTGGAAGCATTTTTGCA
GGATACTTGATCTCCC?????????????????????????????????????????????????????????????????????????

????????????????????????????????????????????????????????????????????????????????????????????
????????????????????????????????????????????????????????????????????????????????????????????
????????????????????????????????????????????????????????????????????????????????????????????
????????????????????????????????????????????????????????????????????????????????????????????
???????????????????AATAAGCAAAGACCAACCAGTGACAACCACTAGCCAGGTTCYAT
AACTATACAGTGCTGCAATTCCTATGGCCTCCTCACTAAAAAACCCCGAGTCACCCG
TATCATAGATCACTCAATCACCCGCACCATTAAACTTAAACACAACCTC?????????????
???????????GCAAGCAGTCAACAACTCCGCTAATACCCCCGTAATAAACGCACCTAATA
CGGCTTTATTAGATGTCCACGCCTCGGGGTAGGGCTCAGTAGCCATAGCTGTAGTGT
ACCCAAACACCACAAGCATGCCCCCCAAATAAATTAAAAAAACTATTAAACCTAAA
AATGACCCCCCAAAATTCAATACAATACCGCAACCAACACCACCAGCCACAATCAAT
CCAAGCCCACCATAAATAGGAGAAGGCTTTGAAGAAAAACTCACAAAGCTCACCAC
GAAAATTGTACTTAAAATAAATACAATGTATGTTATCATAATTCTCACATGGATTCTA
ACCACGACCAATGATATGAAAAACCATCGTTGTATTTCAACTATAAGAACTTAATGA
CCAACATTCGAAAATCACACCCCCTTATCAAAATTATTAATCACTCATTTATTGACCT
ACCCGCCCCATCCAATATTTCAGCATGATGAAACTTTGGCTCCTTACTAGGGGTGTGC
TTAATCTTACAAATCCTCACTGGCCTCTTTCTAGCCATACACTACACATCAGACACAA
TAACCGCTTTCTCATCAGTTACCCACATTTGCCGCGACGTAAACTACGGCTGGATTAT
CCGATATCTACATGCCAACGGAGCCTCCATATTCTTTATCTGTCTATACATGCACGTA
GGACGAGGAATATACTACGGCTCCTACACCTTCTCAGAAACATGAAATATCGGGATT
GTGCTATTGTTTACGGTCATGGCTACAGCCTTCATAGGATATGTCTTACCATGAGGCC
AAATGTCTTTTTGAGGGGCAACCGTAATCACCAACCTCCTGTCAGCAATCCCATATA
TTGGGACCGACCTAGTAGAGTGAATCTGAGGGGGTTTCTCAGTAGACAAAGCTACCC
TGACACGATTCTTTGCCTTCCACTTCATCCTTCCGTTTATCGTCTCAGCCCTAGCAGC
AGTCCACCTCCTATTCCTTCACGAAACAGGATCCAATAACCCCTCAGGAATGGTGTC
CGACTCAGACAAAATCCCATTCCACCCATACTACACAATTAAAGATATCTTAGGCCT
CTTAGTACTAATCCTAACCCTCACACTACTCGTCCTATTCTCACCAGACCTATTAGGA
GACCCTGATAACTACATCCCCGCCAACCCCCTAAATACCCCTCCCCATATTAAGCCC
GAATGGTATTTCCTATTCGCATACGCAATCCTCCGATCTATTCCCAATAAACTAGGAG
GAGTTCTAGCCCTAGTCTTATCCATCTTAATCTTAGCCACTATCCCTGCCCTCCACAC
ATCCAAACAACGAGGAATAATGTTTCGACCGCTAAGCCAATGCTTATTCTGACTCTT
AGTGGCAGACCTTCTAACCCTAACATGAATTGGTGGCCAACCTGTAGAACACCCCTT
TATTGCCATCGGCCAACTAGCCTCTATCCTATACTTCTTCATCCTCCTAGTCCTAATA
CCCATCTCAGGCATTATTGAAAACCGCCTCCTTAAATGAAGAGTCTTCGTAGTATAT
AAATTACTTTGGTCTTGTAAACCAAAAAAGGAGAATATGTACTCTCCCTAAGACTTC
AAGGAAGAAGCAATAGCCCCACCATCAGCACCCAAAGCTGAAATTCTTTCTTAAACT
ATTCCTTGCCAATACCAAAAAACAACCCCATGACTTTCATAATTCATATATTGCATAT
ACCCGTACTGTGCTTGCCCAGTATGTCCTCATCCCCACAAAAAATAAGTGAAAAAAT
CCTCAATCCCCGTTAATACAGAACACACAACACGAAATAACCTGTTAACTACCGGAC
CCCCCCCCTCCCCCCGTTAACACATTACGTAGGGCATACTATGTATATCGKGCATTAA
TCGCCTGTCCCCATGAATATTAAGCATGTACAGTAGTTTATATATWTTACATAAGGC
ATACTATGTATATCGTGCATTAATCGCTTGTCCCCATGAATATTAAGCATGTACAGTA
GTTCATATATATTACATAAGACATAATAGTGCTTAATCGTGCATATTCATGATTTAGA
ACAGTTCTTTCATGGATCTCAACTATCCGAAAGAGCTTAATCACCTGGCCTCGAGAA
ACCAACAACCCTTGCTCGAGCGTGTACCTCTTCTCGCTCCGGGCCCATTTCAACGTGG
GGGTGTCTATAGTGAAACTATACCTGGCATCTGGTTCTTACTTCAGGGTCATGACATT
CTTAAATCCAATCCTTCAACTTTCTCAAATAGGACATCTCGAT

**GD/P11L: *Manis javanica***

>Manis_javanica_GDP11L_mtDNA [consensus reconstructed from 807,747 reads]

GTTAATGTAGCTTAAAACCAAAGCAAAGCATTGAAAATGCTTAGATGAGTCCACAA
ACTCCATAAACACACAGGTTTGGTCCCAGCCTTTTTATTAATTTTCAATAGGATTACA
CATGCAAGTATCCGCCCCCCAGTGAAAATGCCCTTCAAGTTATCACAGTACCTGAAG
GAGCTGGCATCAAGCACGCCAATAAACGCAGCTAACGACGCCTTGCAGAGCCACAC
CCCCACGGGAAACAGCAGTGATAAAAATTAGGCTATAAACGAAAGTTCGACCTAGC
CATATTGCATTGGGTTGGTAAATCTCGTGCCAGCCACCGCGGTCATACGATTAACCC
TAGCTAATAAAAAACCGGCGTAAAGCGTGCTAAGATAAATCCAATCCAAATAAAGT
TAAGCCCTGACCAGGCCGTAAAAAGCCGTGGTTGCCGTAAAAATAAACTACGAAAG
TAACTTTAATTCACATCGACACACGATAGCTAAGACCCAAACTGGGATTAGATACCC
CACTATGCTTAGCCCTAAACCTAAATAATTCACCAGACAAAATTATTCGCCAGAGAA
CTACTGGCAACAGCCTAAAACTCAAAGGACTTGGCGGTGCTTTACATCCCCCTAGAG
GAGCCTGTTCTGTAATCGATAAACCCCGATAAACCCTACCAATTCTAGCTAATACAG
CCTATATACCGCCATCTCCAGCAAACCCTAAAAAGGAAGCACAGTAAGCAAGACTA
TGAAAACATAAATACGTTAGGTCAAGGTGTAGCCCATGAATTGGGAAGAAATGGGC
TACATTTTCTAAAATAGAACACAAACGAACGCCCTAATGAAAACGAGGGCCAAAGG
AGGATTTAGCAGTAAGCTGAGAATAGAAAGCTCAACTGAACCCGGCCCTAAAGCAC
GCACACACCGCCCGTCACCCTCTTCAAATCCCCAAGAATACCTAAACATATTAGACA
AACACCAAGGCATGAGAAGAGACAAGTCGTAACAAGGTAAGCATACTGGAAGGTGT
GCTTGGATCACCAAAGTGTAGCTTAAACAAAGCATCTGGCCTACACCCAGAAGATCT
CAATAACATGACCACTTTGAACAAATCCTAGCCCAACCAACACCCAACATATAACCA
AACAGAGCACATAAACCAAAGCATTTACTAGACTAAAGTATAGGCGATAGAAATTA
CACAACGGCGCTATAGAAAAAGTACCGCAAGGGAAAGATGAAAGATGCATTCAAAG
TACAAAAAAGCAAAGATTACCCCCTGTACCTTTTGCATAATGGACTAACTAGAAACA
CCCTAGCAAAGAGAACTTAAGCTAGAAACCCCGAAACCAGACGAGCTACCTACGAG
CAGTTTAAAGAACCCACTCATCTATGTGGCAAAATAGTGAGAAGACTTATAGGTAGA
GGTGAAAAGCCTAACGAGCCTGGTGATAGCTGGTTGTCCAAGAAATGAATCTAAGTT
CAACTTGAAGCATACCCAAAAGCCCAAAAACTATAATGTAGGCTTCAAGTATAGTCT
AAAAAGGTACAGCTTTTTAGAAACAGAATTAAATCTTAATTAGTGAGTAAACAATAC
AACAACCATAGTTGGCCTAAAAGCAGCCATCAATTAAGAAAGCGTCAAAGCTCAAC
AATAAGACATAAATAATACCACAAATAAAAATCAACTCCTAAACCAATATTGGACC
AATCTATCAATAAATAGAAGAAATACTGTTAGTATGAGTAACAAGAAATAGATCTCC
TTGCACAAGCTTATATCAGAACGGATGACCCACTGATAATTAACAACGAAACAAATC
AAACCCAAAAATAGAACTTTGTTAAATAGATTGTTAACCCAACACAGGCGTGCATAT
TCTCAAAGGAAAGGTTAAAACAAATGAAAGGAACTCGGCAAACACAAGCCCCGCCT
GTTTACCAAAAACATCACCTCTAGCATAACCAGTATTAGAGGCACTGCCTGCCCAGT
GACTCGCGTTAAACGGCCGCGGTWTTCTGACCGTGCAAAGGTAGCATAATCACTTGT
TCTCTAAATAAGGACTAGTATGAACGGCTAGACGAGGGTTTTACTGTCTCTCATCTGT
AACCAGTGAAATTGACCTCCCCGTGAAGAGGCGGGGATAATATAATAAGACGAGAA
GACCCTATGGAGCTTTAATTAAACAGCTTAAAATTAACCCAACACCACTCTAACAGA
GACATAACAAACCAATTAGCCTAAGCTGTCAATTTTGGTTGGGGTGACCTCGGAGCA
AAAAACAACCTACGAGCGGTCAAATCCAGACTAACAAGTCCAGATAATCCGTTAATT
GATCCAACAACTTGATCAACGGAACAAGTTACCCTAGGGATAACAGCGCAATCCTAT
TCGAGAGTCCATATCGACAATAGGGTTTACGACCTCGATGTTGGATCAGGACATCCC
AATGGTGCAGAAGCTATTAATGGTTCGTTTGTTCAACGATTAAAGTCCTACGTGATCT
GAGTTCAGACCGGAGTAATCCAGGTCGGTTTCTATCTATTAATACACTTCTCCCAGTA
CGAAAGGACAAGAGAAGTAGGGCCTACCTCACACAGGCGCCCTCGAATCAATAAAT
GACTACCCTCTTAATTTAGCCAATTCACAACAACCCCAGTCCTAGAACCAGGGCCRK
YKWGGRTWGSAAAGCACGGCAACTGCACAAGACTTAAGCTCTTGTAACAGAGGTTC
AACTCCCCTCCCTAGCAGCAATGTACACAATCAACATAATAATAATGATCATCCCCA
TCCTACTAGCCGTGGCATTCCTAACGCTAGTAGAACGCAAAGTGCTAGGATACATGC
AACTACGAAAGGGCCCAAACATCGTGGGCCCTTGAGGCCTGCTACAACCAATCGCC
GACGCAGTAAAACTATTCACTAAAGAACCCCTACGACCCCTAACATCCTCAATCACA

ATATTCATCATGGCACCTATCCTAGCACTGACACTCGCACTCACCATGTGAGTACCA
CTACCAATGCCACACCCACTAGTCAACATAAACCTAGGAGTGCTATTCATACTGGCC
ATATCAAGCCTCGCCGTATACTCCATCTTATGATCAGGGTGAGCCTCAAACTCCAAA
TACGCCCTCATTGGAGCACTACGTGCAGTGGCCCAAACCATCTCATACGAAGTGACA
CTAGCAATTATCCTGCTGTCCTTACTGCTAATAAGCGGGTCCTTCACCCTCTCCACCC
TAATCACAACCCAAGAAAAGCTATGATTACTAGTGCCCGCATGACCCCTGGCCATAA
TATGATTCATCTCGACCCTAGCCGAGACAAATCGAGCACCCTTCGACCTAACAGAAG
GAGAATCTGARCTCGTATCCGGCTTTAACGTTGAATACGCAGCAGGCCCATTCGCAC
TCTTCTTCCTTGCCGAATACGCAAACATCATCATAATAAACATCTTGTCAGTRACRCT
CTTCATAGGAGCCTTTCACGACCCCCACACCCCAAGCCTCTACACAGTTAATTTTGTA
GTAAAAACACTCGCCCTTACCGCCCTATTCCTATGAATCCGAGCATCCTACCCACGA
TTCCGCTACGACCAACTAATGCACCTACTATGAAAAAACTTCCTACCACT????????????
?????????ACACGTATCAATACCAATCGCCCTGTCAAGCATTCCCCCACAATCATAAGA
AATATGTCTGACAAAAGAATTACTTTGATAGAGTAAATAATAGAGGTTTAAATCCCC
TTATTTCTAGAAAAACAGGCATTGAACCTGCACCTGAGAACTCAAAAATCTCCGTGC
TACCAACTTACACCACAATCTACAGTAAGGTCAGCTAAGCAAGCTATCGGGCCCATA
CCCCGAAAATGTTGGATCACAACCTTCCCGTACTAATAAACCTCACTATACTAGCCA
TCCTAACAACCCTATTCCTAGGGACCATGCTCGTCCTAATCAGCTCTCACTGGCTAAT
GATTTGAATTGGGTTCGAGATAAACATACTAGCAATAGTCCCTATCCTAATGAAAAA
CTTTAGCCCACGAGCCATAGAAGCAGCCACAAAATACTTCCTAATCCAAGCCACCGC
GTCCATACTACTAATGCTAGCCATTACCCTAGACCTGATATCTTCTGGACAATGAACC
ATCACAAAAACACACAACACCCTGCCATCAGCCATCATCACCGTAGCTATGGCCATA
AAACTAGGAATAGCACCATTCCACTTCTGAGTGCCAGAAGTAACACAAGGAAGCCC
ACTATCATCAGGCATGCTACTCCTAACCTGACAAAAAGTCGCACCAATATCGATCCT
ATACCAAATAATACCCACAATCAACACAAATATACTGACAACCATAGCCGCACTCTC
AATCCTCATCGGCGGTTGAGGAGGGCTAAACCAGACCCAACTACGAAAAATCATAG
CATACTCCTCAATCGCCCACATAGGCTGAATAGCAATAATCATAACATACAACCCGG
ACATTGCCATTCTAAACCTACTAGTCTACATCATAATGACACTATCTATATTCGCAAT
CCTACTATACAACTCATCAACAACAACCCTATCACTATCCCACCTATCAAACAAGAC
ACCACTAATCACAGCCCTCGCACTACTAATCCTACTATCACTAGGAGGCCTCCCCCC
ACTGACCGGATTCATGCCCAAATGAATAATCATCCAAGAACTAACCAAAAACAACA
TAGTAATAATACCAACCGTGATAGCAATAACAGCACTACTCAACCTATACTTCTACG
TACGCCTGGCGTACTCCACAGCACTAACCATACTCCCAACTACCAACAACATAAAAA
TAAAATGACAATTCGAAACAACAAAAATCATAAAACTAACCCCACCTCTAATCATTT
TATCCACAATAGCACTCCCACTCACCCCTATAATATCAATCCTAAACTAGGGGCTTA
GGCTACACAGACCGAGAGCCTTCAAAGCTCTAAGCAAATACAAACTATTTAGTCCCT
GATATAAAGACTGTAGAAATTCAACCTACATCACCTAAACGCAAATCAGGCGCTTTA
ATTAAGCTAAATCCTCACTAGATTGGTGGGATACAAACCCACGAAACTTTAGTTAAC
AGCTAAACACCCTAATCAACTGGCTTCAATCTACTTCTCCCGCC?????????????????????
???????????????????????????????????????????????????????????????????????????????????????????
???????????????????????????????????GCCATCTTACCTATGTTCACCAACCGCTGGCTCTTT
TCAACAAATCACAAAGACATTGGCACACTATATCTCTTATTTGGCGCCTGGGCCGGA
ATAGTAGGCACCGCACTAAGTCTTCTAATTCGCGCTGAACTAGGTCAGCCTGGGACC
CTCTTAGGGGACGACCAAATTTACAATGTGATCGTTACCGCACATGCATTTGTAATA
ATCTTCTTCATAGTCATGCCAATCATAATCGGGGGCTTTGGAAACTGACTAGTGCCCC
TGATAATTGGAGCCCCTGACATGGCATTCCCCCGTATAAACAACATAAGTTTCTGAC
TGCTCCCCCCCTCTTTCCTACTTCTCCTGGCCTCTTCCATAGTCGAAGCAGGGGCTGG
AACCGGCTGAACTGTATACCCCCCTTTAGCAGGAAATTTAGCACACGCAGGAGCATC
TGTAGACCTAACCATCTTCTCTCTTCATCTAGCGGGTGTCTCGTCAATCCTTGGGGCT
ATCAATTTTATTACAACAATCATCAACATAAAACCCCCTGCAATAAACCAATACCAA
ACCCCACTATTCGTGTGATCAGTCCTAATTACGGCCGTGCTTCTGCTACTATCTCTAC
CCGTACTAGCCGCTGGCATTACCATACTGTTAACCGACCGTAATCTAAATACAACCT

TTTTTGACCCCGCAGGAGGAGGTGACCCCATTCTATACCAACACCTATTCTGATTCTT
CGGACACCCCGAAGTGTACATTCTCATCCTTCCTGGGTTTGGAATAATCTCCCACATC
GTAACCTATTACTCCGGGAAAAAAGAGCCCTTTGGGTACATGGGCATAGTCTGAGCA
ATAATATCCATTGGCTTCCTGGGCTTCATTGTATGGGCACACCACATGTTCACAGTGG
GAATAGACGTTGATACACGGGCCTACTTCACATCAGCCACCATAATTATTGCTATCC
CCACTGGAGTAAAGGTGTTTAGTTGACTAGCAACCCTGCACGGAGGAAACGTAAAA
TGGGCCCCAGCTATACTATGGGCCCTGGGCTTTATCTTCCTGTTTACAGTCGGGGGTC
TAACTGGCATCGTGTTGGCCAACTCGTCCCTAGATATCGTCCTCCATGATACCTACTA
CGTAGTAGCCCACTTCCACTACGTTCTCTCCATGGGAGCAGTTTTCGCCATCATAGGA
GGCTTCGTCCATTGATTCCCCCTGTTCTCAGGATACACGCTCAACAACACATGGGCA
AAAGTTCACTTTACAATCATATTCGTGGGCGTAAACATGACCTTTTTCCCCCAACACT
TCCTTGGACTGTCTGGAATACCTCGACGATACTCGGACTACCCAGACGCTTACACAA
TGTGAAACACTGTGTCCTCCATAGGATCCTTCATCTCATTAACCGCCGTAATACTCAT
GGCCTTCATAATCTGAGAGGCATTCGCCTCTAAACGGGAAGTCCTAATAGTGGAGTC
CACAAATACCAACCTCGAATGACTACACGGCTGCCCACCACCCTACCACACATTCGA
AGAACCTGCCTTCGTAAATCTGGTCAAAACAAGAGAGGAAGGAATCGAACCCTCAA
ACAATGGTTTCAAGCCAATATCATAACCCCTATGTCTCTCTCAATCACAGAAGTATTA
GTAAAATTTACATAACTTTGTCAAAGTTAAATTATAGGTTCAAGCCCTTTATACTTCC
ATGGCGTACCCGCTCCAATTAGGCTTTCAAGATGCCACATCCCCAATCATAGAAGAA
TTACTCCACTTCCACGATCACACACTAATAATCGTGTTCTTGATCAGCTCCCTAGTAC
TATACATTATCTCTCTCATACTAACAACCAAACTCACCCACACAAGTACAATAGACG
CTCAAGAAGTAGAAACCATTTGAACCATCCTCCCCGCAATTATCCTAATCCTAATCG
CCCTGCCCTCCCTACGCATCCTCTACATAATGGACGAGATTAACAACCCCGCCCTGA
CAGTAAAAACAATGGGCCATCAATGATACTGAAGCTACGAATACACAGACTACGAA
GACCTAAGCTTCGACTCATACATAATCCCAACTCAAGACCTAAAACCCGGCGAACTC
CGACTTCTAGAAGTGGACAACCGACTTGTACTACCCATAGACACAACCATCCGCATG
CTCATCTCATCTGAAGACGTCCTGCACTCTTGAGCCATCCCATCCCTGGGCCTAAAAA
CAGATGCTATCCCGGGGCGTCTAAACCAAACAACCCTGATATCAACCCGGCCCGGCC
TATTCTACGGACAGTGCTCAGAAATCTGTGGCTCAAATCACAGCTTCATGCCAATTG
TCCTCGAACTAGTACCCCTAAAAACATTTGAAAACTGAACTACGTCCCTACTGTAAT
TCATTGAGAAGCTAATAGCGCTAGCCTTTTAAGTTAGAGATTGAGAGCACGCCTCTC
CTCAATGACATGCCTCAACTAGACACTACAACATGA????????????????????????????????
?????????????????????????????????????????????????????????????????????????????????????????????????
?????????????AATGAACGAAAATCTATTCACCTCTTTCATTACCCCAGTAATAATAGGG
ATCCCTATTGTAACAATTATCATTATGTTCCCGGTAATCCTCTTCCCAACATCAAACC
GACTAATCAACAACCGCATTGTATCCATACAACAATGACTCCTAAAACAAACATCCA
AACAAATAATAAGCATCCACAACTACAAAGGACAGACCTGAACCCTGATATTAATA
ACACTAATCATTTTCATCGCATCTACTAACCTACTAGGCCTGCTACCCCACTCATTCA
CCCCCACAGCCCAACTGTCAATAAACCTGAGCATAGCCGTTCCTCTATGGGCAGCCA
CCGTAGTCACAGGTTTTCGACACAATACAAAAACATCTTTAGCCCACTTCCTACCCC
AGGGAACACCAACCCCCCTCATCCCAGTGCTAGTGATCATTGAAACAATCAGCCTGC
TAATCCAACCCATAGCGCTCGCAGTACGACTGACAGCCAACATCACCGCCGGCCACC
TGTTAATACACCTAATCGGAAGCGCAACCCTTGCCCTAATATCAATCAACCTCACCG
TGGCCACAACTACCTTCATCGTCCTAGTCCTGCTTACAATCCTTGAATTCGCAGTCGC
ACTCATTCAGGCCTACGTCTTCACTCTTCTAATCAGCCTCTACTTACATGATAACACA
TAATGACCCACCAAACCCACTCATACCACATAGTAAACCCAAGCCCTTGACCCCTAA
CCGGGGCCCTATCTGCCCTCCTAATAACATCGGGCCTAGCAATATGATTCCACTTCA
ACTCTACAATACTACTCCTCCTAGGACTAACAACAAACCTCTTAACAATATATCAAT
GATGACGTGACATTGTACGAGAAAGCACCTTTCAAGGCCACCACACACCCACAGTCC
AAAAAGGACTACGATACGGCATAATCCTATTCATTGTCTCAGAAGTATTTTTCTTCGC
TGGCTTCTTCTGAGCATTCTACCACTCAAGCCTAGCACCTACCCCCGAACTAGGAGG
ATGCTGACCCCCCACAGGCATCAACCCGCTAAACCCACTGGAAGTACCTCTACTCAA

CACATCCGTTCTTTAGCCTCAGGAGTATCAATTACATGGGCACACCATAGCTTAATA
GAAGGCAGCCGAAACCACATAACCCAGGCCCTGCTCATCACAATCCTCCTAGGCATT
TACTTCACACTACTGCAGGTCTCAGAATATTACGAAGCACCCTTCACAATCTCCGAC
GGCGTGTACGGCTCTACCTTCTTTGTGGCAACTGGGTTCCACGGCCTACACGTCATCA
TCGGCACCTCCTTCCTAACTGTGTGCCTACTACGACAACTAAAATACCACTTTACATC
AAACCACCACTTCGGATTCGAAGCTGCCGCCTGATACTGACACTTCGTAGATGTAGT
GTGACTGTTCCTGTATGTCTCCATTTACTGATGAGGTTCCTATTTTCTAAGTATGCAC
AGTACAGTTGACTTCCAATCAACAAGCTCTGGCCACAACCCAGAAGAAAATAATAA
ACCTACTCCTAGCAATAACAACAAACACCATACTAGCATGCCTGCTCATACTAATCG
CCTTCTGACTCCCTCAATTAAACACGTACTCGGAAAAAATCACCCCCTACGAATGCG
GATTCGACCCCATGGGATCGGCACGACTACCATTCTCCATAAAATTCTTCCTAATCGC
CATCACGTTCCTACTATTCGACCTAGAAATCGCACTACTCCTGCCACTCCCCTGAGCA
TCACAAACAAACAACCTAGGCACCATAATCACCGTAGCCTTGGTCCTCATCCTACTG
CTAGCAATCAGCCTGACCTACGAATGAACACAAAAAGGCCTAGAATGAACTGAGTA
TGGTAGCTAGTTTACACAAAACAAATGATTTCGACTCATTAAACTATGACTTACTCAT
AGCTACCAAATGTCCCTAATTTACATTAACACAACACTGGCATTCACTATCTCCCTAA
TGGGGATATTAATGTACCGATCACACCTAATGTCCTCACTACTATGTCTAGAGGGCA
TAATACTCTCCCTATTCGTAATAATCACCATTACAATCCTAACTAACCACTTCACACT
AGCCAACATAGCACCCATCATCCTCCTCGTACTAGCAGCATGCGAGGCTGCCCTAGG
CCTCTCTCTACTAGTACTTGTATCTAACACATACGGCACGGACCACGTACAAAACCT
AAACCTGCTACAATGCTAAAACTAATTGTTCCCACAGCCATACTAATCCCCCTAACA
TGACTATCGAACAAAAACATAGTGTGAATCAACGTAACATCACAC?????????????????
?????????GGCATCTTGGGCCAGCATGACCACAACAACATAAGCCTCTCAGCCATCTTC
TTCTCTGACCCCMTATCAGCACCCTTAATCGTATTAACAACATGACTCCTCCCCCTAA
TACTAACAGCCAGCCAAGCCCACCTCTCCAATGAGCCCCTGGGCCTAAAAAAACTAT
ACATTACCATACTAATCACCCTCCAAGCACTACTAATCATAACATTCGCCTCCTCTGA
GTTCATTATATTCTACATCCTGTTTGAAGCAACACTCGTACCAACCCTAATCATCATC
ACACGATGAGGAAACCAAGCAGAGCGACTAAACGCAGGGTCCTACTTCCTCTTCTAC
ACAATAGTAGGGTCACTCCCCCTTCTAGTAGTACTCACATACACCCAAAACATAACA
GGAACCCTAAACATACTAGTACTACAATACTGAGCGAAACCCATAAACGACTCTTGA
TCCAACATGCTAATATGACTAGCATGCATAATAGCATTCATAGTAAAAATACCCCTA
TACGGACTACACCTCTGACTGCCAAAAGCCCACGTAGAGGCCCCCATTGCAGGCTCC
ATAGTACTTGCAGCCGTACTACTAAAACTAGGCGGATACGGCATAATACGCATCACT
ATCATGCTAGAACCAATAACAACATTCATAGCATACCCCTTCCTAATACTATCCCTGT
GAGGAATAATCATAACAAGCTCCATCTGCCTTCGCCAAACCGACCTAAAATCACTAA
TCGCCTACTCCTCCGTAAGCCACATAGCACTGGTAATCGTCGCAATCCTAATCCAAA
CCCCATGAAGCTACATAGGGGCCACCGCTCTCATAATCGCCCACGGTCTAACATCCT
CCATACTATTTTGCCTAGCAAACACAAACTATGAGCGAACCCACAGCCGAACTATAA
TACTAGCACGAGGACTGCAAACCCTCCTACCCCTAATAGCTGCCTGATGGCTCCTAG
CAAGCCTAACCAACCTGGCCCTCCCCCCCAGCATCAACCTAATCGGAGAACTATTCG
TAGTAGTATCAACATTCTCGTGGTCTAACACCACCATCATCCTTACTGGAACAAACA
TTATCATCACAGCCGCCTACTCCCTATACATACTAATATCCACCCAACGTGGAAAAT
ACACCCACCACATCAACAACATCTGCCCCTCCTTCACCCGAGAAAACGCTCTAATGG
CACTCCACATACTGCCCCTACTAATGCTATCAACCAACCCCAAAATCATCCTAGGGT
GCCTTTACTGTAAGTATAGTTTAATAAAACCTCAGATTGTGGATCTGACAATAGAAG
ACCACAACTTCTTACTTACCAAAAAAGCATGCAAGAACTGCTAATTCCTGCCCCCAT
GTATAAAAACATGGCTTTTTTTAAACTTTTAGAGGATGACAGACACCCGTTGGTCTTA
GGAACCAAAAGACTTGGTGCAACTCCAAGTAAAAGTAATTAACACCCTAACCTGCA
CCTCCCTTATAACACTAACAACACTAAYCATACCAATCATAATCACCCCCACAAATG
CCTACACAAGCAAAAACTACC????????????????????????????????????????????????????????
???????????????????????????????????????????????????????????????????????????????????
??????????????????????????????????????????????????????????????????????????????????

TCAGACCCCCACATCAACCGATTCTTCAAGTACCTACTAATCTTCCTAATCACAATAA
TAATCCTAGTATCCGCCAACAACATATTCCAACTCTTCATCGGCTGAGAAGGAGTAG
GAATCATATCTTTCCTCCTAATCGGCTGATGACACGGACGAACAGATGCAAACACAG
CAGCCATACAGGCCATCCTATACAACCGCATCGGAGACATTGGACTAATCCTGTCAA
TGGCATGATTCCTCACAAACCTAAACTCATGGGACCTCCAACAAATCCTCATGCTAA
ACCCCGAGAACACAAACATCCCCCTAGCGGGCCTACTACTAGCAGCAACTGGAAAA
TCCGCACAATTTGGGCTACACCCATGACTCCCCTCAGCCATAGAAGGCCCAACCCCA
GTATCAGCCCTACTCCACTCCAGCACAATAGTAGTCGCAGGGGTATTCCTACTAATC
CGATTCCACCCCCTAATAGAAAACAACAAAACAATCCAAACAACAACCCTATGCCTA
GGGGCCATTACAACCCTGTTCACAGCCATCTGTGCTCTCACCCAAAACGACATCAAA
AAGATCGTGGCCTTCTCAACCTCAAGCCAACTAGGTCTAATAATAGTCACCATCGGA
ATCAACCAACCACACCTAGCCTTCATACACATCTGCACCCACGCCTTCTTCAAAGCC
ATACTATTCATGTGCTCCGGATCGATCATCCACAGCCTAAACGACGAACAAGACATC
CGCAAAATAGGAGGACTTTTCCACGCCCTCCCCACTACCACCTCCGCTCTCATTATTG
GCAGCCTAGCCCTAACAGGAACCCCATTCCTAACAGGCTTCTACTCTAAAGACCTAA
TCATCGAAACCGCTAACATATCCCACACAAACGCCTGAGCCCTGCTCATCACACTAG
TAGCCACCTCCCTCACAGCCGCCTACAGCACACGAATCATCTTCTTCACACTATTAGG
ACAACCCCGATCCAACACCCTAATCAACATCAACGAAAACAACCCCTCACTAACAA
ACCCAATCAAACGCCTAATGCTAGGGAGCATCTTCGCCGGATTCCTACTATACAACA
ACATMCCCCCAACAACCATGCCCCAAAC??????????????????????????????AC
AATCCTAGGCCTAATCATAGCCCTAGAACTGTGCAACCTGTCCCTAAACCTAAGCCA
CAAACCCCCCTCAAGTGCCTTCAAATTCTCAACCTTACTAGGCTACTTCCCCACAATC
ATTCACCGCTCAGGACCCCTCTGATCACTAACAACAAGCCAAAAACTAGCATCCCTT
ATTCTAGACCTAACCTGACTAGAAAATATTATACCAAAGTCAATCTCACACTTCCAC
ATAAAAGCCTCCCTAATAACATCCAACCAAAAAGGCTCAATCAAACTATACTTCCTG
TCCTTCGCAGTAACAATAATCCTCGCTCTCCTCATATTCTATTCCCCCGGGTAACCTC
CATAACAACAACAACACAAATAAACAAAGACCACCCGGTCATAATCACAAACCAAA
CACCATGACTATAAAGAGCAGCAACACCCGCAGCTTCCTCACTAAAAAAACTAGAC
CCCCCAACATCATAAGTCGCCCAATCACCAGGACCACCAAGATCAAAAACCACACTC
AACCCCTCACCCTTCAACATATAAAGCACTAAAACCAACTCTATGACCACACCCAAA
ACAAAAGCACCGAGCACAACCACATTAGACACCCAAACTTCAGGATACTCCTCAAT
GGCTATGGCCGCCGTATAACCAAAAACAACCAACATCCCCCCAAGATAAATCAAAA
AAACCATTAAACCCAAAAAAGACCCACCAAAATTAACTACAACCCCACAACCTGCA
CCGCCAGCTACAACCAGACTAAACCCCCCATAAATAGGAGAAGGCTTTGAAGAAAA
CCCTAAAAAACTAATCACAAAAATAACACTCAAAATAAACACAACATATGTCATTRT
TCCCACATGGACTCAAACCATGACCTATGACACGAAAAATCATCGTTGTAATTAAAC
TACGGAAACCTAATGACAAACATCCGAAAAACACACCCCCTACTCAAAATCATCAA
CAACTCCTTTATTGATCTCCCCACCCCCTCTAACATCTCAGCATGATGAAACTTCGGA
TCACTGCTAGGAATCTGCTTAGTCCTACAAATTCTCACAGGACTATTTCTAGCCATAC
ACTACACAGCAGACACAGCAACCGCATTCTCATCAGTAACTCACATCTGCCGAGATG
TAAACTACGGATGAATTATCCGATACATACACGCTAACGGAGCTTCCCTGTTCTTCAT
CTGCCTGTTCGCACACATCGGACGAGGCATCTACTACGGATCCTTTGCCTACAAAGA
GACATGAAACATCGGTATCCTGCTCCTGTTTGCAGTAATAGCAACAGCCTTTATGGG
ATACGTCCTACCATGAGGACAAATGTCCTTCTGAGGTGCTACAGTAATTACAAACCT
TTTATCCGCAATACCCTACATCGGGTCTAGCCTAGTAGAGTGAGTCTGAGGGGGATT
CTCGGTAGACAAAGCAACTCTCACTCGATTCTTCGCTCTTCACTTCATCCTTCCCTTC
GTAATTCTTGCCCTAGTACTAGTACACTTACTATTCTTACACGAAACCGGATCCAACA
ACCCAATAGGAATCGTATCCAACCCCGACGTAATCCCCTTCCACCCATACTACACAG
CCAAGGACACCCTTGGCCTATTCATCATGCTCACAGCACTAATATCCTTAGCCCTATT
CTTCCCCGACCTACTAGGAGACCCAGACAATTATACA???????????????????????
???????????????????????????CCTACGCTCAATTCCCAACAAATTGGGAGGAGT
ACTAGCACTAATCCTATCCATCCTCGTACTAGCACTAATCCCACTACTACACACATCA

AAACAACGAACTATAATGTTCCGACCCCTGAGCCAAACCATCTTCTGACTCCTGGTG
GCCGACCTACTAGTTCTCACATGAATCGGAGGACAACCCGTGGAACACCCCTTCATC
CTAATCGGACAAGTGGCCTCTATCCTTTATTTCACACTAATCCTAGCAGCAATGCCAA
TCGCAGGTATCATCGAAAACAATCTCATAAAATGAAGAGTCTTTGTAGTATATGTCA
ATACACTGGTCTTGTAAACCAGCAAAGGAATTAACCCTCCCCAAGACTCAGGAAGA
AGACAAAAGCCCTACCATCAGCACCCAAAGCTGAAATTCTCAATAAACTACTTCCTG
CAAACAATTAAAGTCCAACAARCTTTAATTGTTTGCAGGAAGTAGTTTATTGAGAAW
WTSRKYTGCCCCATGCATATAAGCAGGTACATTATATTATTATAGTACATAGGACAT
ATTATGTATAATCGTGCATTATTGATCTAGTACATACATATAAGCAGGTACATTATAT
TATTATAGTACATAGGACATATTATGTATAATCGTGCATTATTGATCTAGTACATGCA
TATAAGCAGGTACATAATATCCTTAGAGTACATAGTACATATTATTATTGATCGGAC
ATAGCACATCAAGTCAAATCATTTCCAGTCAACATGCGTATCCCTACCACTGAAGGC
CGCCTAATCACCATGCCGCGTGAAATCATCAACCCGCTCATAGTCGTGTCCCTCTTCT
CGCTCCGGGCCCATACGGACTGTGGGGTAGTTAAAGTGAAACTATACCTGGCATCTG
GTTCTTACTTCATGTTCATCTTATATCTAGCCGCTCACTCGTTCCTCTTAAATAAGACA
TCTCGATGGATTAATTACTAATCAGCCCATGCCTAACATAACTGTGCTGTCATGCCTT
TGGTATCTTTTAATTTTCGGGGTGCGAGGTTCAACTGGGTCACCGACCTTCGAGCAG
GTGGATAACTTGTAGATAGACATTCATTGAATATTATTGGTCGTACATACTAACTCCT
AGGTGTTATTCAGTCAATGGTTACAGGACATAGAGAATTTTACAACAAAATTTTGAG
T??????????????????????????????????????????????????????????????????????????????????????
????????????????????????????????????????????????????????????????????????????????????????
????????????????????????????????????????AAAACTACTAGCGCAGCTTTCTGTCCACTAG
TTAAATACTTAGGTGCTTGAACATATATGGCTGGAAGGCCATGTAGTGACTACGTAA
TGATTTCTATTAGCCACAATAAAAACAATATACAATAAATWYAAMTWTARWTCAAT
TCTTATAGGCGCCGATAGCATAAACTATCTGCCCCCTATGTACAATTGAATTAAGGTT
AATGTAGCTTAAAACCAAAGCAAAGCATTGAAAATGCTTAGATGAGTCCACAAACT
CCATAAACACACAGGTTTGGTCCCAGCCTTTTTATTAATTTTCAATAGGATTACACAT
GCAAGTATCCGCCCCCCAGTGAAAATGCCCTTCAG
